# TIME-CoExpress: Temporal Trajectory Modeling of Dynamic Gene Co-expression Patterns Using Single-Cell Transcriptomics Data

**DOI:** 10.1101/2025.01.23.634392

**Authors:** Shuyi Yang, Anderson Bussing, Giampiero Marra, Michelle L. Brinkmeier, Sally A. Camper, Shannon W. Davis, Yen-Yi Ho

## Abstract

The rapid advancements of single-cell RNA sequencing (scRNAseq) technology provide high-resolution views of transcriptomic activity within a single cell. Most routine analyses of scRNAseq data focus on individual genes; however, the one-gene-at-a-time analysis is likely to miss meaningful genetic interactions. Gene co-expression analysis addresses this issue by identifying coordinated gene expression changes in response to cellular conditions, such as developmental or temporal trajectory. Identifying differential co-expression gene combinations along the cell temporal trajectory using scRNAseq data can provide deeper insight into the biological processes. Existing approaches for gene co-expression analysis assume a restrictive linear change of gene co-expression. In this paper, we propose a copula-based approach with proper data-driven smoothing functions to model non-linear gene co-expression changes along cellular temporal trajectories. Our proposed approach provides flexibility to incorporate characteristics such as over-dispersion and zero-inflation rate observed in scRNAseq data into the modeling framework. We conducted a series of simulation analyses to evaluate the performance of the proposed algorithm. We demonstrate the implementation of the proposed algorithm using a scRNAseq dataset and identify differential co-expression gene pairs along cell temporal trajectory in pituitary embryonic development comparing *Nxn*^−*/*−^ mutated versus wild-type mice.

## 1 Introduction

Most existing tools for studying transcriptional activities using scRNAseq data focus on individual genes. While informative, this one-gene-at-a-time approach ignores the interactions between genes. Arguably, changes in genetic interactions, sometimes without alteration in mean expression levels, have been reported to play a more critical role in downstream cellular transitions (1). Gene co-expression analysis addresses the problem by identifying coordinated gene expression changes in response to the biological conditions in question. Current approaches for gene co-expression analyses, such as WGCNA (2), graphical LASSO (3), MIST (4), and CellTrek (5) for detecting genetic interactions assume static gene correlations and do not seek to determine the dynamics of changes in genetic interactions in living cells in one unified modeling framework.

In the cells, two or more genes can have coordinated expressions that vary dynamically as cells progress through the transitional states. For example, genes associated with pluripotency might show high co-expression early in embryogenesis, and the co-expression patterns of these genes might change over time as cells differentiate into specific lineages (such as ectoderm, mesoderm, or endoderm). During immune cell activation, co-expression patterns of genes related to immune activation may change along the temporal trajectory as cells progress from an inactive state to an activated state. In neuronal differentiation, genes related to neural progenitor identity might exhibit high co-expression initially, with these patterns altering as cells differentiate into distinct neuronal subtypes. Pseudotime is frequently employed in scRNAseq analysis to identify cells in a population at various transitional states. Capturing the dynamic co-expression patterns along cell temporal trajectory is critical for understanding how a cell’s transcriptomic activities change across developmental time.

Pseudotime inference was first proposed by Trapnell et al. (6) to represent the cell order along lineages by sequencing cells according to the gradual changes in gene expression rather than measuring real time. Since then many studies have validated that pseudotime inference is a reliable method by comparing cell pseudotime to known biological timelines such as TSCAN (7), Slingshot (8), and Monocle (6). Another validation approach involves using experimental techniques like lineage tracing. In this paper, we applied Slingshot (8) pseudotime analysis to scRNA-seq data to reconstruct cellular temporal trajectories. This method is chosen based on the characteristics of our dataset. Slingshot is an unsupervised method that does not require predefined clusters. It is robust to noise in the data by estimating smoothed trajectories and allows for multiple lineages. Other methods can be also easily adapted into our proposed analysis framework.

Several statistical methods have emerged to explore gene expression alterations along cellular temporal trajectories. Most existing analyses of temporal trajectories focus on one gene at a time. Van den Berge et al. (9) introduced a generalized additive model framework based on the negative binomial distribution called tradeSeq, which can model the non-linear changes in gene expression along cell pseudotime using zero-inflated data. However, this approach of examining genes individually may overlook crucial genetic interactions. Trapnell et al. (6) applied Monocle to trace the trajectory of single-cell gene expression over time, but it also ignores gene-gene interactions.

In biological system, gene expression patterns could exhibit on-off behavior. Some genes may turn on and then turn off over cell time, while others transit from inactive to active states. This biological on-off phenomenon can be observed through zero-inflation proportions of genes in scRNAseq data (10). The zero-inflation rates of genes often change as cells progress along pseudotime. To appropriately model these covariate-dependent zero-inflation rates, it is important to incorporate time-dependent zero-inflation proportions in the analysis. However, current research on scRNAseq data often fails to capture the dynamic changes of gene zero-inflation along the cellular temporal trajectories.

A few methods have been introduced in the literature to study gene co-expression patterns. Yang and Ho (11) proposed the zero-inflated negative binomial dynamic correlation model (ZENCO), which incorporates zero-inflation for scRNAseq data but assumes a parametric linear function between gene co-expression and the covariates. However, in molecular biology, gene co-expression may relate to cellular time in a non-linear semiparametric or nonparametric manner. Lau et al. (12) employed a network approach that inferred 10 discrete co-expression networks by smoothly sliding along from early to late development using five consecutive time points. However, they only used discrete gene expression networks, while gene co-expression can change continuously. Hans et al. (13) proposed a boosting-based framework to capture complex dependence structures between outcome variables that can be applied to analyze gene co-expression; however, this method could not handle zero values in the expression measurements. Given that scRNA-seq data is typically count-based and zero-inflation rates over temporal trajectory could be of scientific interest, the boosting method may fail to adequately capture the changes of zero-inflation rates over time in the data. scHOT (1) uses Spearman’s correlation for capturing genes’ interactions along cell developmental time; however, it does not model changes in zero-inflation rates along pseudotime or developmental trajectories. In addition, scHOT could not analyze and compare data with multi-groups (wild-type versus mutant, for example) in the same modeling framework. SCODE (14) uses ordinary differential equations (ODEs) to model the temporal dynamics of gene expression and infer interactions between genes; however, it is a strictly linear model and cannot capture the non-linear co-expression changes over pseudotime. Therefore, recognizing non-linear combinations of co-expressed genes across various pseudotime domains is crucial for advancing our understanding of gene interactions.

We propose a flexible copula-based framework, TIME-CoExpress, to model and predict non-linear gene pair co-expression changes along cell pseudotime. One of the unique features of this framework is its ability to accommodate covariate-dependent zero-inflation and correlation changes. It is designed to model the dynamic gene zero-inflation patterns throughout cellular temporal trajectories to capture the biological on-off characteristics of gene expression. The copula model enables us to construct a joint model with flexible marginal distributions. TIME-CoExpress can capture the non-linear dependency structure of gene pair co-expression and explore how predictor variables, such as cell pseudotime, influence the interactions between genes.

An advantage of TIME-CoExpress is that it allows for multi-group analysis, whereas many existing methods, such as scHOT (1), are limited to analyzing each group separately, which can lead to low efficiencies. The multi-group analysis enables the simultaneous examination of different groups of data, such as mutant or wild-type groups, offering a direct comparison of gene co-expression patterns and changes in zero-inflation rates across cellular pseudotime in a unified analytical framework.

To model the correlation structure in a semiparametric manner, we extend generalized additive models for location, scale, and shape (GAMLSS) (15) to include zero-inflation and construct an additive distributional regression framework. The proposed framework allows us to model multiple parameters of a distribution function, rather than just one parameter as in the traditional GAM models. Within distributional copula regression, each response distribution parameter linked to the covariate is modeled through additive predictors. The model is fitted using splines, which accommodate non-linear changes in dependence structures along temporal trajectories derived from scRNAseq data. A trust region method (16) for simultaneously estimating the predictor effects is employed in the proposed framework.

Through a series of simulation analyses, we verified that the proposed framework can capture the non-linear relationships between cell pseudotime and the interactions of gene pairs. It can also provide dynamic gene zero-inflation patterns across cell pseudotime. Our proposed analytical framework achieves higher power compared to the Gaussian-model-based model Liquid Association proposed by Li (17), the non-model-based approach scHOT (1), and the Conditional Normal Model (CNM) (18). Additionally, our model has a significant advantage in terms of computational time compared to these three methods.

We applied the proposed algorithm to a mouse pituitary gland embryological development scRNAseq dataset. Specifically, we identified differentially co-expressed gene pairs along cellular temporal trajectories between *Nxn*^−*/*−^ mice and the wild-type group. We identified several genes with zero-inflation patterns that align with known biological processes.

This paper is structured as follows. The ‘Materials and methods’ section describes the dataset used in this analysis, a brief theoretical overview of the proposed framework, and a series of simulation studies to validate the framework. This section also includes a power analysis to compare TIME-CoExpress with current research. The results of real data analysis using mice *Nxn* pituitary gland embryological scRNAseq data are included in the ‘Results’ section. Finally, a thorough discussion and conclusion are provided in the ‘Discussion’ section.

## 2 Materials and methods

### 2.1 Experimental dataset

The scRNAseq dataset we used in this analysis is derived from dissected tissue containing part of the hypothalamus and the whole pituitary gland from embryonic day of development day 14.5 mouse embryos (e14.5). Two groups are represented in the data set; the control group contains tissue from one *Nxn*^+*/*+^ and one *Nxn*^+*/*−^ embryo, and the mutant group contains tissue from two *Nxn*^−*/*−^ embryos. *Nxn*^+*/*−^ mice are phenotypically normal. At total of 19,625 cells were sequenced, and the mean reads per cell was 27,387. Sequencing reads were processed, and cells were clustered using Seurat 4.1.0. Clusters were identified using marker genes, and clusters for hypothalamic cells, pituitary posterior lobe cells, blood cells, vasculature cells, and mesenchymal cells were removed from the analysis. The data set was further refined to only include anterior lobe pituitary cells that represent the *Sox2* stem cells, the *Prop1* progenitor cells, and the *Pou1f1* progenitor cells. Experimental data demonstrates that *Nxn*^−*/*−^ embryos have a defect that alters the differentiation of pituitary stem cells into progenitors cells and eventually hormone producing cells. We narrowed our data set to focus on this critical transition and employ TIME-CoExpress to identify genes pairs with differential co-expression patterns between the two groups during this critical developmental window. See Martinez-Mayer et al. (19) for a more detailed description of the data collection procedure.

The transcription factors: *Sox2, Prop1*, and *Pou1f1* are essential for cellular differentiation and the proper development of the pituitary gland. Yoshida et al. (20) demonstrated that *Prop1* coexists in *Sox2*-expressing cells. Olson et al. (21) reported that *Prop1* activates *Pou1f1* expression. The embryological developmental pathway for the pituitary gland follows the progression from *Sox2* progenitor to *Prop1* progenitor and then to *Pou1f1* progenitor, as progenitor cells differentiate into hormone secreting cell types (22, 23). In the following analysis, we mainly focus on *Sox2* progenitor with 766 cells, *Prop1* progenitor with 1,249 cells, and *Pou1f1* progenitor with 1,813 cells in this study. There are 2,138 cells from the wild-type group and 1,690 cells from the mutant group.

Figure 1 shows the cell trajectory estimated from Slinghot (8). The shades of red color points are the mutant group, and the shades of blue color are the wild-type group. The triangles, circles, and squares represent *Sox2* stem cells, *Prop1* progenitor cells, and *Pou1f1* progenitor cells respectively. We observe there are more cells in the early cell stage (*Sox2* stem cells) and much fewer cells in the late stage (*Pou1f1* progenitor) in the mutant group compared to the wild-type group. This indicates that the cells without the *Nxn* gene are unable to differentiate successfully and stall in an earlier stage of development. These findings are consistent with a known role for the WNT signaling component CTNNB1 (beta-catenin) interacting with *Prop1* to promote differentiation into *Pou1f1* progenitors and subsequent hormone cell types (21, 24). NXN regulates the availability of Dishevelled (DVL) (25); therefore, the stalled differentiation seen in *Nxn*^−*/*−^ embryos is consistent with a defect in WNT mediated pituitary cell differentiation. The pseudotime analysis shows that more mutant *Nxn*^−*/*−^ cells are in the *Sox2* stem cell and *Prop1* progenitor cell stage compared to the wild-type cells, suggesting that *Nxn* promotes the differentiation of pituitary stem cells. A similar pseudotime analysis and phenotypic description of *Nxn*^−*/*−^ embryos is in preparation (MLB, SAC, and SWD personal communication).

**Fig. 1.**
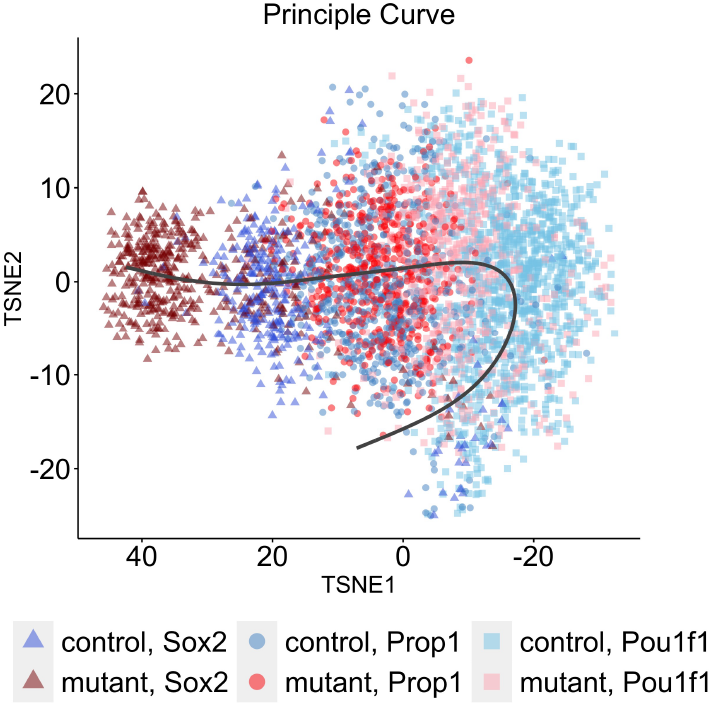
Cell Temporal Trajectory: The shades of red color points represent the mutant group, the shades of blue color represent the wild-type group; triangles, circles, and squares represent *Sox2* stem cells, *Prop1* progenitor cells, and *Pou1f1* progenitor cells respectively. Each point represents one cell, the gray curve is the principal curve. The cell pseudotime increases along the principal curve (from left to right).

### 2.2 Methods

In this section, we first introduce our zero-inflated copula model. Then we illustrate the form of additive predictors and the penalized likelihood for parameter estimation. The model construction is discussed last.

#### 2.2.1 Zero-inflated copula model

Let ***y***_***i***_ = (*y*_*i*1_, *y*_*i*2_)^⊤^denotes the gene expression measurements of a gene pair in cell *i, i* = 1, …, *N*; *N* is the total number of cells. We assume the two genes have zero inflation rates *p*_*i*1_ and *p*_*i*2_ respectively. The TIME-CoExpress implements a two-part model for the zero and non-zero expression values. For the non-zero expression values, denoted by *w*_*i*1_ and *w*_*i*2_, we consider Gamma (GA) distributions with parameters ***ϑ***_***i*1**_ = (*µ*_*i*1_, *σ*_*i*1_)^⊤^and ***ϑ***_***i*2**_ = (*µ*_*i*2_, *σ*_*i*2_)^⊤^respectively. We adopt the parametrization according to Rigby et al. (26). The probability density function (PDF) and cumulative distribution function (CDF) are given by,

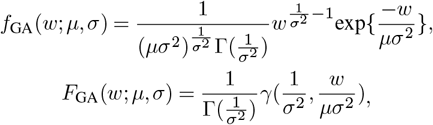

where *w* > 0, *µ* > 0 and *σ* > 0. We consider Gamma distribution as the marginals because the gene expressions in mouse pituitary gland scRNAseq data are normalized continuous measurements. In addition, the Gamma distribution can account for the over-dispersion characteristic inherited inside scRNAseq data. Other marginal distributions such as negative binomial (described in Supplementary Data—Appendix A1) can also be implemented in the proposed framework.

The joint distribution of non-zero values of the gene pair ***y***_***i***_ is then modeled by the Gaussian copula model. The Gaussian copula is widely used in modeling genomic data because of its flexibility and interpretability. The complex dependencies often encountered in genomic data can be modeled using a Gaussian copula, with the dependence parameter specified through splines. Additionally, its simple and intuitive mathematical structure, similar to the multivariate normal distribution, makes it easy to interpret when it comes to constructing joint distributions for different types of outcomes (27). The PDF is given by,

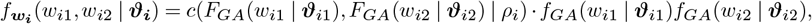

where ***ϑ***_***i***_ = (*µ*_*i*1_, *σ*_*i*1_, *µ*_*i*2_, *σ*_*i*2_, *ρ*_*i*_)^⊤^is the vector of all distributional parameters for cell *i*; *ρ*_*i*_ is the dependence parameter that models the correlation between *w*_*i*1_ and *w*_*i*2_. *F*_*GA*_ and *f*_*GA*_ represent CDF and PDF of the Gamma marginal distribution, respectively. *c* (·, · | *ρ*_*i*_) is a Gaussian copula density.

For simplicity, we denote *F*_*GA*_(*w*_*i*1_ | ***ϑ***_*i*1_) and *F*_*GA*_(*w*_*i*2_ | ***ϑ***_*i*2_) by *u*_*i*1_ and *u*_*i*2_, respectively. The Gaussian copula density *c* (·, · | *ρ*_*i*_) (28) can be written as,

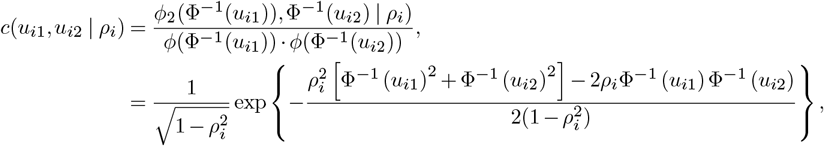

where Φ^−1^(·) is the quantile function(inversed CDF) of a standard univariate normal random variable, *ϕ*_2_(·, ·; ·) is the PDF of bivariate normal distribution. Φ^−1^(*u*_*i*1_) and Φ^−1^(*u*_*i*2_) follow a bivariate normal distribution with correlation matrix ***ϑ***_*i*3_:

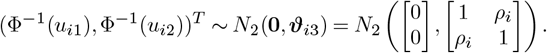

In addition to studying the zero-inflation rate along the temporal trajectory, we incorporate gene zero-inflation characteristics into the model. Define *D*_*ij*_ ~ Bern(*p*_*ij*_), *j* = 1, 2, be the random variables representing whether the expression measurement for gene *j* is zero in cell *i*. Then we can model the measured values of the expression of the two genes in cell *i* as:

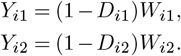

The joint CDF of the expression measurements of a gene pair can be written as:

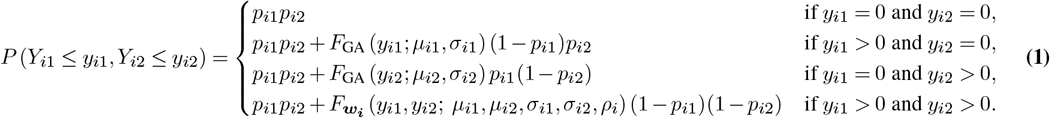

The log-likelihood function can be expressed as,

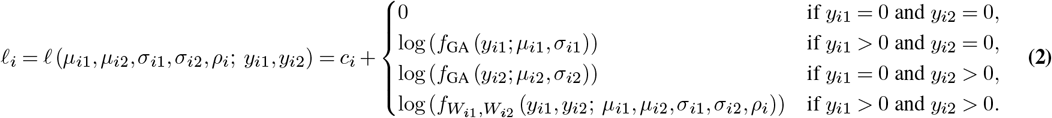

where *c*_*i*_ is the additive constant containing *p*_*i*1_ and *p*_*i*2_. The estimation of the parameters *µ*_*i*1_, *µ*_*i*2_, *σ*_*i*1_, *σ*_*i*2_, *ρ*_*i*_ does not depend on the values of *p*_*i*1_ and *p*_*i*2_. The details of how to get Eq. (1) and Eq. (2) are shown in Supplementary Data—Appendix A2. For *N* cells, the overall log-likelihood function can be expressed as 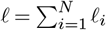.

#### 2.2.2 Additive predictor

Define the parameter vector as ***ϑ*** = (*θ*_1_, …, *θ*_*M*_)^*T*^, where *M* is the number of parameters. In the real data analysis, there are 7 parameters ***ϑ*** = (*µ*_1_, *σ*_1_, *µ*_2_, *σ*_2_, *ρ, p*_1_, *p*_2_)^*T*^. It includes the marginal distribution parameters of Gamma distributions, the copula dependence parameter which captures the correlation between the expression levels of two genes, and the zero-inflation rates for each gene. Let **Z** = (***z***_**1**_, …, ***z***_***K***_)^⊤^as *K* covariates, each covariate is a vector with *N* values for *N* cells. Under mouse pituitary gland scRNAseq data analysis, there are two covariates, the cell pseudotime and the mutant status for each cell.

To study non-linear changes associated with cellular temporal trajectory, each parameter could relate to the covariates in a flexible nonparametric fashion in the proposed framework. The marginal distribution parameters, the copula dependence parameter, and the zero-inflation rates are modeled by their additive predictors denoted as 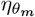 for *m* = 1, …, *M* through one-to-one monotonic and differentiable transformation functions to guarantee the parameters are within a valid range. For distributional parameter *µ* and *σ*, we use log() as link function to ensure *µ* and *σ* are postive values. To make sure the correlation of two gene expression levels has a range from −1 to 1, we use the inverse hyperbolic tangent function as the link function for parameter *ρ*. For zero-inflation rate *p*, we use logit() as a link function to make sure the range of *p* is from 0 to 1.

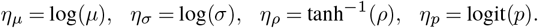

Hence, the inverse link functions on additive predictors related to the model parameters as follows:

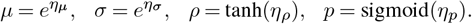

These additive predictors can be expressed as an intercept and a series of smoothing functions that are related to covariates ***z***_***k***_. For each smoothing function, a series of splines are used in additive predictor for each parameter *θ*_*m*_. The choice of the splines can be thin plate splines, Duchon splines, cubic regression splines, B-splines, P-splines, random effects, Markov random fields, Gaussian process smooths, Soap film smooths, splines on the sphere. A general form of the predictor (29) for each parameter *θ*_*m*_ can be written as:

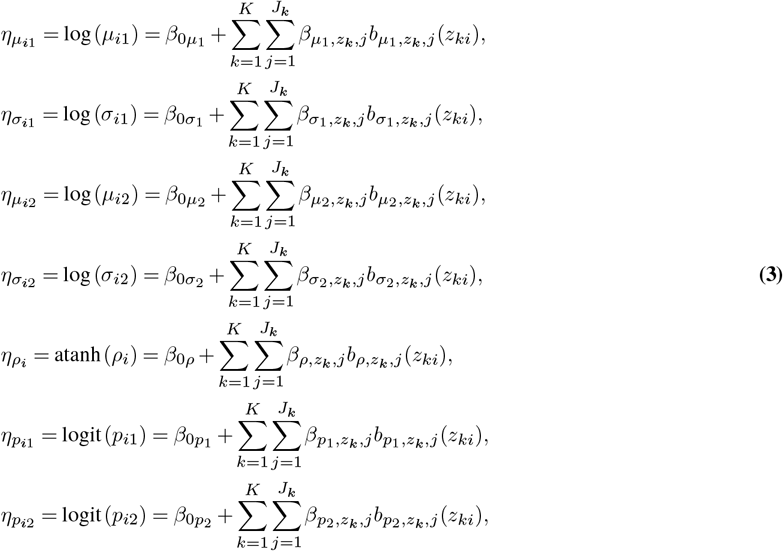

where *i* = 1, …, *N*; *J*_*k*_ is the number of basis functions for *k*th covariate, and *b*_·,*j*_(·) is the *j*th basis function of the spline associated with the particular parameter and covariates; *β*_0·_ and 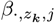 are denoted as the regression coefficients to be estimated. In the experimental data analysis, *K, z*_*i*1_ is the cell pesudotime for *i*th cell, and *z*_*i*2_ is the mutant status for that cell.

For each parameter, *θ*_*m*_, the additive predictor can be written as,

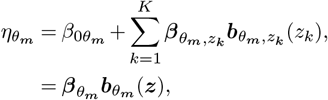

where 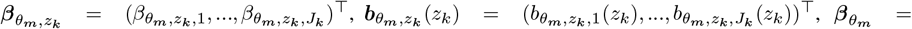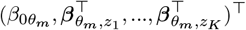 and 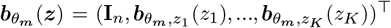. The parameters of our model can also be expressed in the following way:

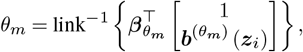

where 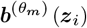 is the vector of basis functions corresponding to *θ*_*m*_ evaluated at their respective subvectors of ***z***_*i*_. The coefficients to be estimated in the model are thus 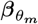.

Estimating zero-inflation rates *p*_1_ and *p*_2_ is different from estimating other parameters. The result from Eq. (4) allows us to estimate ***β***_*p1*_ and ***β***_*p2*_ using logistic regression on **1** {*Y*_*ij*_ = 0}, while ***β***_*µ*_, ***β***_*µ*_, ***β***_*σ*_, ***β***_*σ*_, ***β***_*ρ*_ are estimated by maximizing 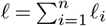. This maximization is achieved using the trust algorithm as detailed in Section 2.2.4, which requires the gradient and hessian to be explicitly calculated. The detailed model setting for our simulation and experimental data analysis is shown in the Supplementary Data—Appendix A2.

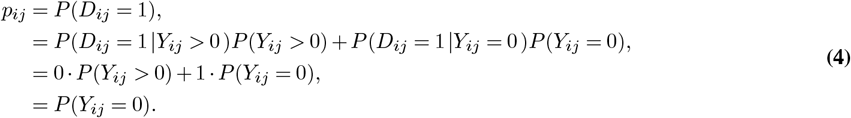

#### 2.2.3 Penalization

The model maximizes the penalized log-likelihood instead of likelihood to avoid over-fitting, it is defined as follows,

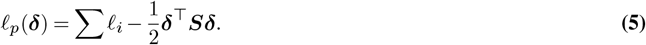

Define 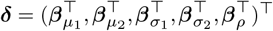 as a coefficient vector that combines all coefficients *β* corresponding to the 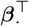 in Eq. (3). 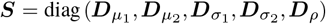; with 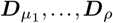 being block-diagonal matrices containing smoothing parameter **λ** that balances the degree of smoothing and the model fitting; they are constructed according to the particular type of smooth used in the corresponding predictor.

#### 2.2.4 Parameter estimation

We implement the extended trust region algorithm (30, 31) to maximize the penalized model’s log-likelihood in Eq. (5) for the estimations of additive predictors ***η*** and smoothing parameters **λ**. The process for estimating the model is,

1. Initialize the smoothing parameters λs.
2. At iteration *a*, hold **λ**^[*a*]^ as constants at their values from the last iteration or step (1) if it’s the first iteration. Trust region algorithm seeks to minimize *ℓ*_*p*_ in Eq. (5), it’s equivalent to find ***δ***^[*a*+1]^ that minimize a simpler quadratic model that contained the penalized gradient and Hessian of Eq. (5).
3. Holding ***δ***^[*a*+1]^ constant at the values found in step (2), find the value of **λ**^[*a*+1]^ that minimizes the special equation involving the hessian and gradient.
4. Repeat steps (2) and (3) until *ℓ*_*p*_ converges.

To maximize *ℓ*_*p*_ with respect to ***δ*** as called for in step 2, we utilize trust region optimization. The trust region algorithm evaluates the objective function only after solving the trust region problem, rather than repeatedly estimating it. This makes it faster and more reliable than standard approaches such as Newton-Raphson (30, 32). It requires both the gradient and hessian of the objective function, they are shown in Supplementary Data—Appendix A3 and A4. The smoothing criterion referenced in step 3 is that of Marra et al. (31), which provides stable and efficient results for copula models whose parameters depend on covariates in a flexible, possibly non-linear fashion.

The spline we used in this framework is the thin plate regression spline. Other splines mentioned in Section 2.2.2 can also be used. To get the estimated parameters 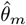, plug in the estimated additive predictors 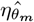 into the inverse link functions, 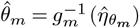. The detailed estimation process is described in Supplementary Data—Appendix A5.

#### 2.2.5 Model building

The mouse pituitary gland embryological development scRNAseq data contains two groups: wild-type and *Nxn*-mutated mice data. We use a factor group with values 0 and 1 to represent the wild-type and mutant groups respectively. For a given gene pair, the model includes 7 equations: two for mean expression levels, two for standard deviation, two for zero-inflation rates, and one for correlation between the gene pair. They are shown below,

~~~
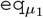 <-gene_1_ ~ group + s(t, by=group)
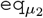 <-gene_2_ ~ group + s(t, by=group),
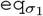 <-~ s(t),
eq_*σ*2_ <-~ s(t),
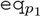 <-~ group + s(t, by=group),
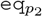 <-~ group + s(t, by=group),
eq_*ρ*_ <-~ group + s(t, by=group),
eqlist <-list(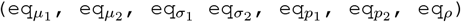),
output <-TIME_CoExpress(formula=eqlist, data, copula=“N”, model=“B”, margins=c(“GA”,”GA”), …),
~~~

where t is the cell pseudotime. `gene`_1_ and `gene`_2_ are the expression levels of two genes. `s()` is the smooth function that models the non-linear changes in distributional parameters over pseudotime. The number of basis functions can be set inside `s()`. The Gaussian Copula model with Gamma margins is employed in the following analysis.

We use quantile residuals to evaluate the goodness of fit for our model. For the non-zero continuous outcomes, the normalized quantile residuals are given by 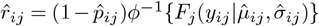, for *i* = 1, …,*N* cells, *j* = 1, 2, where 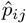 is the predicted zero-inflation rate of gene *j* in cell *i* (29). For the observed zero values, we sample unifrom variables from the range 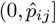 as residuals (33).

### 2.3 Simulation

This section presents the details of the three simulation studies conducted to evaluate the performance of the TIME-CoExpress framework and compares the proposed model with other methods. The purpose of this simulation study is to evaluate the performance of the proposed algorithm in terms of bias and statistical power.

#### Scenario I

We first tested the proposed framework in a linear setting where the additive predictor has a linear relationship with cell pseudotime *x* showed in Eq. (6). Assume *σ*_1_ = 0.2, and *σ*_2_ = 0.27. The zero-inflation levels for gene *y*_1_ and gene *y*_2_ are 0.45 and 0.3 respectively. Cell pseudotime *x* is simulated from a uniform distribution (0,27). The number of observations is set to 500 and 1,000. The number of iterations is 1,000. The results are shown in Figure 2. The coefficient estimations are more accurate with a larger sample size. Overall, the proposed framework has a good performance in a linear setting.

**Fig. 2.**
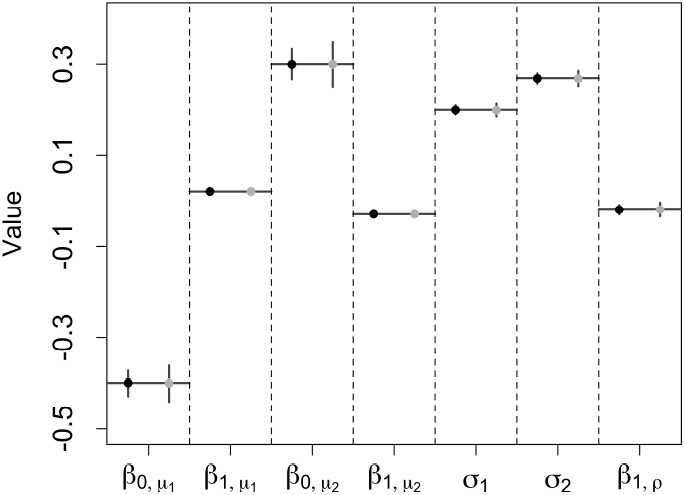
Scenario I simulation: The gray color is for data with 500 observations and the black color is for data with 1,000 observations. The horizontal lines are the true coefficients, the dots are the coefficient estimations. The vertical lines represent the 5% to 95% quantile.

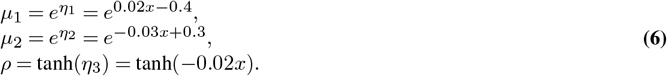

#### Scenario II

We conducted the simulation studies in two non-linear scenarios. In the first non-linear scenario, zero-inflated data is generated to mimic the real mouse pituitary gland embryological development data. The responses *y*_1_ and *y*_2_ are the two genes’ expression levels, and they are simulated from Gamma distributions with zero inflation rates *p*_1_ = 0.4 and *p*_2_ = 0.35 respectively. The covariate *x* is the cellular pseudotime, we sampled *x* from the uniform distribution with interval (0, 27).

We used Gaussian copula distribution to model the correlation *ρ* of two genes’ expression levels *y*_1_ and *y*_2_, the marginal distribution of *y*_1_ and *y*_2_ are Gamma distributions, assuming *σ*_1_ = 0.2, and *σ*_2_ = 0.27. Assume the *µ*_1_, *µ*_2_ and *ρ* have non-linear relationships with cell pseudotime *x*. The link functions for *η*_1_ and *η*_2_ are log functions. Use the inverse hyperbolic tangent function as the link function for the dependence parameter *η*_3_. The parameter models are given by,

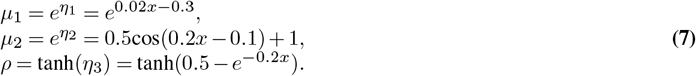

The hyperbolic tangent function for dependence parameter *ρ* can rescale the additive predictor to (−1, 1). The exponential function is used for marginal distribution parameters to make sure the mean and variance are positive values. In the simulation, we chose the thin plate regression spline. The number of iterations is set to 1,000, and the number of observations is set to 1,000 and 3,000. In each iteration, a new zero-inflated data set is generated and fitted using the proposed model. The fitted curves are shown in Figure 3.

**Fig. 3.**
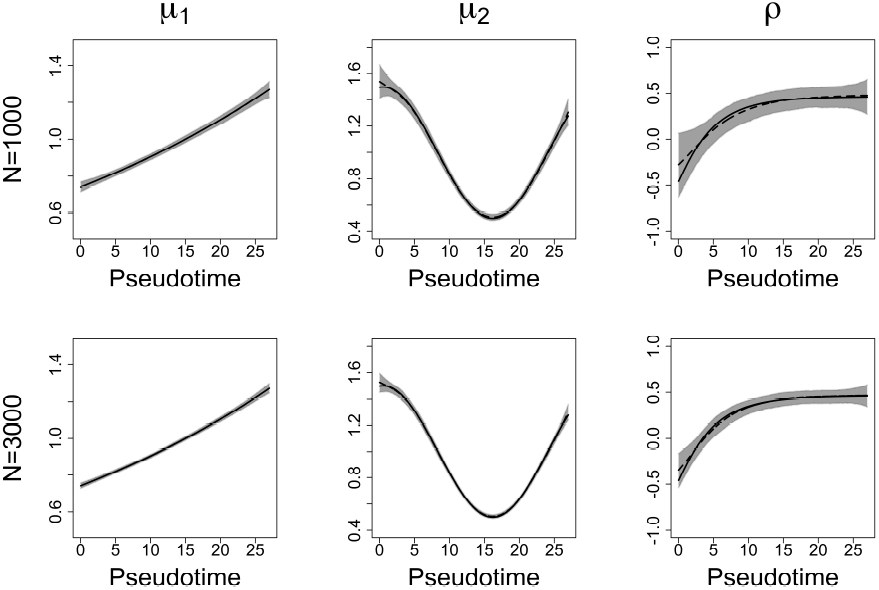
Scenario II simulation: The black solid lines are the true smooth functions, and the black dashed lines are the mean estimates for 1,000 iterations. The shaded areas are point-wise ranges from 5% to 95% quantile. The numbers of observations are 1,000 and 3,000 for each row.

In Figure 3, The black solid lines are the true smooth functions, and the black dashed lines are the mean estimates for 1,000 iterations. The shaded areas are point-wise ranges from 5% to 95% quantile and it covers the true smooth functions. The figure on the left is for *µ*_1_, the figure in the middle is for *µ*_2_, and the one on the right is for correlation *ρ*. The plots of *σ*_1_, *σ*_2_, *p*_1_, and *p*_2_ are shown in Supplementary Data-Appendix B1.

In simulation scenario II, the mean expression level of the first gene increases as the cell pseudotime increases, while the mean expression level of the second gene decreases at the first beginning, it reaches the lowest level around pseudotime 16, then increases. The correlation between these two genes is negative at first since the two genes’ expression levels change oppositely, and it becomes positive when these two genes’ expression levels have the same trend. The average time to run one iteration is 0.111 minutes (6.66 seconds) for 1,000 observations; and 0.176 minutes (10.56 seconds) for 3,000 observations. This simulation is conducted on a high-performance computing cluster, with each node configured with 24 CPUs (cores).

We evaluated the fitting results by calculating Bias and Root Mean Squared Error(RMSE) for each parameter. Bias is defined as 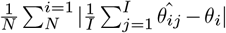 and RMSE is defined as 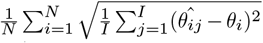, where *N* is the number of observations; *I* = 1, 000 is the number of iterations; *θ* is the parameter; 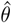 is the parameter estimation. The Bias and RMSE for each parameter are shown in Table 1. The results suggest that the TIME-CoExpress has low bias and RMSE.

**Table 1.** Bias and RMSE for each parameter in simulation Scenario II.

|  | Parameter | Bias | RMSE |
| --- | --- | --- | --- |
| $N = 1,000$ | $\mu_1$ | 0.001 | 0.013 |
| | $\mu_2$ | 0.006 | 0.026 |
| | $\sigma_1$ | 0.001 | 0.006 |
| | $\sigma_2$ | 0.002 | 0.008 |
| | $\rho$ | 0.023 | 0.093 |
| | $p_1$ | 0.001 | 0.015 |
| | $p_2$ | 0.000 | 0.015 |
| $N = 3,000$ | $\mu_1$ | 0.000 | 0.008 |
| | $\mu_2$ | 0.004 | 0.016 |
| | $\sigma_1$ | 0.000 | 0.003 |
| | $\sigma_2$ | 0.001 | 0.004 |
| | $\rho$ | 0.013 | 0.055 |
| | $p_1$ | 0.000 | 0.009 |
| | $p_2$ | 0.000 | 0.009 |

#### Scenario III

In this simulation scenario, we simulated zero-inflated data to mimic the experimental dataset where we have the wild-type group data and the mutant group data. For each cell *i*, we defined cell pseudotime as *z*_*i*1_ and we introduced an additional factor *z*_*i*2_ to identify the two groups of data. The cell is from the wild-type group if *z*_*i*2_ = 0 and it’s from the mutant group if *z*_*i*2_ = 1.

For each cell *i*, we simulated the non-zero values of the two gene expression levels *w*_*i*1_ and *w*_*i*2_ from a Gaussian copula model mentioned in Section 2.2.1 with gene pair correlation *ρ*_*i*_. Both marginal distributions are set to Gamma distributions with distributional parameters (*µ*_*i*1_, *σ*_*i*1_) and (*µ*_*i*2_, *σ*_*i*2_) respectively. The zero-inflation rates of two genes are *p*_*i*1_ and *p*_*i*2_, they both have a non-linear relationship with cell pseudotime *z*_*i*1_. To incorporate the zero-inflation characteristic in our simulation, we simulate *D*_*i*1_ ~ Bern(*p*_*i*1_) and *D*_*i*2_ ~ Bern(*p*_*i*2_). The gene expressions of the gene pair will be *y*_*i*1_ = (1 − *D*_*i*1_)*w*_*i*1_ and *y*_*i*2_ = (1 − *D*_*i*2_)*w*_*i*2_.

We assumed the mean of gene expression level(*µ*), the correlation of two genes(*ρ*), and the gene zero-inflation rates(*p*) change in a non-linear fashion along the cell pseudotime *z*_1_. The parameter simulations are given by,

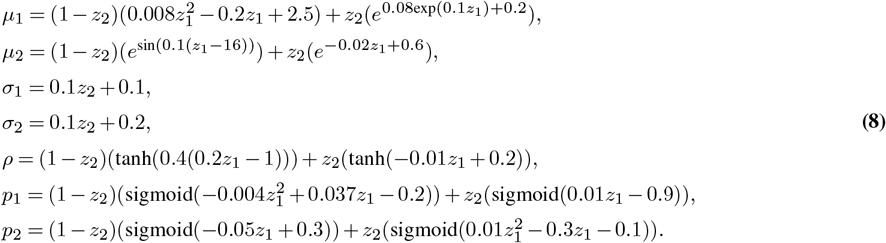

We sampled cell pseudotime *z*_1_ from a uniform distribution with interval (0, 27) based on the experimental data analysis described in Section 3. The simulated functions in Eq. (8) and the parameters are also based on the mice pituitary gland scRNAseq data considered in Section 3. The number of iterations is set to 1,000. The sample size is set to 2,000 or 5,000. The proportions of wild-type group and mutant group data are set to 55% and 45%. The results are shown in Figure 4. The standard deviation plots will be shown in Supplementary Data—Appendix B1.

**Fig. 4.**
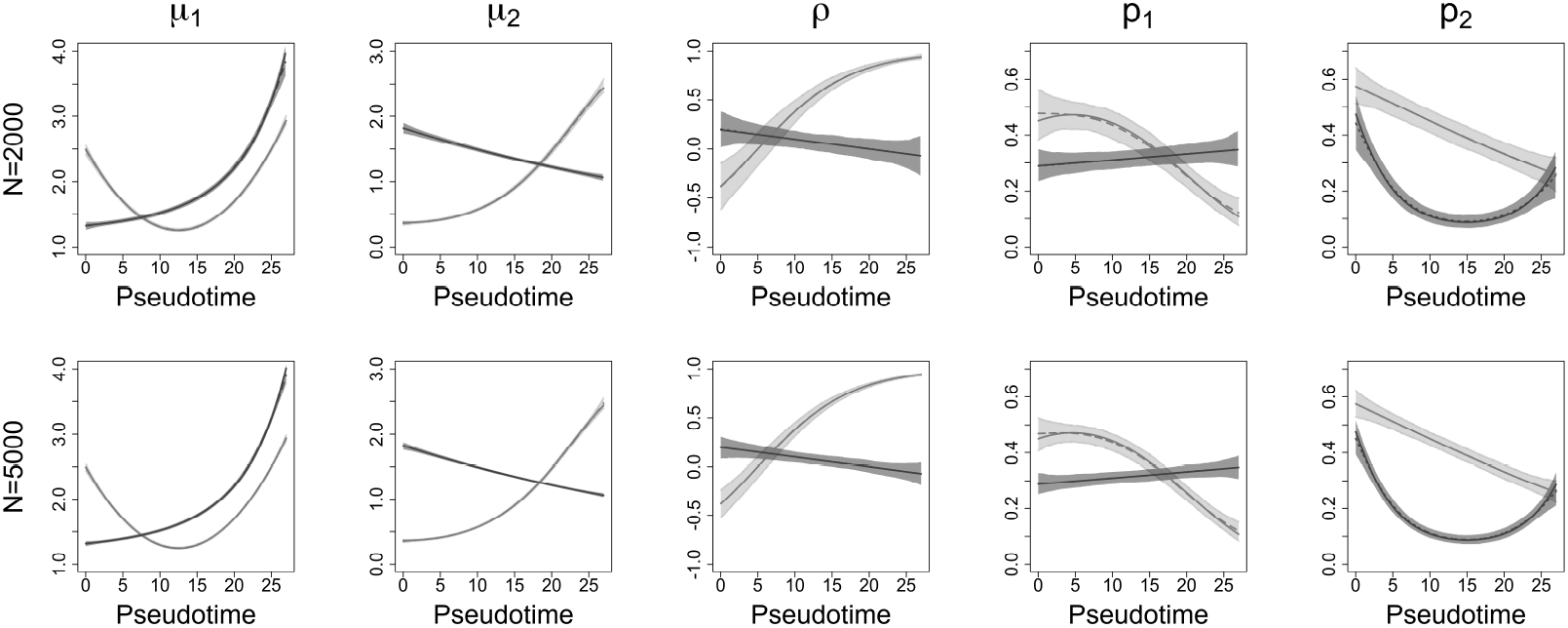
Scenario III simulation: The light gray color is for the wild-type group, and the dark gray color is for the mutant group. The solid lines are the true smooth functions, and the dashed lines are the mean estimates for 1,000 iterations. The shaded areas are point-wise ranges from 5% to 95% quantile. The numbers of observations are 2,000 and 5,000 for each row.

In Figure 4, the light gray color is for the wild-type group, and the dark gray color is for the mutant group. The solid lines are the true smooth functions, and the dashed lines are the mean estimates for 1,000 iterations. The shaded areas are point-wise ranges from 5% to 95% quantile. The Bias and RMSE of each parameter for both the wild-type group and the mutant group are shown in Table 2. The average time to run one iteration is 1.558 minutes (93.480 seconds) for 2,000 observations; and 1.117 minutes (67.020 seconds) for 5,000 observations.

**Table 2.** Bias and RMSE for each parameter in simulation Scenario III.

|  | Parameter | Wild-type |  | Mutant |  |
| --- | --- | --- | --- | --- | --- |
|  |  | Bias | RMSE | Bias | RMSE |
| $N = 2,000$ | $\mu_1$ | 0.005 | 0.017 | 0.008 | 0.034 |
| | $\mu_2$ | 0.005 | 0.019 | 0.000 | 0.024 |
| | $\sigma_1$ | 0.001 | 0.003 | 0.001 | 0.005 |
| | $\sigma_2$ | 0.002 | 0.006 | 0.001 | 0.007 |
| | $\rho$ | 0.003 | 0.062 | 0.001 | 0.071 |
| | $p_1$ | 0.066 | 0.074 | 0.000 | 0.024 |
| | $p_2$ | 0.001 | 0.028 | 0.006 | 0.023 |
| $N = 5,000$ | $\mu_1$ | 0.003 | 0.011 | 0.006 | 0.024 |
| | $\mu_2$ | 0.004 | 0.013 | 0.000 | 0.016 |
| | $\sigma_1$ | 0.000 | 0.002 | 0.000 | 0.003 |
| | $\sigma_2$ | 0.001 | 0.004 | 0.000 | 0.004 |
| | $\rho$ | 0.001 | 0.037 | 0.001 | 0.044 |
| | $p_1$ | 0.066 | 0.070 | 0.000 | 0.015 |
| | $p_2$ | 0.000 | 0.017 | 0.004 | 0.015 |

The results from both scenarios II and III suggest that the 5% to 95% quantile bounds cover the true smooth functions, and the mean estimates are close to the true smooth functions. The Bias and the RMSE are small which means the estimations are close to the real smoothing function. The Bias and the RMSE are smaller with a larger number of observations. Through the simulation studies, we show that the TIME-CoExpress model can capture the non-linear relationship between the correlation of responses and the covariates.

#### Scenario IV

In this Scenario, we evaluated the statistical power of our proposed framework by comparing it with scHOT (1), CNM (18), and liquid association (LA) proposed by Li (17). We designed a simulation with a correlation function shown in Eq. (9).

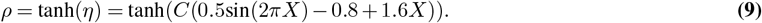

This function contains a constant scaling factor *C*. When *C* is small, the correlation function is close to zero; as *C* increases, the correlation function becomes more and more different from *y* = 0. We set the means of two genes’ expressions as *µ*_1_ = 5, *µ*_2_ = 3 and standard deviations 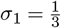 and 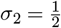. The zero-inflation rates for these two genes are *p*_1_ = 0.3 and *p*_2_ = 0.4.

The null hypothesis is defined as,

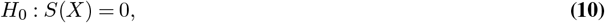

where *X* is the cell pseudotime, function *S*() is the correlation function. We simulated 1,000 observations, and fit these four methods 1,000 times for each value of *C*. We calculated the model power, defined as the proportion of simulations rejecting the null hypothesis, for different values of constant *C*. The resulting power plot is shown in Figure 5. The results present in Figure 5 show that when *C* = 0, all methods maintain proper type I error wild-type at the nominal level (0.05). In addition, this power analysis also indicates that our proposed method outperforms the other three methods. The power of our method reaches 1 faster as *C* increases compared to other methods.

**Fig. 5.**
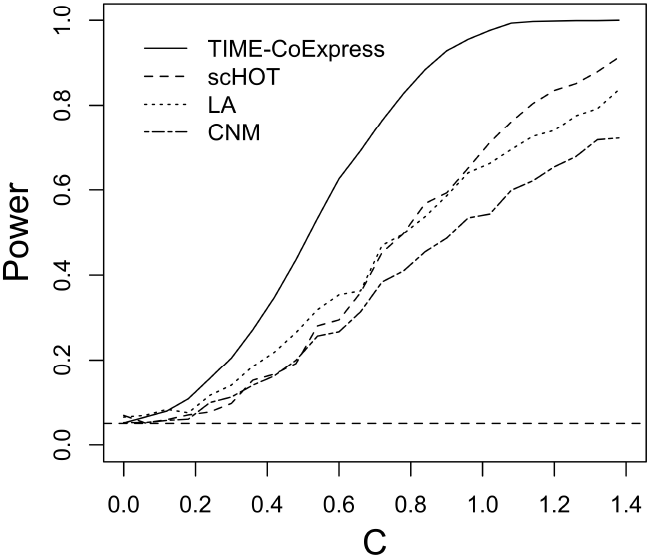
Scenario IV simulation: The power comparison of TIME-CoExpress, scHOT, CNM, and LA (with permutations). The number of observations is 1,000, the number of iterations is 1,000. The power is defined as the rate of rejecting the null hypothesis when testing *S*(*X*) = 0.

## 3 Experimental data analysis

In this section, we applied the TIME-CoExpress model to embryological scRNAseq data described in Section 2.1. Our analysis aims to investigate the impact of *Nxn* mutation on the development of the mouse pituitary gland by identifying changes in gene co-expression along cellular temporal trajectories between wild-type and mutant mice, and incorporates changes in zero inflation rate along the cell temporal trajectory.

We considered two lists of genes. The first list of genes is from the WNT signaling pathway (mmu04310). *Nxn* is known to regulate Disheveled, a key intracellular mediator of the WNT signaling pathway (25). *Wnt5a* is expressed in the embryonic pituitary gland and *Wnt5a*^−*/*−^ embryos have malformed pituitary glands (34). Therefore, we selected genes from the *Wnt5a* gene pathway for analysis using TIME-CoExpress. We removed genes from the gene list with more than 80% zeros since genes with zero expression in most cells do not provide any information. 50 genes (1225 gene pairs) are selected. We studied all possible gene pair combinations by fitting our model to each pair and identified gene pairs with different co-expression patterns between wild-type and mutant groups along the cellular pseudotime. The second gene list is from pituitary transcription factors and we mainly focused on evaluating zero-inflation rates for each gene along pseudotime. 55 pituitary gland transcription factors (1485 combinations) are used. The lists of the 50 genes related to the WNT signaling pathway and the 55 TF can be found in Supplementary Data—Appendix B5.

We applied the Slingshot algorithm to estimate cell the temporal trajectory for all cells. This method fits a single principal curve for all cells based on gene expression levels and produces a pseudotime ordering of cells, which is determined by their positions when projected onto the curve. The cell cluster information is not provided to Slingshot since it is an unsupervised method. Other pseudotime inference methods like TSCAN (7), Monocle (6), etc. can also be used.

For each gene pair with expression levels *y*_1_ and *y*_2_, we fit the proposed framework with cell pseudotime (*z*_1_), and group (*z*_2_). The model provided correlation estimates for each gene pair along pseudotime for both wild-type and mutant groups. This multi-group analysis enables a direct comparison between different groups. To evaluate the difference between the two correlation curves, a p-value is calculated to test the null hypothesis that the correlation trajectories between the wild-type and mutant groups are identical. Besides the gene pair co-expressions, our model also evaluates how zero-inflation rates change through cell pseudotime for individual genes. For each gene, our model provides p-values to assess whether the zero-inflation rate differs significantly between the wild-type and mutant groups. To control the False Discovery Rate (FDR), we use the Benjamini-Hochberg (BH) procedure to adjust the p-values across all gene pairs. We consider the gene pairs with adjusted p-values less than 0.05 for significance. 42 gene pairs were declared to have significant temporal co-expression changes between wild-type and mutant among all 1,225 gene pairs from the WNT pathway.

We also considered a second metric to evaluate the difference between the estimated correlation curves 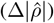:

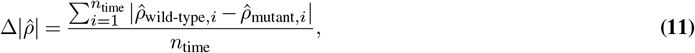

where *n*_time_ is the number of pseudotime values we generated, 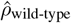 and 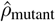 are the gene pair correlation predictions for the wild-type group and the mutant group respectively. Due to inherent differences between cells in the wild-type and mutant groups, their respective cell pseudotimes varied. To facilitate a meaningful comparison of the dynamic correlation between these two groups, we generate evenly spaced pseudotime to predict the dynamic correlation using the fitted model to calculate 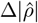.

Table 3 shows the 42 significant gene pairs co-expression with the smallest adjusted p-value between the wild-type group and the mutant group. 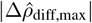 is defined as the maximum predicted correlation difference between these two groups. The range of correlation is between −1 and 1, and the max number 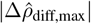 can be no larger than 2. *AUD* is the difference in the area under the predicted correlation curves between the mutant versus wild-type. The larger the AUD, the greater the difference between the two predicted correlation curves.

**Table 3.** 42 significant gene pairs with adjusted p-value less than 0.05. 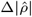 is the overall correlation difference between the wild-type group and the mutant group; 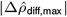 is the max predicted correlation difference between these two groups; AUD is the area under the correlation difference curve.

| | Gene 1 | Gene 2 | P-value | Adjusted P-value | $\Delta\vert \hat{\rho}\vert $ | $\vert \Delta\hat{\rho}_{\text{diff,max}}\vert $ | AUD |
| --- | --- | --- | --- | --- | --- | --- | --- |
| 1 | *Apc* | *Fbxw11* | <0.001 | <0.001 | 0.270 | 0.579 | 8.100 |
| 2 | *Apc* | *Senp2* | <0.001 | <0.001 | 0.365 | 0.814 | 10.945 |
| 3 | *Btrc* | *Ctbp1* | <0.001 | <0.001 | 0.303 | 0.617 | 9.095 |
| 4 | *Btrc* | *Fzd3* | <0.001 | <0.001 | 0.425 | 0.901 | 12.728 |
| 5 | *Btrc* | *Lef1* | <0.001 | <0.001 | 0.490 | 0.994 | 14.688 |
| 6 | *Btrc* | *Senp2* | <0.001 | <0.001 | 0.750 | 1.446 | 22.475 |
| 7 | *Camk2d* | *Tbl1x* | <0.001 | <0.001 | 0.373 | 0.643 | 11.184 |
| 8 | *Ccnd2* | *Lrp6* | <0.001 | <0.001 | 0.242 | 0.634 | 7.251 |
| 9 | *Ccnd3* | *Siah1a* | <0.001 | <0.001 | 0.105 | 0.250 | 3.156 |
| 10 | *Crebbp* | *Fbxw11* | <0.001 | <0.001 | 0.523 | 0.891 | 15.666 |
| 11 | *Csnk1e* | *Lgr4* | <0.001 | <0.001 | 0.165 | 0.344 | 4.938 |
| 12 | *Csnk1e* | *Siah1a* | <0.001 | <0.001 | 0.129 | 0.289 | 3.856 |
| 13 | *Csnk2a1* | *Lrp6* | <0.001 | <0.001 | 0.222 | 0.610 | 6.660 |
| 14 | *Csnk2a2* | *Rock2* | <0.001 | <0.001 | 0.427 | 0.952 | 12.789 |
| 15 | *Csnk2b* | *Siah1a* | <0.001 | <0.001 | 0.154 | 0.409 | 4.628 |
| 16 | *Ctbp1* | *Lrp6* | <0.001 | <0.001 | 0.131 | 0.258 | 3.941 |
| 17 | *Ctbp1* | *Ppp3cb* | <0.001 | <0.001 | 0.284 | 0.654 | 8.508 |
| 18 | *Ctnnb1* | *Jun* | <0.001 | <0.001 | 0.301 | 0.666 | 9.012 |
| 19 | *Ctnnbip1* | *Siah1a* | <0.001 | <0.001 | 0.108 | 0.337 | 3.247 |
| 20 | *Ctnnd2* | *Siah1a* | <0.001 | <0.001 | 0.459 | 0.825 | 13.759 |
| 21 | *Ep300* | *Lgr4* | <0.001 | <0.001 | 0.247 | 0.564 | 7.416 |
| 22 | *Fzd3* | *Ryk* | <0.001 | <0.001 | 0.169 | 0.394 | 5.061 |
| 23 | *Fzd3* | *Tle1* | <0.001 | <0.001 | 0.226 | 0.450 | 6.779 |
| 24 | *Gsk3b* | *Lgr4* | <0.001 | <0.001 | 0.175 | 0.366 | 5.232 |
| 25 | *Gsk3b* | *Prickle1* | <0.001 | <0.001 | 0.264 | 0.748 | 7.899 |
| 26 | *Gsk3b* | *Senp2* | <0.001 | <0.001 | 0.131 | 0.289 | 3.929 |
| 27 | *Lrp6* | *Prkacb* | <0.001 | <0.001 | 0.472 | 1.063 | 14.150 |
| 28 | *Plcb4* | *Prkaca* | <0.001 | <0.001 | 0.267 | 0.616 | 7.998 |
| 29 | *Ppp3ca* | *Ryk* | <0.001 | <0.001 | 0.116 | 0.244 | 3.474 |
| 30 | *Ruvbl1* | *Senp2* | <0.001 | <0.001 | 0.296 | 0.578 | 8.864 |
| 31 | *Ccnd3* | *Fbxw11* | <0.001 | <0.001 | 0.143 | 0.295 | 4.293 |
| 32 | *Gsk3b* | *Tbl1x* | <0.001 | <0.001 | 0.297 | 0.555 | 8.898 |
| 33 | *Ctbp1* | *Fzd3* | <0.001 | <0.001 | 0.147 | 0.304 | 4.392 |
| 34 | *Rock2* | *Ryk* | <0.001 | <0.001 | 0.637 | 1.171 | 19.095 |
| 35 | *Ppp3ca* | *Senp2* | <0.001 | <0.001 | 0.166 | 0.455 | 4.979 |
| 36 | *Camk2d* | *Fbxw11* | <0.001 | <0.001 | 0.930 | 1.597 | 27.870 |
| 37 | *Csnk2a2* | *Lef1* | <0.001 | <0.001 | 0.172 | 0.370 | 5.160 |
| 38 | *Ctnnd2* | *Tcf7l2* | <0.001 | 0.003 | 0.519 | 0.966 | 15.571 |
| 39 | *Rac1* | *Senp2* | <0.001 | 0.004 | 0.317 | 0.541 | 9.506 |
| 40 | *Lgr4* | *Tcf7l2* | <0.001 | 0.023 | 0.212 | 0.544 | 6.342 |
| 41 | *Prkaca* | *Tcf7l2* | <0.001 | 0.030 | 0.643 | 1.223 | 19.279 |
| 42 | *Cby1* | *Siah1a* | <0.001 | 0.035 | 0.604 | 1.284 | 18.098 |

We show three gene paris as examples from our analysis (*Ctnnd2* and *Tcf712*; *Ccnd2* and *Lrp6*; *Ctnnd2* and *Siah1a*) in Figure 6. These selected gene pairs demonstrate significant correlation differences between wild-type (blue) and mutant group (red). The solid curves represent how gene pair correlations or gene zero inflation rates change along the cellular pseudotime in both the wild-type group and mutant group, where the x-axis values are the psedudotime generated from Slingshot. The shaded areas represent 95% CI of the fit. More gene pair correlation plots are available in Section Supplementary Data—Appendix B2.

**Fig. 6.**
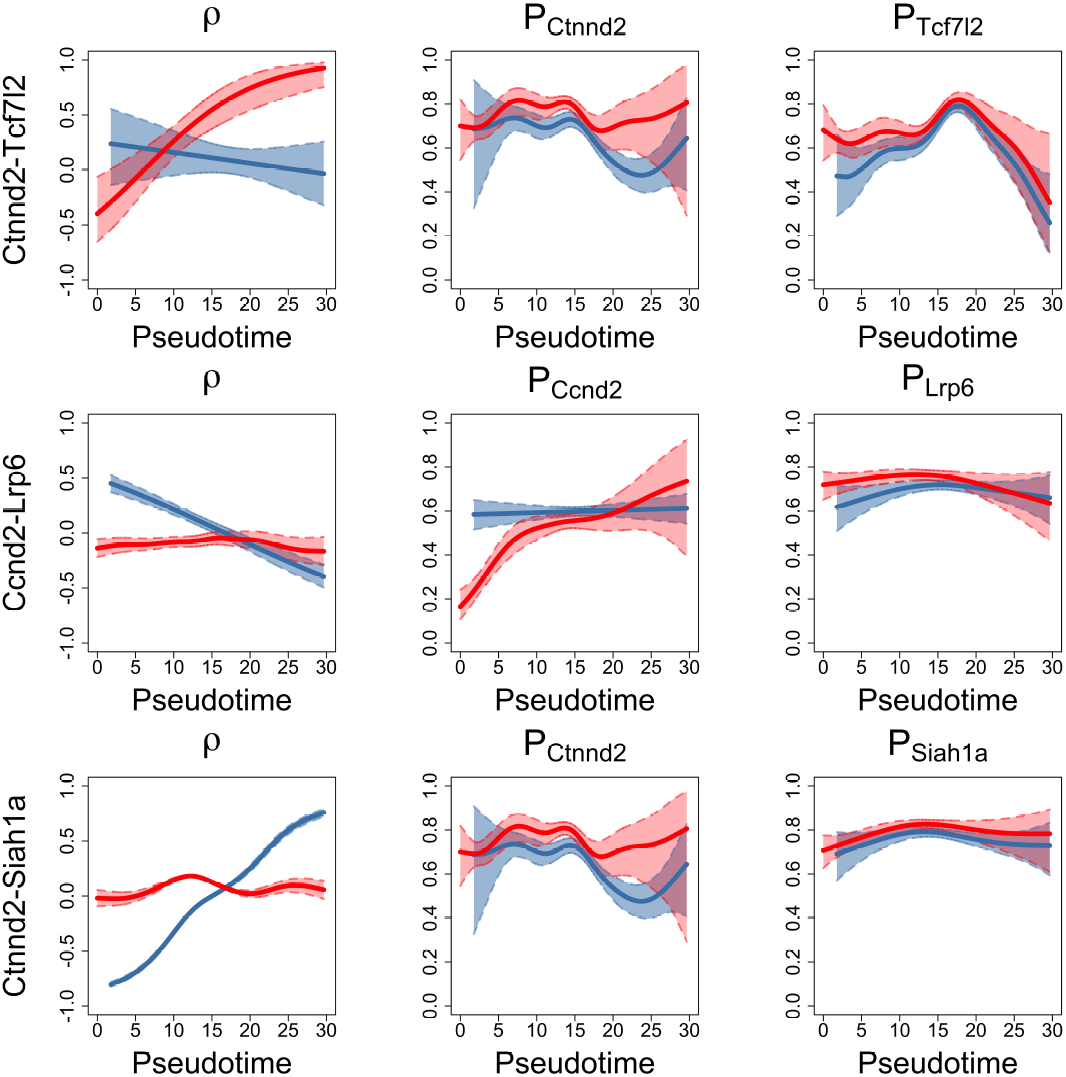
Significant gene pairs examples. The blue solid curve is the fitted line for the wild-type group, and the blue dashed line is 95% CI of the fit. The red solid curve is the fitted line for the mutant group, and the red dashed line is 95% CI of the fit.

We use Quantile-Quantile (Q-Q) plots to evaluate the model’s goodness-of-fit. It provides a visual comparison between the observed and predicted correlation distributions for each gene pair. The QQ plots for the gene pairs fitting are presented in Section Supplementary Data—Appendix B3. The data points align around the *y* = *x* line indicating the model accurately captured the distributional characteristics of the data across pseudotime.

Based on Table 3 and Figure 6, the TIME-CoExpress analysis identifies gene pairs that are likely involved in the differentiation process. For instance, *Ctnnd2*, or Delta-catenin, and the ubiquitin ligase *Siah1a* are negatively correlated in *Sox2*-expressing stem cells but have an increasing correlation as pituitary stem cells differentiate towards the *Pou1f1* expressing progenitor stage. In *Nxn*^−*/*−^ embryos these two genes’ expression patterns are not correlated. The expression correlation for *Ctnnd2* and the transcription factor *Tcf7l2* decreases in wild-type embryos during differentiation, while it is dramatically increased in *Nxn*^−*/*−^ embryos. These results suggest that *Ctnnd2* expression in combination with *Siah1a* promotes differentiation, while *Ctnnd2* expression with *Tcf7l2* maintains the stem cell state, which are experimentally testable hypotheses. The expressions of *Ccnd2* and *Lrp6* are correlated in stem cells in the control group, but this correlation weakens as differentiation progresses. In the mutant group, *Ccnd2* and *Lrp6* show no significant correlation. The distinct correlation patterns in the control and mutant groups emphasize the impact of the mutation on gene pair co-expression during differentiation.

In the gene zero-inflation analysis, we utilized transcription factors from the dataset to reveal how the zero-inflation rate changes for each gene along pseudotime. The figures in the first row in Figure 7 present how the zero-inflation rates of *Sox2, Prop1*, and *Pou1f1* progenitors change through cell pseudotime. The figures in the second row in Figure 7 present three TFs: *Gli2, Isl1, Lhx2* with a small p-value of zero inflation rates between mutant versus wild-type. More genes’ zero-inflation rates plots can be found in Supplementary Data—Appendix B4.

**Fig. 7.**
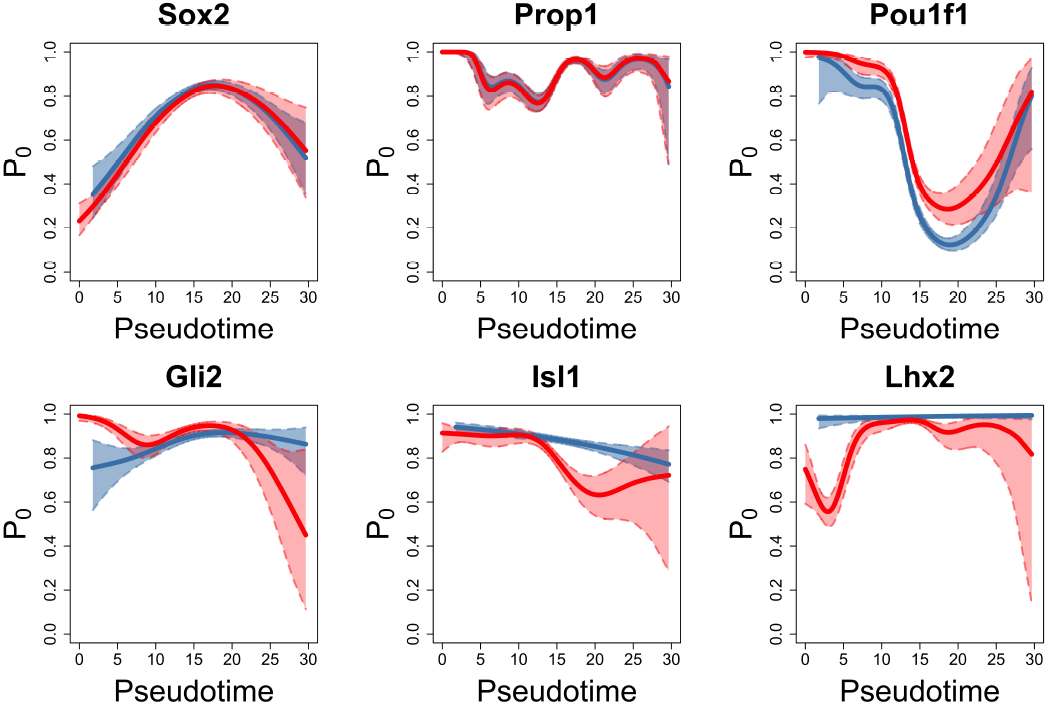
Examples of zero-inflation rate changes from important transcription factors. The blue solid curve is the fitted line for the wild-type group, and the blue dashed line is 95% CI of the fit. The red solid curve is the fitted line for the mutant group, and the red dashed line is 95% CI of the fit.

All analyses were performed using high-performance computing resources, with each gene pair computation averaging approximately 4.463 minutes (267.780 seconds). The computational setup consisted of 200 nodes running simultaneously, each node utilizing 24 CPUs (cores) to parallelize the workload. To test TIME-CoExpress, we limited the analysis by providing a relatively short gene list associated with WNT. Expansion of the gene list, potentially including all mouse genes, would increase the gene discovery capability and provide a more unbiased search. Because TIME-CoExpress is computationally efficient, the use of an expanded gene list for analysis is a practical approach.

## 4 Discussion

Understanding how gene pair co-expression changes with cellular temporal trajectory is crucial for gaining deeper insights into the dynamic interactions within biological systems. In this paper, we move beyond the traditional one-gene-at-a-time analysis and introduce a flexible, zero-inflated, copula-based, TIME-CoExpress analytical framework to capture the dynamic shifts in gene co-expression across cellular pseudotime.

Unlike traditional approaches, which often treat gene co-expression as fixed rather than time-dependent, TIME-CoExpress accounts for the non-linear characteristic of gene-to-gene interactions through a Gaussian copula-based model. This capability is important in biological analysis, as genes exhibit complex temporal changes in gene interactions within a living cell. Moreover, the proposed framework accommodates covariate-dependent zero-inflation. The zero-inflation rate represents a gene’s expression level, and observing how this rate changes across cell pseudotime can provide insights into gene activation or deactivation throughout cell differentiation. By incorporating the non-linear changes in a gene’s zero-inflation rate, TIME-CoExpress provides a more accurate representation of gene pair co-expression dynamics along the temporal trajectory.

The popularity of pseudotime inference enables researchers to investigate gene expression changes along a continuous cellular trajectory. In this paper, we use the Slingshot method (8) to infer cell pseudotime from scRNAseq data. Slingshot is compatible with handling bifurcations or multiple branching in cellular trajectories, and it is flexible to use because it does not require predefined clusters or starting points. Additionally, it is not sensitive to outliers and noise due to its approach to fitting principal curves. Other pseudotime inference methods, such as TSCAN (7), and Monocle (6), can also be applied based on the user’s needs. Our proposed framework is also applicable to real cell time if such data is available.

TIME-CoExpress is designed to analyze scRNAseq data from multiple groups simultaneously in a unified analytical framework. Whereas other methods, such as scHOT, can analyze only one group at a time and hence make comparisons between different treatment conditions difficult. TIME-CoExpress offers more comprehensive insights by including direct comparisons under different experimental conditions in the same model framework.

We conduct a series of simulation analyses to demonstrate the effectiveness of our copula-based framework. The results show that our proposed framework is capable of capturing dynamic correlation changes between two covariates over time. Additionally, the simulation studies illustrate that the framework effectively captures the dynamic zero-inflation patterns of genes as they change across cell pseudotime.

In the experimental data analysis, we apply the proposed model to experimental data generated from mouse pituitary gland embryogenesis. We compare wild-type mouse embryos with *Nxn* mutant embryos in terms of gene co-expression and zero-inflation rate along the cellular temporal trajectory. Our model includes two covariates: pseudotime with non-linear effects on gene pair co-expression, and mutant status as a factor to indicate the wild-type and mutant groups of data. Other covariates, such as cell status, can also be incorporated into the model.

Our model identified differential co-expression gene pair combinations along the cell temporal trajectory between *Nxn* mutant and wild-type mice and uncovers several gene pair interactions that may help explain the observed phenotypic differences. Our method provides a more detailed view of how genes interacted with each other over time during the mouse embryological development. By employing this innovative framework, we also uncover gene zero-inflation patterns that underlie developmental differences and demonstrate the dynamic transcriptional regulation that occurs during organogenesis of the mouse pituitary gland. The altered developmental trajectory for pituitary stem cells demonstrated here is consistent with and supported by experimental data (MLB, SAC, and SWD personal communication). Our TIME-CoExpress analysis identifies gene pairs whose transciptional changes may underlie the altered developmental trajectory.

The goodness of fit for our model is assessed through residual analysis and Q-Q plots, allowing us to visually compare the distribution of model-predicted gene co-expression values with observed data across cell pseudotime. The Q-Q plot of the residuals follows a nearly straight line, suggesting a good fit to the data and validating the correctness of our model in capturing the gene co-expression changes across cellular temporal trajectories.

Additionally, QQ plots serve as a useful tool for parameter tuning. There are a few parameter selections we need to consider in the zero-inflated copula model established in Section 2.2. The inflation factor of splines and the number of splines control the smoothness of the fitting. A larger inflation factor gives smoother curves. Through iterative visualization, we tune this parameter until the Q-Q plot indicates an optimal fit to guarantee our model accurately captures the dynamic, non-linear gene co-expression and zero-inflation patterns in the data. The splines we used in this model are the thin plate splines, other splines are also available to use.

The smoothing parameter λ controls the trade-off between fit and smoothness. Large λ gives high smoothness results but may underfit data. A small λ gives a better fit; however, it risks overfitting. Another parameter is the basis dimension. It affects if the model can capture the complexity of the underlying data. The number of basis functions also affects the fitting smoothness. With more basis functions, the more accurate the fitting results will be, but it may face overfitting issues.

The marginal distributions of the copula model are set to Gamma distributions since gene expressions in mouse pituitary gland embryological scRNAseq data are normalized continuous values and skewed. Other marginals can also be applied in the TIME-CoExpress model depending on the data under study. For example, use negative binomial distribution as marginals if the data is count-based. The Tweedie distribution is another option and can be easily implemented in our framework.

Our proposed method has higher power in detecting the difference between two correlation curves compared to scHOT (1), LA with permutations (17), and CNM (18). In addition to the improved power, our model is more computationally efficient. Under the simulation of 1,000 observations and 1,000 iterations, TIME-CoExpress achieves 11.23 seconds for each iteration on average, while scHOT, LA(with permutations), and CNM using a generalized estimation equations (GEE)-based approach (18) required approximately 84.48, 14.58, and 0.082 seconds respectively. CNM is a fast method, but the power is low. scHOT is computationally intensive compared to other methods.

## Availability of data and materials

Single-cell RNA sequencing data generated in association with this manuscript can be accessed through GEO Database, using series accession number GSE246211 (https://www.ncbi.nlm.nih.gov/geo/query/acc.cgi?acc=GSE246211) (35), and samples GSM7864906 (https://www.ncbi.nlm.nih.gov/geo/query/acc.cgi?acc=GSM7864906) (36), and GSM7864907 (https://www.ncbi.nlm.nih.gov/geo/query/acc.cgi?acc=GSM7864907) (37). The mutant data can be accessed using accession number GSE281783 and GSM8628052 (unpublished data).

## Acknowledgements

The authors appreciate Dr. Leonard Cheung’s contribution in giving advice on the initial scRNAseq analysis.

## References

1. Shila Ghazanfar, Yingxin Lin, Xianbin Su, David M. Lin, Ellis Patrick, Ze-Guang Han, John C. Marioni, and Jean Yee Hwa Yang. Investigating higher-order interactions in single-cell data with scHOT. Nature Methods, 17(8): 799–806, 2020.

2. Peter Langfelder and Steve Horvath. Wgcna: an r package for weighted correlation network analysis. BMC Bioinformatics, 9(1), 2008.

3. Jerome Friedman, Trevor Hastie, and Robert Tibshirani. Sparse inverse covariance estimation with the graphical lasso. Biostatistics, 9(3): 432–441, 2007.

4. Wenpu Lai, Yangqiu Li, and Oscar Junhong Luo. Mist: an interpretable and flexible deep learning framework for single-t cell transcriptome and receptor analysis. bioRxiv, 2024.

5. Runmin Wei, Siyuan He, Shanshan Bai, Emi Sei, Min Hu, Alastair Thompson, Ken Chen, Savitri Krishnamurthy, and Nicholas E. Navin. Spatial charting of single-cell transcriptomes in tissues. Nature Biotechnology, 40(8): 1190–1199, 2022.

6. Cole Trapnell, Davide Cacchiarelli, Jonna Grimsby, Prapti Pokharel, Shuqiang Li, Michael Morse, Niall J Lennon, Kenneth J Livak, Tarjei S Mikkelsen, and John L Rinn. The dynamics and regulators of cell fate decisions are revealed by pseudotemporal ordering of single cells. Nature Biotechnology, 32(4):381—386, 2014.

7. Zhicheng Ji and Hongkai Ji. TSCAN: Pseudo-time reconstruction and evaluation in single-cell RNA-seq analysis. Nucleic Acids Research, 44(13):e117–e117, 2016.

8. Kelly Street, Davide Risso, Russell B. Fletcher, Diya Das, John Ngai, Nir Yosef, Elizabeth Purdom, and Sandrine Dudoit. Slingshot: cell lineage and pseudotime inference for single-cell transcriptomics. BMC Genomics, 19(1): 477, 2018.

9. Koen Van den Berge, Hector Roux de Bézieux, Kelly Street, Wouter Saelens, Robrecht Cannoodt, Yvan Saeys, Sandrine Dudoit, and Lieven Clement. Trajectory-based differential expression analysis for single-cell sequencing data. Nature Communications, 11 (1):1201, 2020.

10. Ruochen Jiang, Tianyi Sun, Dongyuan Song, and Jingyi Jessica Li. Statistics or biology: the zero-inflation controversy about scrna-seq data. Genome Biology, 23, 2022.

11. Zhen Yang and Yen-Yi Ho. Modeling dynamic correlation in zero-inflated bivariate count data with applications to single-cell rna sequencing data. Biometrics, 78(2): 766–776, 2022.

12. Li Yieng Lau, Antonio Reverter, Nicholas J. Hudson, Marina Naval-Sánchez, Marina R S Fortes, and Pâmela Almeida Alexandre. Dynamics of gene co-expression networks in time-series data: a case study in Drosophila melanogaster Embryogenesis. Frontiers in Genetics, 11:517, 2020.

13. Nicolai Hans, Nadja Klein, Florian Faschingbauer, Michael Schneider, and Andreas Mayr. Boosting distributional copula regression. Biometrics, 79(3): 2298–2301, 2022.

14. Hirotaka Matsumoto, Hisanori Kiryu, Chikara Furusawa, Minoru S H Ko, Shigeru B H Ko, Norio Gouda, Tetsutaro Hayashi, and Itoshi Nikaido. SCODE: an efficient regulatory network inference algorithm from single-cell RNA-Seq during differentiation. Bioinformatics, 33(15): 2314–2321, 2017.

15. R. A. Rigby and D. M. Stasinopoulos. Generalized additive models for location, scale and shape. Journal of the Royal Statistical Society Series C: Applied Statistics, 54(3): 507–554, 2005.

16. Charles J. Geyer. trust: Trust Region Optimization, 2020. R package version 0. 1-8.

17. Ker-Chau Li. Genome-wide coexpression dynamics: Theory and application. Proceedings of the National Academy of Sciences, 99(26): 16875–16880, 2002.

18. Yen-Yi Ho, Giovanni Parmigiani, Thomas A. Louis, and Leslie M. Cope. Modeling liquid association. Biometrics, 67(1): 133–141, 2011.

19. Julian Martinez-Mayer, Michelle L. Brinkmeier, Sean P. O’Connell, Arnold Ukagwu, Marcelo A. Marti, Mirta Miras, Maria V. Forclaz, Maria G. Benzrihen, Leonard Y. M. Cheung, Sally A. Camper, Buffy S. Ellsworth, Lori T. Raetzman, Maria I. Perez Millan, and Shannon W. Davis. Knockout mice with pituitary malformations help identify human cases of hypopituitarism. Genome Medicine, 16, 2024.

20. Saishu Yoshida, Takako Kato, Takao Susa, Li yi Cai, Michie Nakayama, and Yukio Kato. PROP1 coexists with SOX2 and induces PIT1-commitment cells. Biochemical and Biophysical Research Communications, 385(1): 11–15, 2009.

21. Lorin E. Olson, Jessica Tollkuhn, Claudio Scafoglio, Anna Krones, Jie Zhang, Kenneth A. Ohgi, Wei Wu, Makoto M. Taketo, Rolf Kemler, Rudolf Grosschedl, Dave Rose, Xue Li, and Michael G. Rosenfeld. Homeodomain-mediated β-catenin-dependent switching events dictate cell-lineage determination. Cell, 125(3): 593–605, 2006.

22. María Inés Pérez Millán, Michelle L Brinkmeier, Amanda H Mortensen, and Sally A Camper. Prop1 triggers epithelial-mesenchymal transition-like process in pituitary stem cells. eLife, 5, 2016.

23. Débora Cristina de Moraes, Mario Vaisman, Flavia Lucia Conceição, and Tânia Maria Ortiga-Carvalho. Pituitary development: a complex, temporal regulated process dependent on specific transcriptional factors. The Journal of endocrinology, 215(2): 239–45, 2012.

24. Julie L Youngblood, Tanner F Coleman, and Shannon W Davis. Regulation of Pituitary Progenitor Differentiation by β-Catenin. Endocrinology, 159(9): 3287–3305, 2018.

25. Yosuke Funato, Takeshi Terabayashi, Reiko Sakamoto, Daisuke Okuzaki, Hirotake Ichise, Hiroshi Nojima, Nobuaki Yoshida, and Hiroaki Miki. Nucleoredoxin sustains Wnt/β-catenin signaling by retaining a pool of inactive dishevelled protein. Current biology, 20 (21):1945–1952, 2010.

26. Robert A. Rigby, Mikis D. Stasinopoulos, Gillian Z. Heller, and Fernanda De Bastiani. Distributions for modeling location, scale, and shape: Using GAMLSS in R. Chapman and Hall/CRC, 2019.

27. Jing He, Hongzhe Li, Andrew C. Edmondson, Daniel J. Rader, and Mingyao Li. A gaussian copula approach for the analysis of secondary phenotypes in case–control genetic association studies. Biostatistics, 13(3): 497–508, 2012.

28. Abe Sklar. Fonctions de répartition à n dimensions et leurs marges. Publications de l’Institut de Statistique de l’Université de Paris, 8: 229–231, 1959.

29. Giampiero Marra and Rosalba Radice. Bivariate copula additive models for location, scale and shape. Computational Statistics & Data Analysis, 112: 99–113, 2017.

30. Rosalba Radice, Giampiero Marra, and Małgorzata Wojtyś. Copula regression spline models for binary outcomes. Statistics and Computing, 26(5): 981–995, 2016.

31. Giampiero Marra, Rosalba Radice, Till Bärnighausen, Simon N. Wood, and Mark E. McGovern. A simultaneous equation approach to estimating HIV prevalence with nonignorable missing responses. Journal of the American Statistical Association, 112(518):484–496, 2017.

32. Jorge Nocedal and Stephen J. Wright. Numerical optimization. New York, NY: Springer, 2006. ISBN New York, NY: Springer.

33. Peter K. Dunn and Gordon K. Smyth. Randomized quantile residuals. Journal of Computational and Graphical Statistics, 5(3): 236–244, 1996.

34. Kelly B. Cha, Kristin R. Douglas, Mary Anne Potok, Huiling Liang, Stephen N. Jones, and Sally A. Camper. Wnt5a signaling affects pituitary gland shape. Mechanisms of Development, 121(2): 183–194, 2004.

35. Julian Martinez-Mayer, Michelle L. Brinkmeier, Sean P. O’Connell, Arnold Ukagwu, Marcelo A. Marti, Mirta Miras, Maria V. Forclaz, Maria G. Benzrihen, Leonard Y. M. Cheung, Sally A. Camper, Buffy S. Ellsworth, Lori T. Raetzman, Maria I. Perez Millan, and Shannon W. Davis. Knockout mice with pituitary malformations help identify human cases of hypopituitarism. https://www.ncbi.nlm.nih.gov/geo/query/acc.cgi?acc=GSE246211, 2023. GSE246211, NCBI Gene Expression Ominbus.

36. Julian Martinez-Mayer, Michelle L. Brinkmeier, Sean P. O’Connell, Arnold Ukagwu, Marcelo A. Marti, Mirta Miras, Maria V. Forclaz, Maria G. Benzrihen, Leonard Y. M. Cheung, Sally A. Camper, Buffy S. Ellsworth, Lori T. Raetzman, Maria I. Perez Millan, and Shannon W. Davis. e12.5 pituitary gland from a retinoic acid reporter mouse (rare-lacz jax strain #008477). https://www.ncbi.nlm.nih.gov/geo/query/acc.cgi?acc=GSM7864906, 2023. GSM786906, NCBI Gene Expres-sion Omnibus.

37. Julian Martinez-Mayer, Michelle L. Brinkmeier, Sean P. O’Connell, Arnold Ukagwu, Marcelo A. Marti, Mirta Miras, Maria V. Forclaz, Maria G. Benzrihen, Leonard Y. M. Cheung, Sally A. Camper, Buffy S. Ellsworth, Lori T. Raetzman, Maria I. Perez Millan, and Shannon W. Davis. Two pooled e14.5 pituitary glands from c57bl6 strain. https://www.ncbi.nlm.nih.gov/geo/query/acc.cgi?acc=GSM7864907, 2023. GMS7864907, NCBI Gene Expression Omnibus.

