## Supplementary Data for "TIME-CoExpress: Temporal Trajectory Modeling of Dynamic Gene Co-expression Patterns Using Single-Cell Transcriptomics Data"

### A. Appendix

#### A.1. Marginal distribution

The marginal distributions can be set to a negative binomial distribution if the data is count-based, the PDF and CDF are given by Rigby et al. (2019),

$$f_{\mathrm{NB}}(w;\mu,\sigma) = \frac{\Gamma(w+\sigma^{-1})}{\Gamma(\sigma^{-1})\Gamma(w+1)}(\frac{\sigma\mu}{1+\sigma\mu})^{w}(\frac{1}{1+\sigma\mu})^{\frac{1}{\sigma}},$$

$$F_{\mathrm{NB}}(w;\mu,\sigma) = 1 - \frac{\mathrm{B}(w+1,\sigma^{-1},\mu\sigma(1+\mu\sigma)^{-1})}{\mathrm{B}(w+1,\sigma^{-1})},$$

where B() is the beta function.

#### A.2. Detailed model

This section provides the details of the proposed framework.

To incorporate gene zero-inflation characteristics into the model. Define $D_{ij} \sim \mathrm{Bern}\,(p_{ij})$, $j = 1, 2$, be the random variables representing whether gene $j$ gets zeroed out in cell $i$ due to dropout. The gene expression levels can be modeled as,

$$\begin{aligned} Y_{i1} &= (1-D_{i1})W_{i1}, \\ Y_{i2} &= (1-D_{i2})W_{i2}. \end{aligned}$$

The joint CDF is given by,

$$\begin{aligned} P\left(Y_{i1} \le y_{i1}, Y_{i2} \le y_{i2}\right) =& P\left(Y_{i1} \le y_{i1}, Y_{i2} \le y_{i2} \mid D_{i1}=1, D_{i2}=1\right) P(D_{i1}=1)P(D_{i2}=1) + \\ &+ P\left(Y_{i1} \le y_{i1}, Y_{i2} \le y_{i2} \mid D_{i1}=1, D_{i2}=0\right) P(D_{i1}=1)P(D_{i2}=0) + \\ &+ P\left(Y_{i1} \le y_{i1}, Y_{i2} \le y_{i2} \mid D_{i1}=0, D_{i2}=1\right) P(D_{i1}=0)P(D_{i2}=1) + \\ &+ P\left(Y_{i1} \le y_{i1}, Y_{i2} \le y_{i2} \mid D_{i1}=0, D_{i2}=0\right) P(D_{i1}=0)P(D_{i2}=0), \\ =& 1 \cdot p_{i1}p_{i2} + \\ &+ P\left(Y_{i2} \le y_{i2} \,|D_{i2}=0\right) p_{i1}(1-p_{i2}) + \\ &+ P\left(Y_{i1} \le y_{i1} \,|D_{i1}=0\right) (1-p_{i1})p_{i2} + \\ &+ P\left(Y_{i1} \le y_{i1}, Y_{i2} \le y_{i2} \mid D_{i1}=0, D_{i2}=0\right) (1-p_{i1})(1-p_{i2}), \\ =& p_{i1}p_{i2} + \\ &+ P\left(W_{i2} \le y_{i2}\right) p_{i1}(1-p_{i2}) + \\ &+ P\left(W_{i1} \le y_{i1}\right) (1-p_{i1})p_{i2} + \\ &+ P\left(W_{i1} \le y_{i1}, W_{i2} \le y_{i2}\right) (1-p_{i1})(1-p_{i2}), \end{aligned}$$

which can be simplified as,

$$P\left(Y_{i1} \le y_{i1}, Y_{i2} \le y_{i2}\right) = \begin{cases} p_{i1}p_{i2} & \text{if } y_{i1}=0 \text{ and } y_{i2}=0, \\ p_{i1}p_{i2} + F_{\mathrm{GA}}\left(y_{i1};\mu_{i1},\sigma_{i1}\right)(1-p_{i1})p_{i2} & \text{if } y_{i1}>0 \text{ and } y_{i2}=0, \\ p_{i1}p_{i2} + F_{\mathrm{GA}}\left(y_{i2};\mu_{i2},\sigma_{i2}\right)p_{i1}(1-p_{i2}) & \text{if } y_{i1}=0 \text{ and } y_{i2}>0, \\ p_{i1}p_{i2} + F_{\boldsymbol{w}_i}\left(y_{i1},y_{i2};\ \mu_{i1},\mu_{i2},\sigma_{i1},\sigma_{i2},\rho_i\right)(1-p_{i1})(1-p_{i2}) & \text{if } y_{i1}>0 \text{ and } y_{i2}>0. \end{cases} \tag{A.1}$$

The log-likelihood can be expressed as,

$$\begin{aligned} &\ell\left(\mu_{i1},\mu_{i2},\sigma_{i1},\sigma_{i2},\rho_i,p_{i1},p_{i2};\ y_{i1},y_{i2}\right) = \\ &\begin{cases} \log(p_{i1}) + \log(p_{i2}) & \text{if } y_{i1}=0 \text{ and } y_{i2}=0, \\ \log\left(f_{\mathrm{GA}}\left(y_{i1};\mu_{i1},\sigma_{i1}\right)\right) + \log\left(1-p_{i1}\right) + \log\left(p_{i2}\right) & \text{if } y_{i1}>0 \text{ and } y_{i2}=0, \\ \log\left(f_{\mathrm{GA}}\left(y_{i2};\mu_{i2},\sigma_{i2}\right)\right) + \log\left(p_{i1}\right) + \log\left(1-p_{i2}\right) & \text{if } y_{i1}=0 \text{ and } y_{i2}>0, \\ \log\left(f_{W_{i1},W_{i2}}\left(y_{i1},y_{i2};\ \mu_{i1},\mu_{i2},\sigma_{i1},\sigma_{i2},\rho_i\right)\right) + \log\left(1-p_{i1}\right) + \log\left(1-p_{i2}\right) & \text{if } y_{i1}>0 \text{ and } y_{i2}>0, \end{cases} \end{aligned}$$

where $p_{i1}, p_{i2}$ appear solely as additive constants in the log-likelihood function, which means the estimation of parameters $\mu_{i1}, \mu_{i2}, \sigma_{i1}, \sigma_{i2}, \rho_i$ does not depend on the values of $p_{i1}, p_{i2}$. So we can simplify the above log-likelihood to,

$$\ell_i = \ell\left(\mu_{i1}, \mu_{i2}, \sigma_{i1}, \sigma_{i2}, \rho_i;\ y_{i1}, y_{i2}\right) = k_i + \begin{cases} 0 & \text{if } y_{i1} = 0 \text{ and } y_{i2} = 0, \\ \log\left(f_{\text{GA}}\left(y_{i1}; \mu_{i1}, \sigma_{i1}\right)\right) & \text{if } y_{i1} > 0 \text{ and } y_{i2} = 0, \\ \log\left(f_{\text{GA}}\left(y_{i2}; \mu_{i2}, \sigma_{i2}\right)\right) & \text{if } y_{i1} = 0 \text{ and } y_{i2} > 0, \\ \log\left(f_{W_{i1}, W_{i2}}\left(y_{i1}, y_{i2};\ \mu_{i1}, \mu_{i2}, \sigma_{i1}, \sigma_{i2}, \rho_i\right)\right) & \text{if } y_{i1} > 0 \text{ and } y_{i2} > 0, \end{cases}$$

and carry out the estimation of $p_{i1}, p_{i2}$ in a separate step.

In our model, there are seven unknown parameters $\boldsymbol{\vartheta} = (p_1, p_2, \mu_1, \sigma_1, \mu_2, \sigma_2, \rho)^T$, where $p_1$ and $p_2$ are the zero-inflation rates of each gene; $(\mu_1, \sigma_1)$ and $(\mu_2, \sigma_2)$ are the marginal distribution parameters for two marginals respectively; $\rho$ is the copula dependence parameter. The two covariates $\mathbf{Z} = (\boldsymbol{z_1}, \boldsymbol{z_2})^T$ are cell pseudotime and an indicator variable with level 0 and 1 to identify the control group and the mutant group, where $\boldsymbol{z_1} = (z_{11}, ..., z_{1N})^T$, $\boldsymbol{z_2} = (z_{21}, ..., z_{2N})^T$, $N$ is the number of cells. We use logit as the monotonic link function for zero-inflation rates $p_1$ and $p_2$, use log() for distribution parameters to transfer positive $\mu$ and $\sigma$ to the range from negative to positive infinity, and use inverse tangent function to transfer copula dependence parameter $\rho$. Assume the zero-inflation rates, standard deviation, mean and correlation parameters have non-linear relationships with these two covariates. We use the thin plate regression splines to construct additive predictors. Each parameter is modeled by its additive predictors denoted as,

$$\begin{aligned}
\eta_{p_{i1}} = \text{logit}(p_{i1}) = \beta_{p_1} &+ \sum_{j=1}^{J} \beta_{p_1, z_1, z_2=1, j} b_{p_1, z_1, z_2=1, j}(z_{i1}) \mathbb{1}(z_{i2} = 1), \\
&+ \sum_{j=1}^{J} \beta_{p_1, z_1, z_2=0, j} b_{p_1, z_1, z_2=0, j}(z_{i1}) \mathbb{1}(z_{i2} = 0), \\
\eta_{p_{i2}} = \text{logit}(p_{i2}) = \beta_{p_2} &+ \sum_{j=1}^{J} \beta_{p_2, z_1, z_2=1, j} b_{p_2, z_1, z_2=1, j}(z_{i1}) \mathbb{1}(z_{i2} = 1), \\
&+ \sum_{j=1}^{J} \beta_{p_2, z_1, z_2=0, j} b_{p_2, z_1, z_2=0, j}(z_{i1}) \mathbb{1}(z_{i2} = 0), \\
\eta_{\mu_{i1}} = \log(\mu_{i1}) = \beta_{\mu_1} &+ \sum_{j=1}^{J} \beta_{\mu_1, z_1, z_2=1, j} b_{\mu_1, z_1, z_2=1, j}(z_{i1}) \mathbb{1}(z_{i2} = 1), \\
&+ \sum_{j=1}^{J} \beta_{\mu_1, z_1, z_2=0, j} b_{\mu_1, z_1, z_2=0, j}(z_{i1}) \mathbb{1}(z_{i2} = 0), \\
\eta_{\mu_{i2}} = \log(\mu_{i2}) = \beta_{\mu_2} &+ \sum_{j=1}^{J} \beta_{\mu_2, z_1, z_2=1, j} b_{\mu_2, z_1, z_2=1, j}(z_{i1}) \mathbb{1}(z_{i2} = 1), \\
&+ \sum_{j=1}^{J} \beta_{\mu_2, z_1, z_2=0, j} b_{\mu_2, z_1, z_2=0, j}(z_{i1}) \mathbb{1}(z_{i2} = 0), \\
\eta_{\sigma_{i1}} = \log(\sigma_{i1}) = \beta_{\sigma_1} &+ \sum_{j=1}^{J} \beta_{\sigma_1, z_1, z_2=1, j} b_{\sigma_1, z_1, z_2=1, j}(z_{i1}) \mathbb{1}(z_{i2} = 1), \\
&+ \sum_{j=1}^{J} \beta_{\sigma_1, z_1, z_2=0, j} b_{\sigma_1, z_1, z_2=0, j}(z_{i1}) \mathbb{1}(z_{i2} = 0), \\
\eta_{\sigma_{i2}} = \log(\sigma_{i2}) = \beta_{\sigma_2} &+ \sum_{j=1}^{J} \beta_{\sigma_2, z_1, z_2=1, j} b_{\sigma_2, z_1, z_2=1, j}(z_{i1}) \mathbb{1}(z_{i2} = 1), \\
&+ \sum_{j=1}^{J} \beta_{\sigma_2, z_1, z_2=0, j} b_{\sigma_2, z_1, z_2=0, j}(z_{i1}) \mathbb{1}(z_{i2} = 0), \\
\eta_{\rho_i} = \text{atanh}(\rho_i) = \beta_{\rho} &+ \sum_{j=1}^{J} \beta_{\rho, z_1, z_2=1, j} b_{\rho, z_1, z_2=1, j}(z_{i1}) \mathbb{1}(z_{i2} = 1), \\
&+ \sum_{j=1}^{J} \beta_{\rho, z_1, z_2=0, j} b_{\rho, z_1, z_2=0, j}(z_{i1}) \mathbb{1}(z_{i2} = 0),
\end{aligned}$$

where $\beta_\theta$ is the intercept, $\beta_{\theta, z_1, z_2}$ is denoted as the regression coefficients to be estimated; $b(\cdot)$ is the $j$th basis function of the spline pertaining to that particular parameter and covariate.

The parameters of our model can also be expressed in the following way:

$$\mu_{i1} = \exp\left\{\boldsymbol{\beta}_1^\top \begin{bmatrix} 1 \\ \boldsymbol{b}^{(\mu_1)}(\boldsymbol{z}_i) \end{bmatrix}\right\},$$

$$\mu_{i2} = \exp\left\{\boldsymbol{\beta}_2^\top \begin{bmatrix} 1 \\ \boldsymbol{b}^{(\mu_2)}(\boldsymbol{z}_i) \end{bmatrix}\right\},$$

$$\sigma_{i1} = \exp\left\{\boldsymbol{\alpha}_1^\top \begin{bmatrix} 1 \\ \boldsymbol{b}^{(\sigma_1)}(\boldsymbol{z}_i) \end{bmatrix}\right\},$$

$$\sigma_{i2} = \exp\left\{\boldsymbol{\alpha}_2^\top \begin{bmatrix} 1 \\ \boldsymbol{b}^{(\sigma_2)}(\boldsymbol{z}_i) \end{bmatrix}\right\},$$

$$\rho_i = \tanh\left\{\boldsymbol{\tau}^\top \begin{bmatrix} 1 \\ \boldsymbol{b}^{(\rho)}(\boldsymbol{z}_i) \end{bmatrix}\right\},$$

$$p_{i1} = \text{sigmoid}\left\{\boldsymbol{\kappa}_1^\top \begin{bmatrix} 1 \\ \boldsymbol{b}^{(p_1)}(\boldsymbol{z}_i) \end{bmatrix}\right\},$$

$$p_{i2} = \text{sigmoid}\left\{\boldsymbol{\kappa}_2^\top \begin{bmatrix} 1 \\ \boldsymbol{b}^{(p_2)}(\boldsymbol{z}_i) \end{bmatrix}\right\},$$

where $\boldsymbol{b}^{(\text{param})}(\boldsymbol{z}_i)$ is the vector of basis functions corresponding to (param) evaluated at their respective subvectors of $\boldsymbol{z}_i$. The coefficients to be estimated in the model are thus $\boldsymbol{\beta}_1, \boldsymbol{\beta}_2, \boldsymbol{\alpha}_1, \boldsymbol{\alpha}_2, \boldsymbol{\tau}, \boldsymbol{\kappa}_1, \boldsymbol{\kappa}_2$.

The following result,

$$\begin{aligned} p_{ji} &= P(D_{ji} = 1), \\ &= P(D_{ji} = 1 \,|Y_{ji} > 0\,)P(Y_{ji} > 0) + P(D_{ji} = 1 \,|Y_{ji} = 0\,)P(Y_{ji} = 0), \\ &= 0 \cdot P(Y_{ji} > 0) + 1 \cdot P(Y_{ji} = 0), \\ &= P(Y_{ji} = 0), \end{aligned}$$

allows us to estimate $\boldsymbol{\kappa}_1, \boldsymbol{\kappa}_2$ using logistic regression on $\mathbf{1}\{Y_{ji} = 0\}$, while $\boldsymbol{\beta}_1, \boldsymbol{\beta}_2, \boldsymbol{\alpha}_1, \boldsymbol{\alpha}_2, \boldsymbol{\tau}$ are estimated by maximizing $\sum_{i=1}^n \ell_i$. This maximization is achieved using the trust algorithm as detailed in section 2.2.4, which requires the gradient and hessian to be explicitly calculated.

#### A.3. Gradient

This section provides the gradient of log-likelihood. Taking the derivative of $\ell_p$ with respect to $\boldsymbol{\delta}$ yields:

$$\begin{aligned} \frac{\partial}{\partial \boldsymbol{\delta}} \ell_p &= \frac{\partial}{\partial \boldsymbol{\delta}} \left( \sum \ell_i - \frac{1}{2} \boldsymbol{\delta}^\top \boldsymbol{S} \boldsymbol{\delta} \right), \\ &= \sum \frac{\partial}{\partial \boldsymbol{\delta}} \ell_i - \boldsymbol{S}\boldsymbol{\delta}. \end{aligned}$$

So we turn our focus to calculating $\frac{\partial}{\partial \boldsymbol{\delta}} \ell_i$. For notational simplicity, we will drop the subscript $i$ for the remainder of this document. We will also let

$$\begin{aligned} u_1 &= F_{\text{GA}}(y_1;\ \mu_1, \sigma_1), \\ u_2 &= F_{\text{GA}}(y_2;\ \mu_2, \sigma_2), \\ u_1' &= f_{\text{GA}}(y_1;\ \mu_1, \sigma_1), \\ u_2' &= f_{\text{GA}}(y_2;\ \mu_2, \sigma_2). \end{aligned}$$

which means the log-likelihood $\ell$ can be written:

$$\ell = k + \begin{cases} 0 & \text{if } y_{i1} = 0 \text{ and } y_{i2} = 0, \\ \log(u_1') & \text{if } y_{i1} > 0 \text{ and } y_{i2} = 0, \\ \log(u_2') & \text{if } y_{i1} = 0 \text{ and } y_{i2} > 0, \\ \log(c(u_1, u_2; \rho)) + \log(u_1') + \log(u_2') & \text{if } y_{i1} > 0 \text{ and } y_{i2} > 0. \end{cases}$$

So in order to calculate the first derivatives of $\ell$ with respect to $\boldsymbol{\beta}_1, \boldsymbol{\beta}_2, \boldsymbol{\alpha}_1, \boldsymbol{\alpha}_2, \boldsymbol{\tau}$, we will need to first calculate the derivatives of $u_1', u_2', u_1, u_2, c(u_1, u_2; \rho)$ with respect to all those coefficients. The first step of that process will be to calculate $\frac{\partial \mu_1}{\partial \boldsymbol{\beta}_1}, \frac{\partial \mu_2}{\partial \boldsymbol{\beta}_2}, \frac{\partial \sigma_1}{\partial \boldsymbol{\alpha}_1}, \frac{\partial \sigma_2}{\partial \boldsymbol{\alpha}_2}, \frac{\partial \rho}{\partial \boldsymbol{\tau}}$, so we will get that out of the way first:

$$\frac{\partial \mu_1}{\partial \boldsymbol{\beta}_1} = \exp\left\{\boldsymbol{\beta}_1^\top \begin{bmatrix} 1, \\ \boldsymbol{b}^{(\mu_1)}(\boldsymbol{x}) \end{bmatrix}\right\} \begin{bmatrix} 1, \\ \boldsymbol{b}^{(\mu_1)}(\boldsymbol{x}) \end{bmatrix},$$

$$\frac{\partial \mu_2}{\partial \boldsymbol{\beta}_2} = \exp\left\{\boldsymbol{\beta}_2^\top \begin{bmatrix} 1, \\ \boldsymbol{b}^{(\mu_2)}(\boldsymbol{x}) \end{bmatrix}\right\} \begin{bmatrix} 1, \\ \boldsymbol{b}^{(\mu_2)}(\boldsymbol{x}) \end{bmatrix},$$

$$\frac{\partial \sigma_1}{\partial \boldsymbol{\alpha}_1} = \exp\left\{\boldsymbol{\alpha}_1^\top \begin{bmatrix} 1, \\ \boldsymbol{b}^{(\sigma_1)}(\boldsymbol{x}) \end{bmatrix}\right\} \begin{bmatrix} 1, \\ \boldsymbol{b}^{(\sigma_1)}(\boldsymbol{x}) \end{bmatrix},$$

$$\frac{\partial \sigma_2}{\partial \boldsymbol{\alpha}_2} = \exp\left\{\boldsymbol{\alpha}_2^\top \begin{bmatrix} 1, \\ \boldsymbol{b}^{(\sigma_2)}(\boldsymbol{x}) \end{bmatrix}\right\} \begin{bmatrix} 1, \\ \boldsymbol{b}^{(\sigma_2)}(\boldsymbol{x}) \end{bmatrix},$$

$$\frac{\partial \rho}{\partial \boldsymbol{\tau}} = \cosh^2\left\{\boldsymbol{\tau}^\top \begin{bmatrix} 1, \\ \boldsymbol{b}^{(\rho)}(\boldsymbol{x}) \end{bmatrix}\right\} \begin{bmatrix} 1, \\ \boldsymbol{b}^{(\rho)}(\boldsymbol{x}) \end{bmatrix}.$$

Now we can calculate the aforementioned derivatives of $u_1', u_2', u_1, u_2, c(u_1, u_2; \rho)$:

$$\frac{\partial u_j'}{\partial \boldsymbol{\beta}_j} = \frac{\partial f_{\mathrm{GA}}(y_j;\ \mu_j, \sigma_j)}{\partial \boldsymbol{\beta}_j} = \frac{\partial f_{\mathrm{GA}}(y_j;\ \mu_j, \sigma_j)}{\partial \mu_j}\frac{\partial \mu_j}{\partial \boldsymbol{\beta}_j} = f_{\mathrm{GA}}(y_j;\ \mu_j, \sigma_j)\left[\frac{-1}{\mu_j} + \frac{y_j}{\mu_j^2 \sigma_j^2}\right]\frac{\partial \mu_j}{\partial \boldsymbol{\beta}_j}.$$

$$\frac{\partial u_j}{\partial \boldsymbol{\beta}_j} = \frac{\partial \int_0^{y_j} f_{\mathrm{GA}}(t;\ \mu_j, \sigma_j)\,dt}{\partial \boldsymbol{\beta}_j} = \int_0^{y_j} \frac{\partial f_{\mathrm{GA}}(t;\ \mu_j, \sigma_j)}{\partial \mu_j}\frac{\partial \mu_j}{\partial \boldsymbol{\beta}_j} dt = \frac{\partial \mu_j}{\partial \boldsymbol{\beta}_j}\int_0^{y_j} f_{\mathrm{GA}}(t;\ \mu_j, \sigma_j)\left[\frac{-1}{\mu_j} + \frac{t}{\mu_j^2 \sigma_j^2}\right] dt.$$

$$\begin{aligned}\frac{\partial u_j'}{\partial \boldsymbol{\alpha}_j} &= \frac{\partial f_{\mathrm{GA}}(y_j;\ \mu_j, \sigma_j)}{\partial \boldsymbol{\alpha}_j} = \frac{\partial f_{\mathrm{GA}}(y_j;\ \mu_j, \sigma_j)}{\partial \sigma_j}\frac{\partial \sigma_j}{\partial \boldsymbol{\alpha}_j}, \\ &= f_{\mathrm{GA}}(y_j;\ \mu_j, \sigma_j)\left[\frac{2}{\sigma_j^3}\log(\mu_j \sigma_j^2) - \frac{1}{\sigma_j^3} + \frac{2}{\sigma_j^3}\psi\left(\frac{1}{\sigma_j^2}\right) - \frac{2\log y_j}{\sigma_j^3} + \frac{2y_j}{\mu_j \sigma_j^3}\right]\frac{\partial \sigma_j}{\partial \boldsymbol{\alpha}_j}.\end{aligned}$$

$$\begin{aligned}\frac{\partial u_j}{\partial \boldsymbol{\alpha}_j} &= \frac{\partial \int_0^{y_j} f_{\mathrm{GA}}(t;\ \mu_j, \sigma_j)\,dt}{\partial \boldsymbol{\alpha}_j} = \int_0^{y_j} \frac{\partial f_{\mathrm{GA}}(t;\ \mu_j, \sigma_j)}{\partial \sigma_j}\frac{\partial \sigma_j}{\partial \boldsymbol{\alpha}_j} dt, \\ &= \frac{\partial \sigma_j}{\partial \boldsymbol{\alpha}_j}\int_0^{y_j} f_{\mathrm{GA}}(t;\ \mu_j, \sigma_j)\left[\frac{2}{\sigma_j^3}\log(\mu_j \sigma_j^2) - \frac{1}{\sigma_j^3} + \frac{2}{\sigma_j^3}\psi\left(\frac{1}{\sigma_j^2}\right) - \frac{2\log y_j}{\sigma_j^3} + \frac{2y_j}{\mu_j \sigma_j^3}\right] dt.\end{aligned}$$

$$\begin{aligned}\frac{\partial c(u_1, u_2; \rho)}{\partial \boldsymbol{\tau}} &= \frac{\partial c(u_1, u_2; \rho)}{\partial \rho}\frac{\partial \rho}{\partial \boldsymbol{\tau}}, \\ &= -(1-\rho^2)^{\frac{-5}{2}}\left(\rho^3 - \Phi^{-1}(u_1)\Phi^{-1}(u_2)\rho^2 + \rho\Phi^{-1}(u_2)^2 + \rho\Phi^{-1}(u_1)^2 - \rho - \Phi^{-1}(u_1)\Phi^{-1}(u_2)\right) \times, \\ &\quad \times \exp\left\{\frac{1}{2}\frac{\rho(\Phi^{-1}(u_1)^2 + \Phi^{-1}(u_2)^2 - 2x_1\Phi^{-1}(u_2))}{(\rho-1)(\rho+1)}\right\}\frac{\partial \rho}{\partial \boldsymbol{\tau}}.\end{aligned}$$

$$\begin{aligned}\frac{\partial c(u_1, u_2; \rho)}{\partial \boldsymbol{\beta}_1} &= \frac{\partial c(u_1, u_2; \rho)}{\partial u_1}\frac{\partial u_1}{\partial \boldsymbol{\beta}_1}, \\ &= c(u_1, u_2; \rho)\left(-\frac{2\rho^2\Phi^{-1}(u_1)\left(\frac{1}{\phi\{\Phi^{-1}(u_1)\}}\right) - 2\rho\Phi^{-1}(u_2)\left(\frac{1}{\phi\{\Phi^{-1}(u_1)\}}\right)}{2(1-\rho^2)}\right)\frac{\partial u_1}{\partial \boldsymbol{\beta}_1}.\end{aligned}$$

$$\begin{aligned}\frac{\partial c(u_1, u_2; \rho)}{\partial \boldsymbol{\beta}_2} &= \frac{\partial c(u_1, u_2; \rho)}{\partial u_2}\frac{\partial u_2}{\partial \boldsymbol{\beta}_2}, \\ &= c(u_1, u_2; \rho)\left(-\frac{2\rho^2\Phi^{-1}(u_2)\left(\frac{1}{\phi\{\Phi^{-1}(u_2)\}}\right) - 2\rho\Phi^{-1}(u_1)\left(\frac{1}{\phi\{\Phi^{-1}(u_2)\}}\right)}{2(1-\rho^2)}\right)\frac{\partial u_2}{\partial \boldsymbol{\beta}_2}.\end{aligned}$$

$$\frac{\partial c(u_1, u_2; \rho)}{\partial \boldsymbol{\alpha}_j} = \frac{\partial c(u_1, u_2; \rho)}{\partial u_j}\frac{\partial u_j}{\partial \boldsymbol{\alpha}_j}.$$

With these derivatives done, it will be easy to calculate the gradient of our log-likelihood $\ell$.

##### A.3.1. Derivative of log-likelihood with respect to $\boldsymbol{\beta}_1$

$$\frac{\partial \ell}{\partial \boldsymbol{\beta}_1} = \frac{1}{\mathcal{L}} \begin{cases} 0 & \text{if } y_1 = 0 \text{ and } y_2 = 0, \\ \frac{\partial u_1'}{\partial \boldsymbol{\beta}_1} & \text{if } y_1 > 0 \text{ and } y_2 = 0, \\ 0 & \text{if } y_1 = 0 \text{ and } y_2 > 0, \\ \frac{\partial c(u_1,u_2;\rho)}{\partial \boldsymbol{\beta}_1} u_1' u_2' + c(u_1,u_2;\rho) \frac{\partial u_1'}{\partial \boldsymbol{\beta}_1} u_2' & \text{if } y_1 > 0 \text{ and } y_2 > 0. \end{cases}$$

##### A.3.2. Derivative of log-likelihood with respect to $\boldsymbol{\beta}_2$

$$\frac{\partial \ell}{\partial \boldsymbol{\beta}_2} = \frac{1}{\mathcal{L}} \begin{cases} 0 & \text{if } y_1 = 0 \text{ and } y_2 = 0, \\ 0 & \text{if } y_1 > 0 \text{ and } y_2 = 0, \\ \frac{\partial u_2'}{\partial \boldsymbol{\beta}_2} & \text{if } y_1 = 0 \text{ and } y_2 > 0, \\ \frac{\partial c(u_1,u_2;\rho)}{\partial \boldsymbol{\beta}_2} u_1' u_2' + c(u_1,u_2;\rho) u_1' \frac{\partial u_2'}{\partial \boldsymbol{\beta}_2} & \text{if } y_1 > 0 \text{ and } y_2 > 0. \end{cases}$$

##### A.3.3. Derivative of log-likelihood with respect to $\boldsymbol{\alpha}_1$

$$\frac{\partial \ell}{\partial \boldsymbol{\alpha}_1} = \frac{1}{\mathcal{L}} \begin{cases} 0 & \text{if } y_1 = 0 \text{ and } y_2 = 0, \\ \frac{\partial u_1'}{\partial \boldsymbol{\alpha}_1} & \text{if } y_1 > 0 \text{ and } y_2 = 0, \\ 0 & \text{if } y_1 = 0 \text{ and } y_2 > 0, \\ \frac{\partial c(u_1,u_2;\rho)}{\partial \boldsymbol{\alpha}_1} u_1' u_2' + c(u_1,u_2;\rho) \frac{\partial u_1'}{\partial \boldsymbol{\alpha}_1} u_2' & \text{if } y_1 > 0 \text{ and } y_2 > 0. \end{cases}$$

##### A.3.4. Derivative of log-likelihood with respect to $\boldsymbol{\alpha}_2$

$$\frac{\partial \ell}{\partial \boldsymbol{\alpha}_2} = \frac{1}{\mathcal{L}} \begin{cases} 0 & \text{if } y_1 = 0 \text{ and } y_2 = 0, \\ 0 & \text{if } y_1 > 0 \text{ and } y_2 = 0, \\ \frac{\partial u_2'}{\partial \boldsymbol{\alpha}_2} & \text{if } y_1 = 0 \text{ and } y_2 > 0, \\ \frac{\partial c(u_1,u_2;\rho)}{\partial \boldsymbol{\alpha}_2} u_1' u_2' + c(u_1,u_2;\rho) u_1' \frac{\partial u_2'}{\partial \boldsymbol{\alpha}_2} & \text{if } y_1 > 0 \text{ and } y_2 > 0. \end{cases}$$

##### A.3.5. Derivative of log-likelihood with respect to $\boldsymbol{\tau}$

$$\frac{\partial \ell}{\partial \boldsymbol{\tau}} = \frac{1}{\mathcal{L}} \begin{cases} 0 & \text{if } y_1 = 0 \text{ and } y_2 = 0, \\ 0 & \text{if } y_1 > 0 \text{ and } y_2 = 0, \\ 0 & \text{if } y_1 = 0 \text{ and } y_2 > 0, \\ \frac{\partial c(u_1,u_2;\rho)}{\partial \boldsymbol{\tau}} u_1' u_2' & \text{if } y_1 > 0 \text{ and } y_2 > 0. \end{cases}$$

#### A.4. Hessian

This section provides the Hessian of log-likelihood. Taking the second derivative of $\ell_p$ with respect to $\boldsymbol{\delta}$ yields:

$$\begin{aligned} \frac{\partial^2}{\partial \boldsymbol{\delta}^2} \ell_p &= \frac{\partial^2}{\partial \boldsymbol{\delta}^2} \left( \sum \ell_i - \frac{1}{2} \boldsymbol{\delta}^\top \boldsymbol{S} \boldsymbol{\delta} \right), \\ &= \sum \frac{\partial^2}{\partial \boldsymbol{\delta}^2} \ell_i - \boldsymbol{S}. \end{aligned}$$

So we turn our focus to calculating $\frac{\partial^2}{\partial \boldsymbol{\delta}^2} \ell_i$. To calculate the second derivatives of $\ell$ with respect to $\boldsymbol{\beta}_1, \boldsymbol{\beta}_2, \boldsymbol{\alpha}_1, \boldsymbol{\alpha}_2, \boldsymbol{\tau}$, we will need to calculate the second derivatives of $u_1', u_2', u_1, u_2, c(u_1,u_2;\rho)$ with respect to all those coefficients. First, we will calculate some of the basic components that will appear in the chain rule calculations throughout:

$$\begin{aligned} \frac{\partial^2 \mu_1}{\partial \beta_1^2} &= \exp\left\{ \boldsymbol{\beta}_1^\top \begin{bmatrix} 1 \\ \boldsymbol{b}^{(\mu_1)}(\boldsymbol{x}) \end{bmatrix} \right\} \begin{bmatrix} 1 \\ \boldsymbol{b}^{(\mu_1)}(\boldsymbol{x}) \end{bmatrix} \begin{bmatrix} 1 \\ \boldsymbol{b}^{(\mu_1)}(\boldsymbol{x}) \end{bmatrix}^\top, \\ \frac{\partial \mu_2}{\partial \boldsymbol{\beta}_2} &= \exp\left\{ \boldsymbol{\beta}_2^\top \begin{bmatrix} 1 \\ \boldsymbol{b}^{(\mu_2)}(\boldsymbol{x}) \end{bmatrix} \right\} \begin{bmatrix} 1 \\ \boldsymbol{b}^{(\mu_2)}(\boldsymbol{x}) \end{bmatrix} \begin{bmatrix} 1 \\ \boldsymbol{b}^{(\mu_2)}(\boldsymbol{x}) \end{bmatrix}^\top, \\ \frac{\partial \sigma_1}{\partial \boldsymbol{\alpha}_1} &= \exp\left\{ \boldsymbol{\alpha}_1^\top \begin{bmatrix} 1 \\ \boldsymbol{b}^{(\sigma_1)}(\boldsymbol{x}) \end{bmatrix} \right\} \begin{bmatrix} 1 \\ \boldsymbol{b}^{(\sigma_1)}(\boldsymbol{x}) \end{bmatrix} \begin{bmatrix} 1 \\ \boldsymbol{b}^{(\sigma_1)}(\boldsymbol{x}) \end{bmatrix}^\top, \\ \frac{\partial \sigma_2}{\partial \boldsymbol{\alpha}_2} &= \exp\left\{ \boldsymbol{\alpha}_2^\top \begin{bmatrix} 1 \\ \boldsymbol{b}^{(\sigma_2)}(\boldsymbol{x}) \end{bmatrix} \right\} \begin{bmatrix} 1 \\ \boldsymbol{b}^{(\sigma_2)}(\boldsymbol{x}) \end{bmatrix} \begin{bmatrix} 1 \\ \boldsymbol{b}^{(\sigma_2)}(\boldsymbol{x}) \end{bmatrix}^\top, \\ \frac{\partial \rho}{\partial \boldsymbol{\tau}} &= -2\text{sech}^2 \left\{ \boldsymbol{\tau}^\top \begin{bmatrix} 1 \\ \boldsymbol{b}^{(\rho)}(\boldsymbol{x}) \end{bmatrix} \right\} \tanh \left\{ \boldsymbol{\tau}^\top \begin{bmatrix} 1 \\ \boldsymbol{b}^{(\rho)}(\boldsymbol{x}) \end{bmatrix} \right\} \begin{bmatrix} 1 \\ \boldsymbol{b}^{(\rho)}(\boldsymbol{x}) \end{bmatrix} \begin{bmatrix} 1 \\ \boldsymbol{b}^{(\rho)}(\boldsymbol{x}) \end{bmatrix}^\top. \end{aligned}$$

Now we can calculate the aforementioned second derivatives of $u_1', u_2', u_1, u_2, c(u_1,u_2;\rho)$:

$$\frac{\partial^2 u_j'}{\partial \boldsymbol{\beta}_j^2} = \frac{\partial}{\partial \boldsymbol{\beta}_j}\left[\frac{\partial f_{\mathrm{GA}}\left(y_j;\ \mu_j,\sigma_j\right)}{\partial \boldsymbol{\beta}_j}\right] = \frac{\partial}{\partial \boldsymbol{\beta}_j}\left[\frac{\partial f_{\mathrm{GA}}\left(y_j;\ \mu_j,\sigma_j\right)}{\partial \mu_j}\frac{\partial \mu_j}{\partial \boldsymbol{\beta}_j}\right],$$
$$= \left(\frac{\partial}{\partial \boldsymbol{\beta}_j}\frac{\partial f_{\mathrm{GA}}\left(y_j;\ \mu_j,\sigma_j\right)}{\partial \mu_j}\right)\frac{\partial \mu_j}{\partial \boldsymbol{\beta}_j} + \frac{\partial f_{\mathrm{GA}}\left(y_j;\ \mu_j,\sigma_j\right)}{\partial \mu_j}\left(\frac{\partial}{\partial \boldsymbol{\beta}_j}\frac{\partial \mu_j}{\partial \boldsymbol{\beta}_j}\right),$$
$$= \frac{\partial^2 f_{\mathrm{GA}}\left(y_j;\ \mu_j,\sigma_j\right)}{\partial \mu_j^2}\left[\frac{\partial \mu_j}{\partial \boldsymbol{\beta}_j}\right]^2 + \frac{\partial f_{\mathrm{GA}}\left(y_j;\ \mu_j,\sigma_j\right)}{\partial \mu_j}\frac{\partial^2 \mu_j}{\partial \boldsymbol{\beta}_j^2},$$
$$= f_{\mathrm{GA}}(y_j;\mu_j,\sigma_j)\left[\frac{2}{\mu_j^2} - \frac{4y_j}{\mu_j^3\sigma_j^2} + \frac{y_j^2}{\mu_j^4\sigma_j^4}\right]\left[\frac{\partial \mu_j}{\partial \boldsymbol{\beta}_j}\right]^2 + \frac{\partial f_{\mathrm{GA}}\left(y_j;\ \mu_j,\sigma_j\right)}{\partial \mu_j}\frac{\partial^2 \mu_j}{\partial \boldsymbol{\beta}_j^2}.$$

$$\frac{\partial^2 u_j}{\partial \boldsymbol{\beta}_j^2} = \left[\frac{\partial \mu_j}{\partial \boldsymbol{\beta}_j}\right]^2 \int_0^{y_j} f_{\mathrm{GA}}(t;\mu_j,\sigma_j)\left[\frac{2}{\mu_j^2} - \frac{4t}{\mu_j^3\sigma_j^2} + \frac{t^2}{\mu_j^4\sigma_j^4}\right]dt + \frac{\partial^2 \mu_j}{\partial \boldsymbol{\beta}_j^2}\int_0^{y_j}\frac{\partial f_{\mathrm{GA}}\left(t;\ \mu_j,\sigma_j\right)}{\partial \mu_j}dt.$$

$$\frac{\partial^2 u_j'}{\partial \boldsymbol{\alpha}_j^2} = \frac{\partial}{\partial \boldsymbol{\alpha}_j}\left[\frac{\partial f_{\mathrm{GA}}\left(y_j;\ \mu_j,\sigma_j\right)}{\partial \boldsymbol{\alpha}_j}\right] = \frac{\partial}{\partial \boldsymbol{\alpha}_j}\left[\frac{\partial f_{\mathrm{GA}}\left(y_j;\ \mu_j,\sigma_j\right)}{\partial \sigma_j}\frac{\partial \sigma_j}{\partial \boldsymbol{\alpha}_j}\right],$$
$$= \left(\frac{\partial}{\partial \boldsymbol{\alpha}_j}\frac{\partial f_{\mathrm{GA}}\left(y_j;\ \mu_j,\sigma_j\right)}{\partial \sigma_j}\right)\frac{\partial \sigma_j}{\partial \boldsymbol{\alpha}_j} + \frac{\partial f_{\mathrm{GA}}\left(y_j;\ \mu_j,\sigma_j\right)}{\partial \sigma_j}\left(\frac{\partial}{\partial \boldsymbol{\alpha}_j}\frac{\partial \sigma_j}{\partial \boldsymbol{\alpha}_j}\right),$$
$$= \frac{\partial^2 f_{\mathrm{GA}}\left(y_j;\ \mu_j,\sigma_j\right)}{\partial \sigma_j^2}\left[\frac{\partial \sigma_j}{\partial \boldsymbol{\alpha}_j}\right]^2 + \frac{\partial f_{\mathrm{GA}}\left(y_j;\ \mu_j,\sigma_j\right)}{\partial \sigma_j}\frac{\partial^2 \sigma_j}{\partial \boldsymbol{\alpha}_j^2},$$
$$= f(y_j;\mu_j,\sigma_j)\left[\frac{2}{\sigma_j^4}\left(3\log(\mu_j\sigma_j^2)-1\right) + \frac{3}{\sigma_j^4} + \frac{2}{\sigma_j^4}\left(3\psi\left(\frac{1}{\sigma_j^2}\right) + \frac{1}{\sigma_j^2}\psi'\left(\frac{1}{\sigma_j^2}\right)\right) + \frac{6\log y_j}{\sigma_j^4} + \frac{6y_j}{\mu_j\sigma_j^4}\right]\left[\frac{\partial \sigma_j}{\partial \boldsymbol{\alpha}_j}\right]^2 +$$
$$+ \frac{\partial f_{\mathrm{GA}}\left(y_j;\ \mu_j,\sigma_j\right)}{\partial \sigma_j}\frac{\partial^2 \sigma_j}{\partial \boldsymbol{\alpha}_j^2}.$$

$$\frac{\partial^2 u_j}{\partial \boldsymbol{\alpha}_j^2} = \left[\frac{\partial \sigma_j}{\partial \boldsymbol{\alpha}_j}\right]^2 \int_0^{y_j} f(t;\mu_j,\sigma_j)\left[\frac{2}{\sigma_j^4}\left(3\log(\mu_j\sigma_j^2)-1\right) + \frac{3}{\sigma_j^4} + \frac{2}{\sigma_j^4}\left(3\psi\left(\frac{1}{\sigma_j^2}\right) + \frac{1}{\sigma_j^2}\psi'\left(\frac{1}{\sigma_j^2}\right)\right) + \frac{6\log t}{\sigma_j^4} + \frac{6t}{\mu_j\sigma_j^4}\right]dt +$$
$$+ \frac{\partial^2 \sigma_j}{\partial \boldsymbol{\alpha}_j^2}\int_0^{y_j}\frac{\partial f_{\mathrm{GA}}\left(t;\ \mu_j,\sigma_j\right)}{\partial \sigma_j}dt.$$

$$\frac{\partial^2 u_j'}{\partial \boldsymbol{\alpha}_j \partial \boldsymbol{\beta}_j} = \frac{\partial}{\partial \boldsymbol{\alpha}_j}\left[\frac{\partial u_j'}{\partial \boldsymbol{\beta}_j}\right] = \frac{\partial}{\partial \boldsymbol{\alpha}_j}\left[f_{\mathrm{GA}}\left(y_j;\ \mu_j,\sigma_j\right)\left[\frac{-1}{\mu_j} + \frac{y_j}{\mu_j^2\sigma_j^2}\right]\frac{\partial \mu_j}{\partial \boldsymbol{\beta}_j}\right],$$
$$= \frac{\partial f_{\mathrm{GA}}\left(y_j;\ \mu_j,\sigma_j\right)}{\partial \boldsymbol{\alpha}_j}\left[\frac{-1}{\mu_j} + \frac{y_j}{\mu_j^2\sigma_j^2}\right]\frac{\partial \mu_j}{\partial \boldsymbol{\beta}_j} + f_{\mathrm{GA}}\left(y_j;\ \mu_j,\sigma_j\right)\frac{\partial}{\partial \sigma_j}\left[\frac{-1}{\mu_j} - \frac{2y_j}{\mu_j^2\sigma_j^3}\right]\frac{\partial \sigma_j}{\partial \boldsymbol{\alpha}_j}\frac{\partial \mu_j}{\partial \boldsymbol{\beta}_j}.$$

$$\frac{\partial^2 u_j}{\partial \boldsymbol{\alpha}_j \partial \boldsymbol{\beta}_j} = \frac{\partial \mu_j}{\partial \boldsymbol{\beta}_j}\int_0^{y_j}\frac{\partial f_{\mathrm{GA}}\left(t;\ \mu_j,\sigma_j\right)}{\partial \boldsymbol{\alpha}_j}\left[\frac{-1}{\mu_j} + \frac{t}{\mu_j^2\sigma_j^2}\right]dt + \frac{\partial \mu_j}{\partial \boldsymbol{\beta}_j}\int_0^{y_j} f_{\mathrm{GA}}\left(t;\ \mu_j,\sigma_j\right)\left[\frac{-1}{\mu_j} - \frac{2t}{\mu_j^2\sigma_j^3}\right]dt.$$

$$
\begin{aligned}
\frac{\partial^2 c(u_1,u_2;\rho)}{\partial \boldsymbol{\tau}^2} &= \frac{\partial}{\partial \boldsymbol{\tau}}\left[\frac{\partial c(u_1,u_2;\rho)}{\partial \boldsymbol{\tau}}\right],\\
&= \frac{1}{(1-\rho^2)^5}\Bigg(-(1-\rho^2)^{\frac{5}{2}}\Bigg(3\rho^2 - \frac{\partial \Phi^{-1}(u_1)}{\partial \rho}2\rho^2 - \Phi^{-1}(u_1)\frac{\partial \Phi^{-1}(u_2)}{\partial \rho}\rho^2 - 2\Phi^{-1}(u_1)\Phi^{-1}(u_2)\rho + \Phi^{-1}(u_2)^2 +\\
&\quad + 2\rho\frac{\partial \Phi^{-1}(u_2)}{\partial \rho}\Phi^{-1}(u_2) + \Phi^{-1}(u_1)^2 + 2\rho\frac{\partial \Phi^{-1}(u_1)}{\partial \rho}\Phi^{-1}(u_1) - 1 - \frac{\partial \Phi^{-1}(u_1)}{\partial \rho} - \Phi^{-1}(u_1)\frac{\partial \Phi^{-1}(u_2)}{\partial \rho} - \Phi^{-1}(u_2)\Bigg)\times\\
&\quad \times \exp\left(\frac{1}{2}\frac{\rho(\Phi^{-1}(u_1)^2 + \Phi^{-1}(u_2)^2 - 2\Phi^{-1}(u_1)\Phi^{-1}(u_2))}{(\rho-1)(\rho+1)}\right) +\\
&\quad + \left(\rho^3 - \Phi^{-1}(u_1)\Phi^{-1}(u_2)\rho^2 + \Phi^{-1}(u_2)^2 + \Phi^{-1}(u_1)^2 - \rho - \Phi^{-1}(u_1)\Phi^{-1}(u_2)\right)\times\\
&\quad \times \exp\left(\frac{1}{2}\frac{\rho(\Phi^{-1}(u_1)^2 + \Phi^{-1}(u_2)^2 - 2\Phi^{-1}(u_1)\Phi^{-1}(u_2))}{(\rho-1)(\rho+1)}\right) +\\
&\quad + \frac{1}{2}\Bigg(\left(\Phi^{-1}(u_1)^2 + \Phi^{-1}(u_2)^2 - 2\Phi^{-1}(u_1)\Phi^{-1}(u_2)\right) + \rho\Bigg(\Phi^{-1}(u_1)^2 + 2\Phi^{-1}(u_1)\frac{\partial \Phi^{-1}(u_1)}{\partial \rho} + \Phi^{-1}(u_2)^2 +\\
&\quad + 2\Phi^{-1}(u_2)\frac{\partial \Phi^{-1}(u_2)}{\partial \rho}\Bigg)\Bigg)\cdot(\rho-1)(\rho+1) - \left(\rho(\Phi^{-1}(u_1)^2 + \Phi^{-1}(u_2)^2 - 2\Phi^{-1}(u_1)\Phi^{-1}(u_2))\right)\times\\
&\quad \times \left((\rho-1)^2 + (\rho+1)^2\right)\Big/((\rho-1)^2(\rho+1)^2) +\\
&\quad - \left(\rho^3 - \Phi^{-1}(u_1)\Phi^{-1}(u_2)\rho^2 + \Phi^{-1}(u_2)^2 + \Phi^{-1}(u_1)^2 - \rho - \Phi^{-1}(u_1)\Phi^{-1}(u_2)\right)\times\\
&\quad \times \exp\left(\frac{1}{2}\frac{\rho(\Phi^{-1}(u_1)^2 + \Phi^{-1}(u_2)^2 - 2\Phi^{-1}(u_1)\Phi^{-1}(u_2))}{(\rho-1)(\rho+1)}\right)(1-\rho^2)^{\frac{3}{2}}5\rho\Bigg).
\end{aligned}
$$

$$
\begin{aligned}
\frac{\partial^2 c(u_1,u_2;\rho)}{\partial \boldsymbol{\beta}_1^2} &= \frac{\partial}{\partial \boldsymbol{\beta}_1}\left[\frac{\partial c(u_1,u_2;\rho)}{\partial \boldsymbol{\beta}_1}\right] = \left[\frac{\partial}{\partial u_1}\left(\frac{\partial c(u_1,u_2;\rho)}{\partial u_1}\frac{\partial u_1}{\partial \boldsymbol{\beta}_1}\right)\right]\frac{\partial u_1}{\partial \boldsymbol{\beta}_1},\\
&= \left[\frac{\partial^2 c(u_1,u_2;\rho)}{\partial u_1^2}\frac{\partial u_1}{\partial \boldsymbol{\beta}_1} + \frac{\partial c(u_1,u_2;\rho)}{\partial u_1}\frac{\partial^2 u_1}{\partial \boldsymbol{\beta}_1^2}\left(\frac{\partial u_1}{\partial \boldsymbol{\beta}_1}\right)^{-1}\right]\frac{\partial u_1}{\partial \boldsymbol{\beta}_1},\\
&= \frac{\partial^2 c(u_1,u_2;\rho)}{\partial u_1^2}\left[\frac{\partial u_1}{\partial \boldsymbol{\beta}_1}\right]^2 + \frac{\partial c(u_1,u_2;\rho)}{\partial u_1}\frac{\partial^2 u_1}{\partial \boldsymbol{\beta}_1^2},\\
&= \Bigg\{\frac{\partial c(u_1,u_2;\rho)}{\partial u_1}\left(\frac{-2\rho^2\Phi^{-1}(u_1)\frac{1}{\phi\{\Phi^{-1}(u_1)\}} - 2\rho\Phi^{-1}(u_2)\frac{1}{\phi\{\Phi^{-1}(u_1)\}}}{2(1-\rho^2)}\right) +\\
&\quad + c(u_1,u_2;\rho)\left(\frac{-2\rho^2\left(\left(\frac{1}{\phi\{\Phi^{-1}(u_1)\}}\right)^2 + \Phi^{-1}(u_1)\frac{\partial^2\Phi^{-1}(u_1)}{\partial u_1^2}\right) - 2\rho\Phi^{-1}(u_2)\frac{\partial^2\Phi^{-1}(u_1)}{\partial u_1^2}}{2(1-\rho^2)}\right)\Bigg\}\left[\frac{\partial u_1}{\partial \boldsymbol{\beta}_1}\right]^2 +\\
&\quad + \frac{\partial c(u_1,u_2;\rho)}{\partial u_1}\frac{\partial^2 u_1}{\partial \boldsymbol{\beta}_1^2}.
\end{aligned}
$$

$$\begin{aligned}
\frac{\partial^2 c(u_1,u_2;\rho)}{\partial \boldsymbol{\beta}_2^2} &= \frac{\partial}{\partial \boldsymbol{\beta}_2}\left[\frac{\partial c(u_1,u_2;\rho)}{\partial \boldsymbol{\beta}_2}\right] = \left[\frac{\partial}{\partial u_2}\left(\frac{\partial c(u_1,u_2;\rho)}{\partial u_2}\frac{\partial u_2}{\partial \boldsymbol{\beta}_2}\right)\right]\frac{\partial u_2}{\partial \boldsymbol{\beta}_2},\\
&= \left[\frac{\partial^2 c(u_1,u_2;\rho)}{\partial u_2^2}\frac{\partial u_2}{\partial \boldsymbol{\beta}_2} + \frac{\partial c(u_1,u_2;\rho)}{\partial u_2}\frac{\partial^2 u_2}{\partial \boldsymbol{\beta}_2^2}\left(\frac{\partial u_2}{\partial \boldsymbol{\beta}_2}\right)^{-1}\right]\frac{\partial u_2}{\partial \boldsymbol{\beta}_2},\\
&= \frac{\partial^2 c(u_1,u_2;\rho)}{\partial u_2^2}\left[\frac{\partial u_2}{\partial \boldsymbol{\beta}_2}\right]^2 + \frac{\partial c(u_1,u_2;\rho)}{\partial u_2}\frac{\partial^2 u_2}{\partial \boldsymbol{\beta}_2^2},\\
&= \left\{\frac{\partial c(u_1,u_2;\rho)}{\partial u_2}\left(\frac{-2\rho^2\Phi^{-1}(u_2)\frac{1}{\phi\{\Phi^{-1}(u_2)\}} - 2\rho\Phi^{-1}(u_1)\frac{1}{\phi\{\Phi^{-1}(u_2)\}}}{2(1-\rho^2)}\right)+\right.\\
&\quad \left. + c(u_1,u_2;\rho)\left(\frac{-2\rho^2\left(\left(\frac{1}{\phi\{\Phi^{-1}(u_2)\}}\right)^2 + \Phi^{-1}(u_2)\frac{\partial^2\Phi^{-1}(u_2)}{\partial u_2^2}\right) - 2\rho\Phi^{-1}(u_1)\frac{\partial^2\Phi^{-1}(u_2)}{\partial u_2^2}}{2(1-\rho^2)}\right)\right\}\left[\frac{\partial u_2}{\partial \boldsymbol{\beta}_2}\right]^2+\\
&\quad + \frac{\partial c(u_1,u_2;\rho)}{\partial u_2}\frac{\partial^2 u_2}{\partial \boldsymbol{\beta}_2^2}.
\end{aligned}$$

$$\begin{aligned}
\frac{\partial^2 c(u_1,u_2;\rho)}{\partial \boldsymbol{\alpha}_j^2} &= \frac{\partial}{\partial \boldsymbol{\alpha}_j}\left[\frac{\partial c(u_1,u_2;\rho)}{\partial \boldsymbol{\alpha}_j}\right] = \left[\frac{\partial}{\partial u_j}\left(\frac{\partial c(u_1,u_2;\rho)}{\partial u_j}\frac{\partial u_j}{\partial \boldsymbol{\alpha}_j}\right)\right]\frac{\partial u_j}{\partial \boldsymbol{\alpha}_j},\\
&= \left[\frac{\partial^2 c(u_1,u_2;\rho)}{\partial u_j^2}\frac{\partial u_j}{\partial \boldsymbol{\alpha}_j} + \frac{\partial c(u_1,u_2;\rho)}{\partial u_j}\frac{\partial^2 u_j}{\partial \boldsymbol{\alpha}_j^2}\left(\frac{\partial u_j}{\partial \boldsymbol{\alpha}_j}\right)^{-1}\right]\frac{\partial u_j}{\partial \boldsymbol{\alpha}_j},\\
&= \frac{\partial^2 c(u_1,u_2;\rho)}{\partial u_j^2}\left[\frac{\partial u_j}{\partial \boldsymbol{\alpha}_j}\right]^2 + \frac{\partial c(u_1,u_2;\rho)}{\partial u_j}\frac{\partial^2 u_j}{\partial \boldsymbol{\alpha}_j^2}.
\end{aligned}$$

$$\begin{aligned}
\frac{\partial^2 c(u_1,u_2;\rho)}{\partial \boldsymbol{\beta}_1\partial \boldsymbol{\beta}_2} &= \frac{\partial}{\partial \boldsymbol{\beta}_1}\left[\frac{\partial c(u_1,u_2;\rho)}{\partial \boldsymbol{\beta}_2}\right] = \left[\frac{\partial}{\partial u_1}\left(\frac{\partial c(u_1,u_2;\rho)}{\partial u_2}\frac{\partial u_2}{\partial \boldsymbol{\beta}_2}\right)\right]\frac{\partial u_1}{\partial \boldsymbol{\beta}_1},\\
&= \left[\frac{\partial^2 c(u_1,u_2;\rho)}{\partial u_1\partial u_2}\frac{\partial u_2}{\partial \boldsymbol{\beta}_2} + \frac{\partial c(u_1,u_2;\rho)}{\partial u_2}\left(\frac{\partial}{\partial u_1}\frac{\partial u_2}{\partial \boldsymbol{\beta}_2}\right)\right]\frac{\partial u_1}{\partial \boldsymbol{\beta}_1},\\
&= \frac{\partial^2 c(u_1,u_2;\rho)}{\partial u_1\partial u_2}\frac{\partial u_2}{\partial \boldsymbol{\beta}_2}\frac{\partial u_1}{\partial \boldsymbol{\beta}_1},\\
&= c(u_1,u_2;\rho)\left[\left(\frac{2\rho^2\Phi^{-1}(u_2)\frac{1}{\phi\{\Phi^{-1}(u_2)\}} - 2\rho\Phi^{-1}(u_1)\frac{1}{\phi\{\Phi^{-1}(u_2)\}}}{2(1-\rho^2)}\right)\left(\frac{\left(2\rho^2\Phi^{-1}(u_1)\frac{1}{\phi\{\Phi^{-1}(u_1)\}} - 2\rho\Phi^{-1}(u_2)\frac{1}{\phi\{\Phi^{-1}(u_1)\}}\right)}{2(1-\rho^2)}\right)+\right.\\
&\quad \left. + \frac{\rho\frac{1}{\phi\{\Phi^{-1}(u_1)\}}\frac{1}{\phi\{\Phi^{-1}(u_2)\}}}{1-\rho^2}\right]\frac{\partial u_2}{\partial \boldsymbol{\beta}_2}\frac{\partial u_1}{\partial \boldsymbol{\beta}_1}.
\end{aligned}$$

$$\begin{aligned}
\frac{\partial^2 c(u_1,u_2;\rho)}{\partial \boldsymbol{\alpha}_1\partial \boldsymbol{\alpha}_2} &= \frac{\partial}{\partial \boldsymbol{\alpha}_1}\left[\frac{\partial c(u_1,u_2;\rho)}{\partial \boldsymbol{\alpha}_2}\right] = \left[\frac{\partial}{\partial u_1}\left(\frac{\partial c(u_1,u_2;\rho)}{\partial u_2}\frac{\partial u_2}{\partial \boldsymbol{\alpha}_2}\right)\right]\frac{\partial u_1}{\partial \boldsymbol{\alpha}_1},\\
&= \left[\frac{\partial^2 c(u_1,u_2;\rho)}{\partial u_1\partial u_2}\frac{\partial u_2}{\partial \boldsymbol{\alpha}_2} + \frac{\partial c(u_1,u_2;\rho)}{\partial u_2}\left(\frac{\partial}{\partial u_1}\frac{\partial u_2}{\partial \boldsymbol{\alpha}_2}\right)\right]\frac{\partial u_1}{\partial \boldsymbol{\alpha}_1},\\
&= \frac{\partial^2 c(u_1,u_2;\rho)}{\partial u_1\partial u_2}\frac{\partial u_2}{\partial \boldsymbol{\alpha}_2}\frac{\partial u_1}{\partial \boldsymbol{\alpha}_1}.
\end{aligned}$$

$$\begin{aligned}
\frac{\partial^2 c(u_1,u_2;\rho)}{\partial \boldsymbol{\alpha}_j\partial \boldsymbol{\beta}_j} &= \frac{\partial}{\partial \boldsymbol{\alpha}_j}\left[\frac{\partial c(u_1,u_2;\rho)}{\partial \boldsymbol{\beta}_j}\right] = \left[\frac{\partial}{\partial u_j}\left(\frac{\partial c(u_1,u_2;\rho)}{\partial u_j}\frac{\partial u_j}{\partial \boldsymbol{\beta}_j}\right)\right]\frac{\partial u_j}{\partial \boldsymbol{\alpha}_j},\\
&= \left[\frac{\partial^2 c(u_1,u_2;\rho)}{\partial u_j^2}\frac{\partial u_j}{\partial \boldsymbol{\beta}_j} + \frac{\partial c(u_1,u_2;\rho)}{\partial u_j}\frac{\partial^2 u_j}{\partial \boldsymbol{\alpha}_j\partial \boldsymbol{\beta}_j}\left(\frac{\partial u_j}{\partial \boldsymbol{\beta}_j}\right)^{-1}\right]\frac{\partial u_j}{\partial \boldsymbol{\alpha}_j},\\
&= \frac{\partial^2 c(u_1,u_2;\rho)}{\partial u_j^2}\frac{\partial u_j}{\partial \boldsymbol{\beta}_j}\frac{\partial u_j}{\partial \boldsymbol{\alpha}_j} + \frac{\partial c(u_1,u_2;\rho)}{\partial u_j}\frac{\partial^2 u_j}{\partial \boldsymbol{\alpha}_j\partial \boldsymbol{\beta}_j}\left(\frac{\partial u_j}{\partial \boldsymbol{\beta}_j}\right)^{-1}\frac{\partial u_j}{\partial \boldsymbol{\alpha}_j}.
\end{aligned}$$

$$
\begin{aligned}
\frac{\partial^2 c(u_1,u_2;\rho)}{\partial \boldsymbol{\alpha}_2 \partial \boldsymbol{\beta}_1} &= \frac{\partial}{\partial \boldsymbol{\alpha}_2}\left[\frac{\partial c(u_1,u_2;\rho)}{\partial \boldsymbol{\beta}_1}\right] = \left[\frac{\partial}{\partial u_2}\left(\frac{\partial c(u_1,u_2;\rho)}{\partial u_1}\frac{\partial u_1}{\partial \boldsymbol{\beta}_1}\right)\right]\frac{\partial u_2}{\partial \boldsymbol{\alpha}_2},\\
&= \left[\frac{\partial^2 c(u_1,u_2;\rho)}{\partial u_2 \partial u_1}\frac{\partial u_1}{\partial \boldsymbol{\beta}_1} + \frac{\partial c(u_1,u_2;\rho)}{\partial u_1}\left(\frac{\partial}{\partial u_2}\frac{\partial u_1}{\partial \boldsymbol{\beta}_1}\right)\right]\frac{\partial u_2}{\partial \boldsymbol{\alpha}_2},\\
&= \frac{\partial^2 c(u_1,u_2;\rho)}{\partial u_2 \partial u_1}\frac{\partial u_1}{\partial \boldsymbol{\beta}_1}\frac{\partial u_2}{\partial \boldsymbol{\alpha}_2}.
\end{aligned}
$$

$$
\begin{aligned}
\frac{\partial^2 c(u_1,u_2;\rho)}{\partial \boldsymbol{\alpha}_1 \partial \boldsymbol{\beta}_2} &= \frac{\partial}{\partial \boldsymbol{\alpha}_1}\left[\frac{\partial c(u_1,u_2;\rho)}{\partial \boldsymbol{\beta}_2}\right] = \left[\frac{\partial}{\partial u_1}\left(\frac{\partial c(u_1,u_2;\rho)}{\partial u_2}\frac{\partial u_2}{\partial \boldsymbol{\beta}_2}\right)\right]\frac{\partial u_1}{\partial \boldsymbol{\alpha}_1},\\
&= \left[\frac{\partial^2 c(u_1,u_2;\rho)}{\partial u_1 \partial u_2}\frac{\partial u_2}{\partial \boldsymbol{\beta}_2} + \frac{\partial c(u_1,u_2;\rho)}{\partial u_2}\left(\frac{\partial}{\partial u_1}\frac{\partial u_2}{\partial \boldsymbol{\beta}_2}\right)\right]\frac{\partial u_1}{\partial \boldsymbol{\alpha}_1},\\
&= \frac{\partial^2 c(u_1,u_2;\rho)}{\partial u_1 \partial u_2}\frac{\partial u_2}{\partial \boldsymbol{\beta}_2}\frac{\partial u_1}{\partial \boldsymbol{\alpha}_1}.
\end{aligned}
$$

$$
\begin{aligned}
\frac{\partial^2 c(u_1,u_2;\rho)}{\partial \boldsymbol{\tau} \partial \boldsymbol{\beta}_j} &= \frac{\partial}{\partial \boldsymbol{\tau}}\left[\frac{\partial c(u_1,u_2;\rho)}{\partial \boldsymbol{\beta}_j}\right] = \left[\frac{\partial}{\partial \rho}\left(\frac{\partial c(u_1,u_2;\rho)}{\partial u_j}\frac{\partial u_j}{\partial \boldsymbol{\beta}_j}\right)\right]\frac{\partial \rho}{\partial \boldsymbol{\tau}},\\
&= \left[\frac{\partial^2 c(u_1,u_2;\rho)}{\partial \rho \partial u_j}\frac{\partial u_j}{\partial \boldsymbol{\beta}_j} + \frac{\partial c(u_1,u_2;\rho)}{\partial u_j}\left(\frac{\partial}{\partial \rho}\frac{\partial u_j}{\partial \boldsymbol{\beta}_j}\right)\right]\frac{\partial \rho}{\partial \boldsymbol{\tau}},\\
&= \frac{\partial^2 c(u_1,u_2;\rho)}{\partial \rho \partial u_j}\frac{\partial u_j}{\partial \boldsymbol{\beta}_j}\frac{\partial \rho}{\partial \boldsymbol{\tau}},\\
&= \left[-\frac{1}{(1-\rho^2)^{\frac{5}{2}}}\left(-\frac{1}{\phi\left\{\Phi^{-1}(u_1)\right\}}\Phi^{-1}(u_2)\rho^2 + 2\rho\Phi^{-1}(u_1)\frac{1}{\phi\left\{\Phi^{-1}(u_1)\right\}} - \frac{1}{\phi\left\{\Phi^{-1}(u_1)\right\}}\right)\times\right.\\
&\quad \times \exp\left\{\frac{1}{2}\frac{\rho\left(\rho\left\{\Phi^{-1}(u_1)\right\}^2 + \rho\left\{\Phi^{-1}(u_2)\right\}^2 - 2\Phi^{-1}(u_1)\Phi^{-1}(u_2)\right)}{(\rho-1)(\rho+1)}\right\} +\\
&\quad \left. + \frac{\partial c(u_1,u_2;\rho)}{\partial \rho}\frac{\rho}{2(\rho-1)(\rho+1)}\left(2\Phi^{-1}(u_1)\frac{1}{\phi\left\{\Phi^{-1}(u_1)\right\}} - 2\frac{1}{\phi\left\{\Phi^{-1}(u_1)\right\}}\Phi^{-1}(u_2)\right)\right]\frac{\partial u_j}{\partial \boldsymbol{\beta}_j}\frac{\partial \rho}{\partial \boldsymbol{\tau}}.
\end{aligned}
$$

$$
\begin{aligned}
\frac{\partial^2 c(u_1,u_2;\rho)}{\partial \boldsymbol{\tau} \partial \boldsymbol{\alpha}_j} &= \frac{\partial}{\partial \boldsymbol{\tau}}\left[\frac{\partial c(u_1,u_2;\rho)}{\partial \boldsymbol{\alpha}_j}\right] = \left[\frac{\partial}{\partial \rho}\left(\frac{\partial c(u_1,u_2;\rho)}{\partial u_j}\frac{\partial u_j}{\partial \boldsymbol{\alpha}_j}\right)\right]\frac{\partial \rho}{\partial \boldsymbol{\tau}},\\
&= \left[\frac{\partial^2 c(u_1,u_2;\rho)}{\partial \rho \partial u_j}\frac{\partial u_j}{\partial \boldsymbol{\alpha}_j} + \frac{\partial c(u_1,u_2;\rho)}{\partial u_j}\left(\frac{\partial}{\partial \rho}\frac{\partial u_j}{\partial \boldsymbol{\alpha}_j}\right)\right]\frac{\partial \rho}{\partial \boldsymbol{\tau}},\\
&= \frac{\partial^2 c(u_1,u_2;\rho)}{\partial \rho \partial u_j}\frac{\partial u_j}{\partial \boldsymbol{\alpha}_j}\frac{\partial \rho}{\partial \boldsymbol{\tau}}.
\end{aligned}
$$

With these second derivatives done, it will be easy to calculate the Hessian of our log-likelihood $\ell$.

##### A.4.1. 2nd derivative with respect to $\boldsymbol{\beta}_1$

$$
\begin{aligned}
\frac{\partial^2 \ell}{\partial \boldsymbol{\beta}_1^2} &= \frac{\partial}{\partial \boldsymbol{\beta}_1}\left(\frac{1}{\mathcal{L}}\frac{\partial \mathcal{L}}{\partial \boldsymbol{\beta}_1}\right) = \left(\frac{\partial}{\partial \boldsymbol{\beta}_1}\frac{1}{\mathcal{L}}\right)\frac{\partial \mathcal{L}}{\partial \boldsymbol{\beta}_1} + \frac{1}{\mathcal{L}}\frac{\partial^2 \mathcal{L}}{\partial \boldsymbol{\beta}_1^2},\\
&= \frac{-1}{\mathcal{L}^2}\left[\frac{\partial \mathcal{L}}{\partial \boldsymbol{\beta}_1}\right]^2 + \frac{1}{\mathcal{L}}\left(\frac{\partial}{\partial \boldsymbol{\beta}_1}\begin{cases} 0 & \text{if } y_1 = 0 \text{ and } y_2 = 0\\ \frac{\partial u_1'}{\partial \boldsymbol{\beta}_1} & \text{if } y_1 > 0 \text{ and } y_2 = 0\\ 0 & \text{if } y_1 = 0 \text{ and } y_2 > 0\\ \frac{\partial c(u_1,u_2;\rho)}{\partial \boldsymbol{\beta}_1}u_1'u_2' + c(u_1,u_2;\rho)\frac{\partial u_1'}{\partial \boldsymbol{\beta}_1}u_2' & \text{if } y_1 > 0 \text{ and } y_2 > 0\end{cases}\right),\\
&= \frac{-1}{\mathcal{L}^2}\left[\frac{\partial \mathcal{L}}{\partial \boldsymbol{\beta}_1}\right]^2 + \frac{1}{\mathcal{L}}\begin{cases} 0 & \text{if } y_1 = 0 \text{ and } y_2 = 0\\ \frac{\partial^2 u_1'}{\partial \boldsymbol{\beta}_1^2} & \text{if } y_1 > 0 \text{ and } y_2 = 0\\ 0 & \text{if } y_1 = 0 \text{ and } y_2 > 0\\ \frac{\partial^2 c(u_1,u_2;\rho)}{\partial \boldsymbol{\beta}_1^2}u_1'u_2' + \frac{\partial c(u_1,u_2;\rho)}{\partial \boldsymbol{\beta}_1}\frac{\partial u_1'}{\partial \boldsymbol{\beta}_1}u_2' + \frac{\partial c(u_1,u_2;\rho)}{\partial \boldsymbol{\beta}_1}\frac{\partial u_1'}{\partial \boldsymbol{\beta}_1}u_2' + c(u_1,u_2;\rho)\frac{\partial^2 u_1'}{\partial \boldsymbol{\beta}_1^2}u_2' & \text{if } y_1 > 0 \text{ and } y_2 > 0.\end{cases}
\end{aligned}
$$

##### A.4.2. 2nd derivative with respect to $\boldsymbol{\beta_2}$

$$\frac{\partial^2 \ell}{\partial \boldsymbol{\beta}_2^2} = \frac{\partial}{\partial \boldsymbol{\beta}_2}\left(\frac{1}{\mathcal{L}}\frac{\partial \mathcal{L}}{\partial \boldsymbol{\beta}_2}\right) = \left(\frac{\partial}{\partial \boldsymbol{\beta}_2}\frac{1}{\mathcal{L}}\right)\frac{\partial \mathcal{L}}{\partial \boldsymbol{\beta}_2} + \frac{1}{\mathcal{L}}\frac{\partial^2 \mathcal{L}}{\partial \boldsymbol{\beta}_2^2},$$

$$= \frac{-1}{\mathcal{L}^2}\left[\frac{\partial \mathcal{L}}{\partial \boldsymbol{\beta}_2}\right]^2 + \frac{1}{\mathcal{L}}\left(\frac{\partial}{\partial \boldsymbol{\beta}_2}\begin{cases} 0 & \text{if } y_1 = 0 \text{ and } y_2 = 0 \\ 0 & \text{if } y_1 > 0 \text{ and } y_2 = 0 \\ \frac{\partial u_2'}{\partial \boldsymbol{\beta}_2} & \text{if } y_1 = 0 \text{ and } y_2 > 0 \\ \frac{\partial c(u_1,u_2;\rho)}{\partial \boldsymbol{\beta}_2}u_1'u_2' + c(u_1,u_2;\rho)u_1'\frac{\partial u_2'}{\partial \boldsymbol{\beta}_2} & \text{if } y_1 > 0 \text{ and } y_2 > 0 \end{cases}\right),$$

$$= \frac{-1}{\mathcal{L}^2}\left[\frac{\partial \mathcal{L}}{\partial \boldsymbol{\beta}_2}\right]^2 + \frac{1}{\mathcal{L}}\begin{cases} 0 & \text{if } y_1 = 0 \text{ and } y_2 = 0 \\ 0 & \text{if } y_1 > 0 \text{ and } y_2 = 0 \\ \frac{\partial^2 u_2'}{\partial \boldsymbol{\beta}_2^2} & \text{if } y_1 = 0 \text{ and } y_2 > 0 \\ \frac{\partial^2 c(u_1,u_2;\rho)}{\partial \boldsymbol{\beta}_2^2}u_1'u_2' + \frac{\partial c(u_1,u_2;\rho)}{\partial \boldsymbol{\beta}_2}u_1'\frac{\partial u_2'}{\partial \boldsymbol{\beta}_2} + \frac{\partial c(u_1,u_2;\rho)}{\partial \boldsymbol{\beta}_2}u_1'\frac{\partial u_2'}{\partial \boldsymbol{\beta}_2} + c(u_1,u_2;\rho)u_1'\frac{\partial^2 u_2'}{\partial \boldsymbol{\beta}_2^2} & \text{if } y_1 > 0 \text{ and } y_2 > 0. \end{cases}$$

##### A.4.3. 2nd derivative with respect to $\boldsymbol{\alpha_1}$

$$\frac{\partial^2 \ell}{\partial \boldsymbol{\alpha}_1^2} = \frac{\partial}{\partial \boldsymbol{\alpha}_1}\left(\frac{1}{\mathcal{L}}\frac{\partial \mathcal{L}}{\partial \boldsymbol{\alpha}_1}\right) = \left(\frac{\partial}{\partial \boldsymbol{\alpha}_1}\frac{1}{\mathcal{L}}\right)\frac{\partial \mathcal{L}}{\partial \boldsymbol{\alpha}_1} + \frac{1}{\mathcal{L}}\frac{\partial^2 \mathcal{L}}{\partial \boldsymbol{\alpha}_1^2},$$

$$= \frac{-1}{\mathcal{L}^2}\left[\frac{\partial \mathcal{L}}{\partial \boldsymbol{\alpha}_1}\right]^2 + \frac{1}{\mathcal{L}}\left(\frac{\partial}{\partial \boldsymbol{\alpha}_1}\begin{cases} 0 & \text{if } y_1 = 0 \text{ and } y_2 = 0 \\ \frac{\partial u_1'}{\partial \boldsymbol{\alpha}_1} & \text{if } y_1 > 0 \text{ and } y_2 = 0 \\ 0 & \text{if } y_1 = 0 \text{ and } y_2 > 0 \\ \frac{\partial c(u_1,u_2;\rho)}{\partial \boldsymbol{\alpha}_1}u_1'u_2' + c(u_1,u_2;\rho)\frac{\partial u_1'}{\partial \boldsymbol{\alpha}_1}u_2' & \text{if } y_1 > 0 \text{ and } y_2 > 0 \end{cases}\right),$$

$$= \frac{-1}{\mathcal{L}^2}\left[\frac{\partial \mathcal{L}}{\partial \boldsymbol{\alpha}_1}\right]^2 + \frac{1}{\mathcal{L}}\begin{cases} 0 & \text{if } y_1 = 0 \text{ and } y_2 = 0 \\ \frac{\partial^2 u_1'}{\partial \boldsymbol{\alpha}_1^2} & \text{if } y_1 > 0 \text{ and } y_2 = 0 \\ 0 & \text{if } y_1 = 0 \text{ and } y_2 > 0 \\ \frac{\partial^2 c(u_1,u_2;\rho)}{\partial \boldsymbol{\alpha}_1^2}u_1'u_2' + \frac{\partial c(u_1,u_2;\rho)}{\partial \boldsymbol{\alpha}_1}\frac{\partial u_1'}{\partial \boldsymbol{\alpha}_1}u_2' + \frac{\partial c(u_1,u_2;\rho)}{\partial \boldsymbol{\alpha}_1}\frac{\partial u_1'}{\partial \boldsymbol{\alpha}_1}u_2' + c(u_1,u_2;\rho)\frac{\partial^2 u_1'}{\partial \boldsymbol{\alpha}_1^2}u_2' & \text{if } y_1 > 0 \text{ and } y_2 > 0. \end{cases}$$

##### A.4.4. 2nd derivative with respect to $\boldsymbol{\alpha_2}$

$$\frac{\partial^2 \ell}{\partial \boldsymbol{\alpha}_2^2} = \frac{\partial}{\partial \boldsymbol{\alpha}_2}\left(\frac{1}{\mathcal{L}}\frac{\partial \mathcal{L}}{\partial \boldsymbol{\alpha}_2}\right) = \left(\frac{\partial}{\partial \boldsymbol{\alpha}_2}\frac{1}{\mathcal{L}}\right)\frac{\partial \mathcal{L}}{\partial \boldsymbol{\alpha}_2} + \frac{1}{\mathcal{L}}\frac{\partial^2 \mathcal{L}}{\partial \boldsymbol{\alpha}_2^2},$$

$$= \frac{-1}{\mathcal{L}^2}\left[\frac{\partial \mathcal{L}}{\partial \boldsymbol{\alpha}_2}\right]^2 + \frac{1}{\mathcal{L}}\left(\frac{\partial}{\partial \boldsymbol{\alpha}_2}\begin{cases} 0 & \text{if } y_1 = 0 \text{ and } y_2 = 0 \\ 0 & \text{if } y_1 > 0 \text{ and } y_2 = 0 \\ \frac{\partial u_2'}{\partial \boldsymbol{\alpha}_2} & \text{if } y_1 = 0 \text{ and } y_2 > 0 \\ \frac{\partial c(u_1,u_2;\rho)}{\partial \boldsymbol{\alpha}_2}u_1'u_2' + c(u_1,u_2;\rho)u_1'\frac{\partial u_2'}{\partial \boldsymbol{\alpha}_2} & \text{if } y_1 > 0 \text{ and } y_2 > 0 \end{cases}\right),$$

$$= \frac{-1}{\mathcal{L}^2}\left[\frac{\partial \mathcal{L}}{\partial \boldsymbol{\alpha}_2}\right]^2 + \frac{1}{\mathcal{L}}\begin{cases} 0 & \text{if } y_1 = 0 \text{ and } y_2 = 0 \\ 0 & \text{if } y_1 > 0 \text{ and } y_2 = 0 \\ \frac{\partial^2 u_2'}{\partial \boldsymbol{\alpha}_2^2} & \text{if } y_1 = 0 \text{ and } y_2 > 0 \\ \frac{\partial^2 c(u_1,u_2;\rho)}{\partial \boldsymbol{\alpha}_2^2}u_1'u_2' + \frac{\partial c(u_1,u_2;\rho)}{\partial \boldsymbol{\alpha}_2}u_1'\frac{\partial u_2'}{\partial \boldsymbol{\alpha}_2} + \frac{\partial c(u_1,u_2;\rho)}{\partial \boldsymbol{\alpha}_2}u_1'\frac{\partial u_2'}{\partial \boldsymbol{\alpha}_2} + c(u_1,u_2;\rho)u_1'\frac{\partial^2 u_2'}{\partial \boldsymbol{\alpha}_2^2} & \text{if } y_1 > 0 \text{ and } y_2 > 0. \end{cases}$$

##### A.4.5. 2nd derivative with respect to $\boldsymbol{\tau}$

$$\frac{\partial^2 \ell}{\partial \boldsymbol{\tau}^2} = \frac{\partial}{\partial \boldsymbol{\tau}}\left(\frac{1}{\mathcal{L}}\frac{\partial \mathcal{L}}{\partial \boldsymbol{\tau}}\right) = \left(\frac{\partial}{\partial \boldsymbol{\tau}}\frac{1}{\mathcal{L}}\right)\frac{\partial \mathcal{L}}{\partial \boldsymbol{\tau}} + \frac{1}{\mathcal{L}}\frac{\partial^2 \mathcal{L}}{\partial \boldsymbol{\tau}^2},$$

$$= \frac{-1}{\mathcal{L}^2}\left[\frac{\partial \mathcal{L}}{\partial \boldsymbol{\tau}}\right]^2 + \frac{1}{\mathcal{L}}\left(\frac{\partial}{\partial \boldsymbol{\tau}}\begin{cases} 0 & \text{if } y_1 = 0 \text{ and } y_2 = 0 \\ 0 & \text{if } y_1 > 0 \text{ and } y_2 = 0 \\ 0 & \text{if } y_1 = 0 \text{ and } y_2 > 0 \\ \frac{\partial c(u_1,u_2;\rho)}{\partial \boldsymbol{\tau}}u_1'u_2' & \text{if } y_1 > 0 \text{ and } y_2 > 0 \end{cases}\right),$$

$$= \frac{-1}{\mathcal{L}^2}\left[\frac{\partial \mathcal{L}}{\partial \boldsymbol{\tau}}\right]^2 + \frac{1}{\mathcal{L}}\begin{cases} 0 & \text{if } y_1 = 0 \text{ and } y_2 = 0 \\ 0 & \text{if } y_1 > 0 \text{ and } y_2 = 0 \\ 0 & \text{if } y_1 = 0 \text{ and } y_2 > 0 \\ \frac{\partial^2 c(u_1,u_2;\rho)}{\partial \boldsymbol{\tau}^2}u_1'u_2' & \text{if } y_1 > 0 \text{ and } y_2 > 0. \end{cases}$$

**A.4.6. Mixed derivative with respect to $\boldsymbol{\beta_1}$ and $\boldsymbol{\beta_2}$**

$$\frac{\partial^2 \ell}{\partial \boldsymbol{\beta}_1 \partial \boldsymbol{\beta}_2} = \frac{\partial}{\partial \boldsymbol{\beta}_1}\left(\frac{1}{\mathcal{L}}\frac{\partial \mathcal{L}}{\partial \boldsymbol{\beta}_2}\right) = \left(\frac{\partial}{\partial \boldsymbol{\beta}_1}\frac{1}{\mathcal{L}}\right)\frac{\partial \mathcal{L}}{\partial \boldsymbol{\beta}_2} + \frac{1}{\mathcal{L}}\frac{\partial^2 \mathcal{L}}{\partial \boldsymbol{\beta}_1 \partial \boldsymbol{\beta}_2},$$

$$= \frac{-1}{\mathcal{L}^2}\frac{\partial \mathcal{L}}{\partial \boldsymbol{\beta}_1}\frac{\partial \mathcal{L}}{\partial \boldsymbol{\beta}_2} + \frac{1}{\mathcal{L}}\left(\frac{\partial}{\partial \boldsymbol{\beta}_1}\begin{cases} 0 & \text{if } y_1 = 0 \text{ and } y_2 = 0 \\ 0 & \text{if } y_1 > 0 \text{ and } y_2 = 0 \\ \frac{\partial u_2'}{\partial \boldsymbol{\beta}_2} & \text{if } y_1 = 0 \text{ and } y_2 > 0 \\ \frac{\partial c(u_1,u_2;\rho)}{\partial \boldsymbol{\beta}_2} u_1' u_2' + c(u_1,u_2;\rho) u_1' \frac{\partial u_2'}{\partial \boldsymbol{\beta}_2} & \text{if } y_1 > 0 \text{ and } y_2 > 0 \end{cases}\right),$$

$$= \frac{-1}{\mathcal{L}^2}\frac{\partial \mathcal{L}}{\partial \boldsymbol{\beta}_1}\frac{\partial \mathcal{L}}{\partial \boldsymbol{\beta}_2} + \frac{1}{\mathcal{L}}\begin{cases} 0 & \text{if } y_1 = 0 \text{ and } y_2 = 0 \\ 0 & \text{if } y_1 > 0 \text{ and } y_2 = 0 \\ 0 & \text{if } y_1 = 0 \text{ and } y_2 > 0 \\ \frac{\partial^2 c(u_1,u_2;\rho)}{\partial \boldsymbol{\beta}_1 \partial \boldsymbol{\beta}_2} u_1' u_2' + \frac{\partial c(u_1,u_2;\rho)}{\partial \boldsymbol{\beta}_2}\frac{\partial u_1'}{\partial \boldsymbol{\beta}_1} u_2' + \frac{\partial c(u_1,u_2;\rho)}{\partial \boldsymbol{\beta}_1} u_1' \frac{\partial u_2'}{\partial \boldsymbol{\beta}_2} + c(u_1,u_2;\rho)\frac{\partial u_1'}{\partial \boldsymbol{\beta}_1}\frac{\partial^2 u_2'}{\partial \boldsymbol{\beta}_2^2} & \text{if } y_1 > 0 \text{ and } y_2 > 0. \end{cases}$$

**A.4.7. Mixed derivative with respect to $\boldsymbol{\alpha_1}$ and $\boldsymbol{\alpha_2}$**

$$\frac{\partial^2 \ell}{\partial \boldsymbol{\alpha}_1 \partial \boldsymbol{\alpha}_2} = \frac{\partial}{\partial \boldsymbol{\alpha}_1}\left(\frac{1}{\mathcal{L}}\frac{\partial \mathcal{L}}{\partial \boldsymbol{\alpha}_2}\right) = \left(\frac{\partial}{\partial \boldsymbol{\alpha}_1}\frac{1}{\mathcal{L}}\right)\frac{\partial \mathcal{L}}{\partial \boldsymbol{\alpha}_2} + \frac{1}{\mathcal{L}}\frac{\partial^2 \mathcal{L}}{\partial \boldsymbol{\alpha}_1 \partial \boldsymbol{\alpha}_2},$$

$$= \frac{-1}{\mathcal{L}^2}\frac{\partial \mathcal{L}}{\partial \boldsymbol{\alpha}_1}\frac{\partial \mathcal{L}}{\partial \boldsymbol{\alpha}_2} + \frac{1}{\mathcal{L}}\left(\frac{\partial}{\partial \boldsymbol{\alpha}_1}\begin{cases} 0 & \text{if } y_1 = 0 \text{ and } y_2 = 0 \\ 0 & \text{if } y_1 > 0 \text{ and } y_2 = 0 \\ \frac{\partial u_2'}{\partial \boldsymbol{\alpha}_2} & \text{if } y_1 = 0 \text{ and } y_2 > 0 \\ \frac{\partial c(u_1,u_2;\rho)}{\partial \boldsymbol{\alpha}_2} u_1' u_2' + c(u_1,u_2;\rho) u_1' \frac{\partial u_2'}{\partial \boldsymbol{\alpha}_2} & \text{if } y_1 > 0 \text{ and } y_2 > 0 \end{cases}\right),$$

$$= \frac{-1}{\mathcal{L}^2}\frac{\partial \mathcal{L}}{\partial \boldsymbol{\alpha}_1}\frac{\partial \mathcal{L}}{\partial \boldsymbol{\alpha}_2} + \frac{1}{\mathcal{L}}\begin{cases} 0 & \text{if } y_1 = 0 \text{ and } y_2 = 0 \\ 0 & \text{if } y_1 > 0 \text{ and } y_2 = 0 \\ 0 & \text{if } y_1 = 0 \text{ and } y_2 > 0 \\ \frac{\partial^2 c(u_1,u_2;\rho)}{\partial \boldsymbol{\alpha}_1 \partial \boldsymbol{\alpha}_2} u_1' u_2' + \frac{\partial c(u_1,u_2;\rho)}{\partial \boldsymbol{\alpha}_2}\frac{\partial u_1'}{\partial \boldsymbol{\alpha}_1} u_2' + \frac{\partial c(u_1,u_2;\rho)}{\partial \boldsymbol{\alpha}_1} u_1' \frac{\partial u_2'}{\partial \boldsymbol{\alpha}_2} + c(u_1,u_2;\rho)\frac{\partial u_1'}{\partial \boldsymbol{\alpha}_1}\frac{\partial^2 u_2'}{\partial \boldsymbol{\alpha}_2^2} & \text{if } y_1 > 0 \text{ and } y_2 > 0. \end{cases}$$

**A.4.8. Mixed derivative with respect to $\boldsymbol{\tau}$ and $\boldsymbol{\beta_1}$**

$$\frac{\partial^2 \ell}{\partial \boldsymbol{\beta}_1 \partial \boldsymbol{\tau}} = \frac{\partial}{\partial \boldsymbol{\beta}_1}\left(\frac{1}{\mathcal{L}}\frac{\partial \mathcal{L}}{\partial \boldsymbol{\tau}}\right) = \left(\frac{\partial}{\partial \boldsymbol{\beta}_1}\frac{1}{\mathcal{L}}\right)\frac{\partial \mathcal{L}}{\partial \boldsymbol{\tau}} + \frac{1}{\mathcal{L}}\frac{\partial^2 \mathcal{L}}{\partial \boldsymbol{\beta}_1 \partial \boldsymbol{\tau}},$$

$$= \frac{-1}{\mathcal{L}^2}\frac{\partial \mathcal{L}}{\partial \boldsymbol{\beta}_1}\frac{\partial \mathcal{L}}{\partial \boldsymbol{\tau}} + \frac{1}{\mathcal{L}}\left(\frac{\partial}{\partial \boldsymbol{\beta}_1}\begin{cases} 0 & \text{if } y_1 = 0 \text{ and } y_2 = 0 \\ 0 & \text{if } y_1 > 0 \text{ and } y_2 = 0 \\ 0 & \text{if } y_1 = 0 \text{ and } y_2 > 0 \\ \frac{\partial c(u_1,u_2;\rho)}{\partial \boldsymbol{\tau}} u_1' u_2' & \text{if } y_1 > 0 \text{ and } y_2 > 0 \end{cases}\right),$$

$$= \frac{-1}{\mathcal{L}^2}\frac{\partial \mathcal{L}}{\partial \boldsymbol{\beta}_1}\frac{\partial \mathcal{L}}{\partial \boldsymbol{\tau}} + \frac{1}{\mathcal{L}}\begin{cases} 0 & \text{if } y_1 = 0 \text{ and } y_2 = 0 \\ 0 & \text{if } y_1 > 0 \text{ and } y_2 = 0 \\ 0 & \text{if } y_1 = 0 \text{ and } y_2 > 0 \\ \frac{\partial^2 c(u_1,u_2;\rho)}{\partial \boldsymbol{\beta}_1 \partial \boldsymbol{\tau}} u_1' u_2' + \frac{\partial c(u_1,u_2;\rho)}{\partial \boldsymbol{\tau}}\frac{\partial u_1'}{\partial \boldsymbol{\beta}_1} u_2' & \text{if } y_1 > 0 \text{ and } y_2 > 0. \end{cases}$$

**A.4.9. Mixed derivative with respect to $\boldsymbol{\tau}$ and $\boldsymbol{\beta_2}$**

$$\frac{\partial^2 \ell}{\partial \boldsymbol{\beta}_2 \partial \boldsymbol{\tau}} = \frac{\partial}{\partial \boldsymbol{\beta}_2}\left(\frac{1}{\mathcal{L}}\frac{\partial \mathcal{L}}{\partial \boldsymbol{\tau}}\right) = \left(\frac{\partial}{\partial \boldsymbol{\beta}_2}\frac{1}{\mathcal{L}}\right)\frac{\partial \mathcal{L}}{\partial \boldsymbol{\tau}} + \frac{1}{\mathcal{L}}\frac{\partial^2 \mathcal{L}}{\partial \boldsymbol{\beta}_2 \partial \boldsymbol{\tau}},$$

$$= \frac{-1}{\mathcal{L}^2}\frac{\partial \mathcal{L}}{\partial \boldsymbol{\beta}_2}\frac{\partial \mathcal{L}}{\partial \boldsymbol{\tau}} + \frac{1}{\mathcal{L}}\left(\frac{\partial}{\partial \boldsymbol{\beta}_2}\begin{cases} 0 & \text{if } y_1 = 0 \text{ and } y_2 = 0 \\ 0 & \text{if } y_1 > 0 \text{ and } y_2 = 0 \\ 0 & \text{if } y_1 = 0 \text{ and } y_2 > 0 \\ \frac{\partial c(u_1,u_2;\rho)}{\partial \boldsymbol{\tau}} u_1' u_2' & \text{if } y_1 > 0 \text{ and } y_2 > 0 \end{cases}\right),$$

$$= \frac{-1}{\mathcal{L}^2}\frac{\partial \mathcal{L}}{\partial \boldsymbol{\beta}_2}\frac{\partial \mathcal{L}}{\partial \boldsymbol{\tau}} + \frac{1}{\mathcal{L}}\begin{cases} 0 & \text{if } y_1 = 0 \text{ and } y_2 = 0 \\ 0 & \text{if } y_1 > 0 \text{ and } y_2 = 0 \\ 0 & \text{if } y_1 = 0 \text{ and } y_2 > 0 \\ \frac{\partial^2 c(u_1,u_2;\rho)}{\partial \boldsymbol{\beta}_2 \partial \boldsymbol{\tau}} u_1' u_2' + \frac{\partial c(u_1,u_2;\rho)}{\partial \boldsymbol{\tau}} u_1' \frac{\partial u_2'}{\partial \boldsymbol{\beta}_2} & \text{if } y_1 > 0 \text{ and } y_2 > 0. \end{cases}$$

**A.4.10. Mixed derivative with respect to $\boldsymbol{\tau}$ and $\boldsymbol{\alpha}_1$**

$$\begin{aligned}
\frac{\partial^2 \ell}{\partial\boldsymbol{\alpha}_1\partial\boldsymbol{\tau}} &= \frac{\partial}{\partial\boldsymbol{\alpha}_1}\left(\frac{1}{\mathcal{L}}\frac{\partial\mathcal{L}}{\partial\boldsymbol{\tau}}\right) = \left(\frac{\partial}{\partial\boldsymbol{\alpha}_1}\frac{1}{\mathcal{L}}\right)\frac{\partial\mathcal{L}}{\partial\boldsymbol{\tau}} + \frac{1}{\mathcal{L}}\frac{\partial^2\mathcal{L}}{\partial\boldsymbol{\alpha}_1\partial\boldsymbol{\tau}},\\
&= \frac{-1}{\mathcal{L}^2}\frac{\partial\mathcal{L}}{\partial\boldsymbol{\alpha}_1}\frac{\partial\mathcal{L}}{\partial\boldsymbol{\tau}} + \frac{1}{\mathcal{L}}\left(\frac{\partial}{\partial\boldsymbol{\alpha}_1}\begin{cases}0 & \text{if } y_1 = 0 \text{ and } y_2 = 0\\ 0 & \text{if } y_1 > 0 \text{ and } y_2 = 0\\ 0 & \text{if } y_1 = 0 \text{ and } y_2 > 0\\ \frac{\partial c(u_1,u_2;\rho)}{\partial\boldsymbol{\tau}}u_1'u_2' & \text{if } y_1 > 0 \text{ and } y_2 > 0\end{cases}\right),\\
&= \frac{-1}{\mathcal{L}^2}\frac{\partial\mathcal{L}}{\partial\boldsymbol{\alpha}_1}\frac{\partial\mathcal{L}}{\partial\boldsymbol{\tau}} + \frac{1}{\mathcal{L}}\begin{cases}0 & \text{if } y_1 = 0 \text{ and } y_2 = 0\\ 0 & \text{if } y_1 > 0 \text{ and } y_2 = 0\\ 0 & \text{if } y_1 = 0 \text{ and } y_2 > 0\\ \frac{\partial^2 c(u_1,u_2;\rho)}{\partial\boldsymbol{\alpha}_1\partial\boldsymbol{\tau}}u_1'u_2' + \frac{\partial c(u_1,u_2;\rho)}{\partial\boldsymbol{\tau}}\frac{\partial u_1'}{\partial\boldsymbol{\alpha}_1}u_2' & \text{if } y_1 > 0 \text{ and } y_2 > 0.\end{cases}
\end{aligned}$$

**A.4.11. Mixed derivative with respect to $\boldsymbol{\tau}$ and $\boldsymbol{\alpha}_2$**

$$\begin{aligned}
\frac{\partial^2 \ell}{\partial\boldsymbol{\alpha}_2\partial\boldsymbol{\tau}} &= \frac{\partial}{\partial\boldsymbol{\alpha}_2}\left(\frac{1}{\mathcal{L}}\frac{\partial\mathcal{L}}{\partial\boldsymbol{\tau}}\right) = \left(\frac{\partial}{\partial\boldsymbol{\alpha}_2}\frac{1}{\mathcal{L}}\right)\frac{\partial\mathcal{L}}{\partial\boldsymbol{\tau}} + \frac{1}{\mathcal{L}}\frac{\partial^2\mathcal{L}}{\partial\boldsymbol{\alpha}_2\partial\boldsymbol{\tau}},\\
&= \frac{-1}{\mathcal{L}^2}\frac{\partial\mathcal{L}}{\partial\boldsymbol{\alpha}_2}\frac{\partial\mathcal{L}}{\partial\boldsymbol{\tau}} + \frac{1}{\mathcal{L}}\left(\frac{\partial}{\partial\boldsymbol{\alpha}_2}\begin{cases}0 & \text{if } y_1 = 0 \text{ and } y_2 = 0\\ 0 & \text{if } y_1 > 0 \text{ and } y_2 = 0\\ 0 & \text{if } y_1 = 0 \text{ and } y_2 > 0\\ \frac{\partial c(u_1,u_2;\rho)}{\partial\boldsymbol{\tau}}u_1'u_2' & \text{if } y_1 > 0 \text{ and } y_2 > 0\end{cases}\right),\\
&= \frac{-1}{\mathcal{L}^2}\frac{\partial\mathcal{L}}{\partial\boldsymbol{\alpha}_2}\frac{\partial\mathcal{L}}{\partial\boldsymbol{\tau}} + \frac{1}{\mathcal{L}}\begin{cases}0 & \text{if } y_1 = 0 \text{ and } y_2 = 0\\ 0 & \text{if } y_1 > 0 \text{ and } y_2 = 0\\ 0 & \text{if } y_1 = 0 \text{ and } y_2 > 0\\ \frac{\partial^2 c(u_1,u_2;\rho)}{\partial\boldsymbol{\alpha}_2\partial\boldsymbol{\tau}}u_1'u_2' + \frac{\partial c(u_1,u_2;\rho)}{\partial\boldsymbol{\tau}}u_1'\frac{\partial u_2'}{\partial\boldsymbol{\alpha}_2} & \text{if } y_1 > 0 \text{ and } y_2 > 0.\end{cases}
\end{aligned}$$

**A.4.12. Mixed derivative with respect to $\boldsymbol{\beta}_1$ and $\boldsymbol{\alpha}_1$**

$$\begin{aligned}
\frac{\partial^2 \ell}{\partial\boldsymbol{\alpha}_1\partial\boldsymbol{\beta}_1} &= \frac{\partial}{\partial\boldsymbol{\alpha}_1}\left(\frac{1}{\mathcal{L}}\frac{\partial\mathcal{L}}{\partial\boldsymbol{\beta}_1}\right) = \left(\frac{\partial}{\partial\boldsymbol{\alpha}_1}\frac{1}{\mathcal{L}}\right)\frac{\partial\mathcal{L}}{\partial\boldsymbol{\beta}_1} + \frac{1}{\mathcal{L}}\frac{\partial^2\mathcal{L}}{\partial\boldsymbol{\alpha}_1\partial\boldsymbol{\beta}_1},\\
&= \frac{-1}{\mathcal{L}^2}\frac{\partial\mathcal{L}}{\partial\boldsymbol{\alpha}_1}\frac{\partial\mathcal{L}}{\partial\boldsymbol{\beta}_1} + \frac{1}{\mathcal{L}}\left(\frac{\partial}{\partial\boldsymbol{\alpha}_1}\begin{cases}0 & \text{if } y_1 = 0 \text{ and } y_2 = 0\\ \frac{\partial u_1'}{\partial\boldsymbol{\beta}_1} & \text{if } y_1 > 0 \text{ and } y_2 = 0\\ 0 & \text{if } y_1 = 0 \text{ and } y_2 > 0\\ \frac{\partial c(u_1,u_2;\rho)}{\partial\boldsymbol{\beta}_1}u_1'u_2' + c(u_1,u_2;\rho)\frac{\partial u_1'}{\partial\boldsymbol{\beta}_1}u_2' & \text{if } y_1 > 0 \text{ and } y_2 > 0\end{cases}\right),\\
&= \frac{-1}{\mathcal{L}^2}\frac{\partial\mathcal{L}}{\partial\boldsymbol{\alpha}_1}\frac{\partial\mathcal{L}}{\partial\boldsymbol{\beta}_1} + \frac{1}{\mathcal{L}}\begin{cases}0 & \text{if } y_1 = 0 \text{ and } y_2 = 0\\ \frac{\partial^2 u_1'}{\partial\boldsymbol{\alpha}_1\partial\boldsymbol{\beta}_1} & \text{if } y_1 > 0 \text{ and } y_2 = 0\\ 0 & \text{if } y_1 = 0 \text{ and } y_2 > 0\\ \frac{\partial^2 c(u_1,u_2;\rho)}{\partial\boldsymbol{\alpha}_1\partial\boldsymbol{\beta}_1}u_1'u_2' + \frac{\partial c(u_1,u_2;\rho)}{\partial\boldsymbol{\beta}_1}\frac{\partial u_1'}{\partial\boldsymbol{\alpha}_1}u_2' + \frac{\partial c(u_1,u_2;\rho)}{\partial\boldsymbol{\alpha}_1}\frac{\partial u_1'}{\partial\boldsymbol{\beta}_1}u_2' + c(u_1,u_2;\rho)\frac{\partial^2 u_1'}{\partial\boldsymbol{\alpha}_1\partial\boldsymbol{\beta}_1}u_2' & \text{if } y_1 > 0 \text{ and } y_2 > 0.\end{cases}
\end{aligned}$$

**A.4.13. Mixed derivative with respect to $\boldsymbol{\beta}_2$ and $\boldsymbol{\alpha}_2$**

$$\begin{aligned}
\frac{\partial^2 \ell}{\partial\boldsymbol{\alpha}_2\partial\boldsymbol{\beta}_2} &= \frac{\partial}{\partial\boldsymbol{\alpha}_2}\left(\frac{1}{\mathcal{L}}\frac{\partial\mathcal{L}}{\partial\boldsymbol{\beta}_2}\right) = \left(\frac{\partial}{\partial\boldsymbol{\alpha}_2}\frac{1}{\mathcal{L}}\right)\frac{\partial\mathcal{L}}{\partial\boldsymbol{\beta}_2} + \frac{1}{\mathcal{L}}\frac{\partial^2\mathcal{L}}{\partial\boldsymbol{\alpha}_2\partial\boldsymbol{\beta}_2},\\
&= \frac{-1}{\mathcal{L}^2}\frac{\partial\mathcal{L}}{\partial\boldsymbol{\alpha}_2}\frac{\partial\mathcal{L}}{\partial\boldsymbol{\beta}_2} + \frac{1}{\mathcal{L}}\left(\frac{\partial}{\partial\boldsymbol{\alpha}_2}\begin{cases}0 & \text{if } y_1 = 0 \text{ and } y_2 = 0\\ 0 & \text{if } y_1 > 0 \text{ and } y_2 = 0\\ \frac{\partial u_2'}{\partial\boldsymbol{\beta}_2} & \text{if } y_1 = 0 \text{ and } y_2 > 0\\ \frac{\partial c(u_1,u_2;\rho)}{\partial\boldsymbol{\beta}_2}u_1'u_2' + c(u_1,u_2;\rho)u_1'\frac{\partial u_2'}{\partial\boldsymbol{\beta}_2} & \text{if } y_1 > 0 \text{ and } y_2 > 0\end{cases}\right),\\
&= \frac{-1}{\mathcal{L}^2}\frac{\partial\mathcal{L}}{\partial\boldsymbol{\alpha}_2}\frac{\partial\mathcal{L}}{\partial\boldsymbol{\beta}_2} + \frac{1}{\mathcal{L}}\begin{cases}0 & \text{if } y_1 = 0 \text{ and } y_2 = 0\\ 0 & \text{if } y_1 > 0 \text{ and } y_2 = 0\\ \frac{\partial^2 u_2'}{\partial\boldsymbol{\alpha}_2\partial\boldsymbol{\beta}_2} & \text{if } y_1 = 0 \text{ and } y_2 > 0\\ \frac{\partial^2 c(u_1,u_2;\rho)}{\partial\boldsymbol{\alpha}_2\partial\boldsymbol{\beta}_2}u_1'u_2' + \frac{\partial c(u_1,u_2;\rho)}{\partial\boldsymbol{\beta}_2}u_1'\frac{\partial u_2'}{\partial\boldsymbol{\alpha}_2} + \frac{\partial c(u_1,u_2;\rho)}{\partial\boldsymbol{\alpha}_2}u_1'\frac{\partial u_2'}{\partial\boldsymbol{\beta}_2} + c(u_1,u_2;\rho)u_1'\frac{\partial^2 u_2'}{\partial\boldsymbol{\alpha}_2\partial\boldsymbol{\beta}_2} & \text{if } y_1 > 0 \text{ and } y_2 > 0.\end{cases}
\end{aligned}$$

###### A.4.14. Mixed derivative with respect to $\boldsymbol{\beta_2}$ and $\boldsymbol{\alpha_1}$

$$\frac{\partial^2 \ell}{\partial \boldsymbol{\alpha}_1 \partial \boldsymbol{\beta}_2} = \frac{\partial}{\partial \boldsymbol{\alpha}_1}\left(\frac{1}{\mathcal{L}}\frac{\partial \mathcal{L}}{\partial \boldsymbol{\beta}_2}\right) = \left(\frac{\partial}{\partial \boldsymbol{\alpha}_1}\frac{1}{\mathcal{L}}\right)\frac{\partial \mathcal{L}}{\partial \boldsymbol{\beta}_2} + \frac{1}{\mathcal{L}}\frac{\partial^2 \mathcal{L}}{\partial \boldsymbol{\alpha}_1 \partial \boldsymbol{\beta}_2},$$

$$= \frac{-1}{\mathcal{L}^2}\frac{\partial \mathcal{L}}{\partial \boldsymbol{\alpha}_1}\frac{\partial \mathcal{L}}{\partial \boldsymbol{\beta}_2} + \frac{1}{\mathcal{L}}\left(\frac{\partial}{\partial \boldsymbol{\alpha}_1}\begin{cases} 0 & \text{if } y_1 = 0 \text{ and } y_2 = 0 \\ 0 & \text{if } y_1 > 0 \text{ and } y_2 = 0 \\ \frac{\partial u_2'}{\partial \boldsymbol{\beta}_2} & \text{if } y_1 = 0 \text{ and } y_2 > 0 \\ \frac{\partial c(u_1,u_2;\rho)}{\partial \boldsymbol{\beta}_2} u_1' u_2' + c(u_1,u_2;\rho) u_1' \frac{\partial u_2'}{\partial \boldsymbol{\beta}_2} & \text{if } y_1 > 0 \text{ and } y_2 > 0 \end{cases}\right),$$

$$= \frac{-1}{\mathcal{L}^2}\frac{\partial \mathcal{L}}{\partial \boldsymbol{\alpha}_1}\frac{\partial \mathcal{L}}{\partial \boldsymbol{\beta}_2} + \frac{1}{\mathcal{L}}\begin{cases} 0 & \text{if } y_1 = 0 \text{ and } y_2 = 0 \\ 0 & \text{if } y_1 > 0 \text{ and } y_2 = 0 \\ \frac{\partial u_2'}{\partial \boldsymbol{\beta}_2} & \text{if } y_1 = 0 \text{ and } y_2 > 0 \\ \frac{\partial^2 c(u_1,u_2;\rho)}{\partial \boldsymbol{\alpha}_1 \partial \boldsymbol{\beta}_2} u_1' u_2' + \frac{\partial c(u_1,u_2;\rho)}{\partial \boldsymbol{\beta}_2}\frac{\partial u_1'}{\partial \boldsymbol{\alpha}_1} u_2' + \frac{\partial c(u_1,u_2;\rho)}{\partial \boldsymbol{\alpha}_1} u_1' \frac{\partial u_2'}{\partial \boldsymbol{\beta}_2} + c(u_1,u_2;\rho)\frac{\partial u_1'}{\partial \boldsymbol{\alpha}_1}\frac{\partial u_2'}{\partial \boldsymbol{\beta}_2} & \text{if } y_1 > 0 \text{ and } y_2 > 0. \end{cases}$$

###### A.4.15. Mixed derivative with respect to $\boldsymbol{\beta_1}$ and $\boldsymbol{\alpha_2}$

$$\frac{\partial^2 \ell}{\partial \boldsymbol{\alpha}_2 \partial \boldsymbol{\beta}_1} = \frac{\partial}{\partial \boldsymbol{\alpha}_2}\left(\frac{1}{\mathcal{L}}\frac{\partial \mathcal{L}}{\partial \boldsymbol{\beta}_1}\right) = \left(\frac{\partial}{\partial \boldsymbol{\alpha}_2}\frac{1}{\mathcal{L}}\right)\frac{\partial \mathcal{L}}{\partial \boldsymbol{\beta}_1} + \frac{1}{\mathcal{L}}\frac{\partial^2 \mathcal{L}}{\partial \boldsymbol{\alpha}_2 \partial \boldsymbol{\beta}_1},$$

$$= \frac{-1}{\mathcal{L}^2}\frac{\partial \mathcal{L}}{\partial \boldsymbol{\alpha}_2}\frac{\partial \mathcal{L}}{\partial \boldsymbol{\beta}_1} + \frac{1}{\mathcal{L}}\left(\frac{\partial}{\partial \boldsymbol{\alpha}_2}\begin{cases} 0 & \text{if } y_1 = 0 \text{ and } y_2 = 0 \\ \frac{\partial u_1'}{\partial \boldsymbol{\beta}_1} & \text{if } y_1 > 0 \text{ and } y_2 = 0 \\ 0 & \text{if } y_1 = 0 \text{ and } y_2 > 0 \\ \frac{\partial c(u_1,u_2;\rho)}{\partial \boldsymbol{\beta}_1} u_1' u_2' + c(u_1,u_2;\rho)\frac{\partial u_1'}{\partial \boldsymbol{\beta}_1} u_2' & \text{if } y_1 > 0 \text{ and } y_2 > 0 \end{cases}\right),$$

$$= \frac{-1}{\mathcal{L}^2}\frac{\partial \mathcal{L}}{\partial \boldsymbol{\alpha}_2}\frac{\partial \mathcal{L}}{\partial \boldsymbol{\beta}_1} + \frac{1}{\mathcal{L}}\begin{cases} 0 & \text{if } y_1 = 0 \text{ and } y_2 = 0 \\ 0 & \text{if } y_1 > 0 \text{ and } y_2 = 0 \\ 0 & \text{if } y_1 = 0 \text{ and } y_2 > 0 \\ \frac{\partial^2 c(u_1,u_2;\rho)}{\partial \boldsymbol{\alpha}_2 \partial \boldsymbol{\beta}_1} u_1' u_2' + \frac{\partial c(u_1,u_2;\rho)}{\partial \boldsymbol{\beta}_1} u_1' \frac{\partial u_2'}{\partial \boldsymbol{\alpha}_2} + \frac{\partial c(u_1,u_2;\rho)}{\partial \boldsymbol{\alpha}_2}\frac{\partial u_1'}{\partial \boldsymbol{\beta}_1} u_2' + c(u_1,u_2;\rho)\frac{\partial u_1'}{\partial \boldsymbol{\beta}_1}\frac{\partial u_2'}{\partial \boldsymbol{\alpha}_2} & \text{if } y_1 > 0 \text{ and } y_2 > 0. \end{cases}$$

##### A.5. Parameter estimation

We use the extended trust region algorithm (Radice et al. (2016) and Marra et al. (2017)) to maximize the penalized model's log-likelihood function and get the estimations of additive predictors $\boldsymbol{\eta}$ and smoothing parameters $\boldsymbol{\lambda}$. The process for estimating the model is,

1. Initialize the smoothing parameters $\lambda$s.
2. At iteration $a$, hold $\boldsymbol{\lambda}^{[a]}$ as constants at their values from the last iteration or step (1) if it's the first iteration. Trust region algorithm seeks to minimize $\ell_p$ in Equation 9, it's equivalent to find the step $\boldsymbol{e}$ that minimizes a simpler quadratic model $\tilde{\ell_p}$ defined as,

$$\begin{aligned} \tilde{\ell_p}(\boldsymbol{\delta}^{[a]}) &= -\{\ell_p(\boldsymbol{\delta}^{[a]}) + \boldsymbol{e}^\top \boldsymbol{g}_p(\boldsymbol{\delta}^{[a]}) + \frac{1}{2}\boldsymbol{e}^\top \boldsymbol{H}_p(\boldsymbol{\delta}^{[a]})\boldsymbol{e}\} \quad \text{such that } \|\boldsymbol{e}\| \leq r^{[a]}, \\ \boldsymbol{\delta}^{[a+1]} &= \boldsymbol{\delta}^{[a]} + \operatorname*{argmin}_{\mathbf{e}} \tilde{\ell_p}(\boldsymbol{\delta}^{[a]}), \end{aligned} \tag{A.2}$$

where $\boldsymbol{g}_p(\boldsymbol{\delta}^{[a]}) = \boldsymbol{g}(\boldsymbol{\delta}^{[a]}) - \boldsymbol{S}_{\boldsymbol{\lambda}^{[a]}}\boldsymbol{\delta}^{[a]}$ and $\boldsymbol{H}_p(\boldsymbol{\delta}^{[a]}) = \boldsymbol{H}(\boldsymbol{\delta}^{[a]}) - \boldsymbol{S}_{\boldsymbol{\lambda}^{[a]}}$ are the penalized gradient and Hessian of Equation 9; $\boldsymbol{g}(\boldsymbol{\delta}^{[a]}) = (\partial\ell(\boldsymbol{\delta})/\partial\boldsymbol{\beta}_{\theta_1}|_{\boldsymbol{\beta}_{\theta_1}=\boldsymbol{\beta}_{\theta_1}^{[a]}}, ..., \partial\ell(\boldsymbol{\delta})/\partial\boldsymbol{\beta}_{\theta_M}|_{\boldsymbol{\beta}_{\theta_M}=\boldsymbol{\beta}_{\theta_M}^{[a]}})^\top$, and $\boldsymbol{H}(\boldsymbol{\delta}^{[a]})$ has elements $\boldsymbol{H}(\boldsymbol{\delta}^{[a]})_{s,t} = \partial^2\ell(\boldsymbol{\delta})/\partial\boldsymbol{\beta}_{\theta_s}\partial\boldsymbol{\beta}_{\theta_t}^\top|_{\boldsymbol{\beta}_s=\boldsymbol{\beta}_s^{[a]},\boldsymbol{\beta}_t=\boldsymbol{\beta}_t^{[a]}}$, where $s,t = 1,,,M$. $\|\cdot\|$ is the Euclidean norm. $\boldsymbol{e}$ is the step to update the parameters, how far $\boldsymbol{e}$ can move within each iteration is constrained by a trust region of radius $r^{[a]}$.
3. Holding $\boldsymbol{\delta}^{[a+1]}$ constant at the values found in step (2), find the value of $\boldsymbol{\lambda}^{[a+1]}$ that minimizes the special equation involving the hessian and gradient derived by Marra et al. (2017),

$$\boldsymbol{\lambda}^{[a+1]} = \operatorname*{argmin}_{\boldsymbol{\lambda}}\|\boldsymbol{M}^{[a+1]} - \boldsymbol{A}^{[a+1]}\boldsymbol{M}^{[a+1]}\|^2 - \check{n} + 2\text{tr}(\boldsymbol{A}^{[a+1]}), \tag{A.3}$$

where $\boldsymbol{M}^{[a+1]} = \sqrt{-\boldsymbol{H}(\boldsymbol{\delta}^{[a+1]})}\boldsymbol{\delta}^{[a+1]} + \sqrt{-\boldsymbol{H}(\boldsymbol{\delta}^{[a+1]})}^{-1}\boldsymbol{g}(\boldsymbol{\delta}^{[a+1]})$, $\boldsymbol{A}^{[a+1]} = \sqrt{-\boldsymbol{H}(\boldsymbol{\delta}^{[a+1]})}(-\boldsymbol{H}(\boldsymbol{\delta}^{[a+1]}) + \boldsymbol{S})^{-1}\sqrt{-\boldsymbol{H}(\boldsymbol{\delta}^{[a+1]})}$. $\text{tr}(\boldsymbol{A}^{[a+1]})$ is the number of effective degrees of freedom (edf) of the penalized model; we have two marginal parameters and one copula distributional parameter, so $\check{n} = 5n$. Equation A.3 is solved by using method from Wood (2004).
4. Repeat steps (2) and (3) until $\ell_p$ converges.

The spline we used in this framework is the thin plate regression spline. The penalized cubic regression splines, cyclic penalized cubic regression splines, etc. can also be used. To get the estimated parameters $\hat{\theta}_m$, plug in the estimated additive predictors $\eta\hat{_{\theta_m}}$ into the inverse link functions, $\hat{\theta}_m = g_m^{-1}(\hat{\eta}_{\theta_m})$.

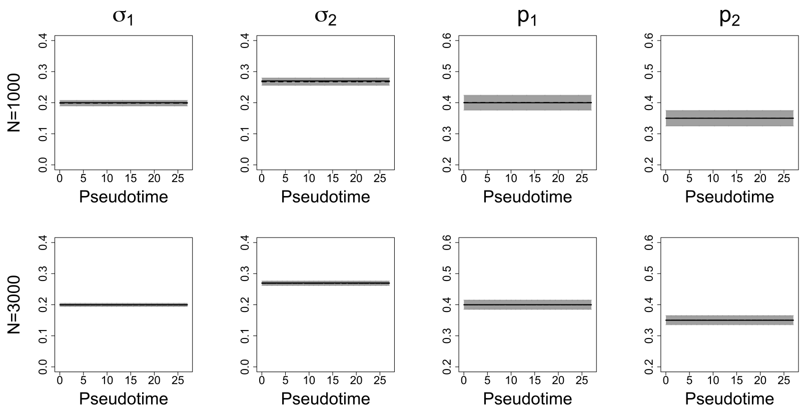


**Fig. 1**: $\sigma$ and $p$ plots in Scenario II simulation: The light gray color is for the control group, and the dark gray color is for the mutant group. The solid lines are the true smooth functions, and the dashed lines are the mean estimates for 1,000 iterations. The shaded areas are point-wise ranges from 5% to 95% quantile. The numbers of observations are 1,000 and 3,000 for each row.

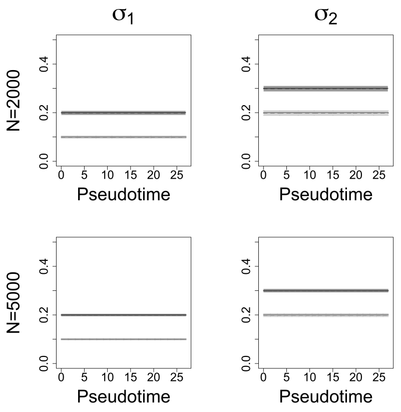


**Fig. 2**: $\sigma$ plots in Scenario III simulation: The light gray color is for the control group, and the dark gray color is for the mutant group. The solid lines are the true smooth functions, and the dashed lines are the mean estimates for 1,000 iterations. The shaded areas are point-wise ranges from 5% to 95% quantile. The numbers of observations are 2,000 and 5,000 for each row.

### B. Appendix

#### B.1. Additional Simulation Plots

This section contains the plots of standard deviation $\sigma$ and zero-inflation rates $p$ in Scenario II Simulation shown in Figure 1. The plots of standard deviation $\sigma$ in Scenario III Simulation are shown in Figure 2.

#### B.2. Significant Gene Pair Plots

This section contains the remaining significant 37 gene pairs correlation plots shown in Figure 3.

#### B.3. QQ Plots

This section shows the QQ plots of 4 gene pairs model fitting in Section 3.

#### B.4. Zero-inflation Plots

This section shows the more Zero-inflation Plots in Figure 5.

#### B.5. Data Analysis Gene List

Table 1 provides the two gene lists we used in Experiment Data Analysis.

### References

G. Marra, R. Radice, T. Bärnighausen, S. N. Wood, and M. E. McGovern. A simultaneous equation approach to estimating HIV prevalence with nonignorable missing responses. *Journal of the American Statistical Association*, 112(518):484–496, 2017.

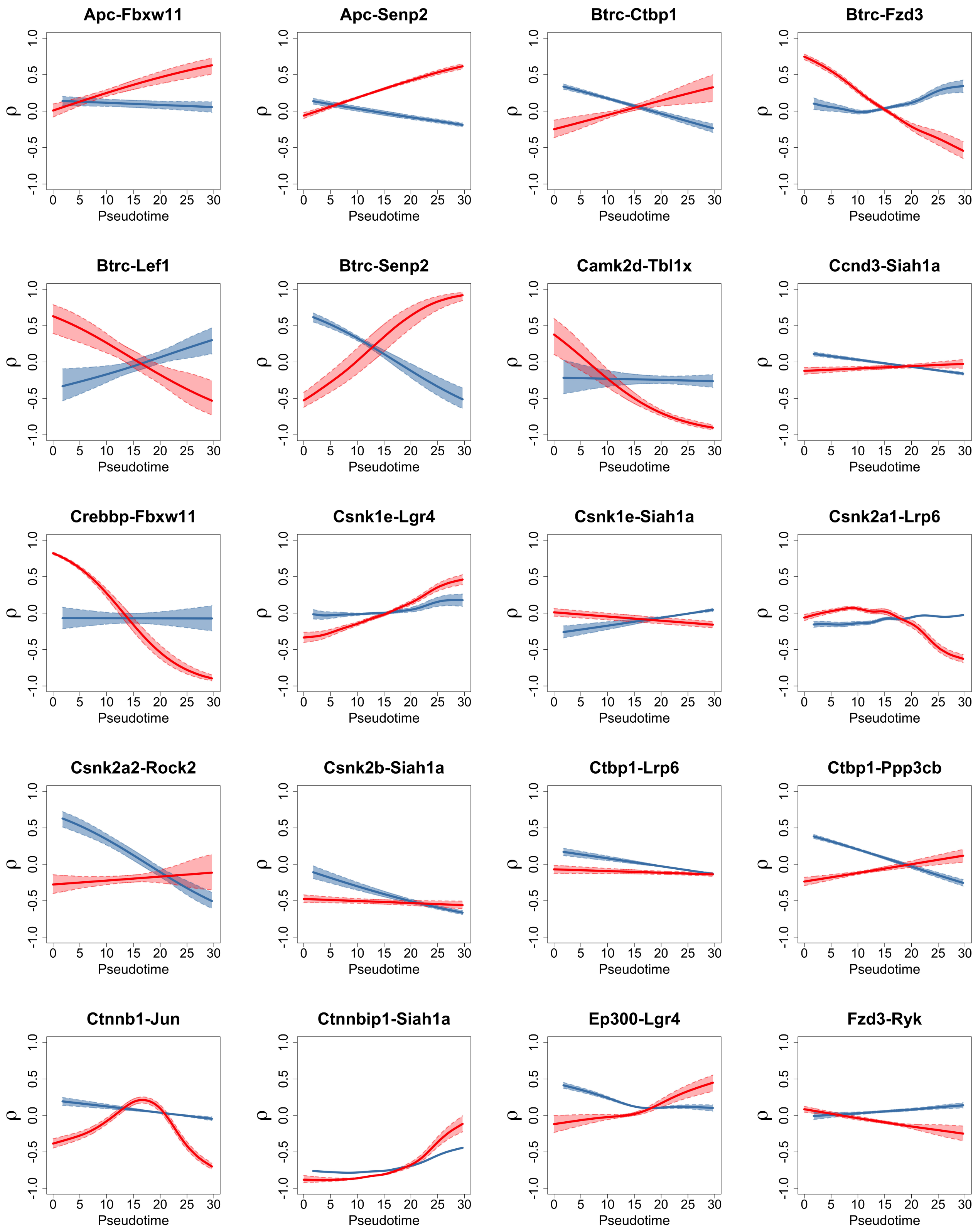


**Fig. 3**: Significant gene pairs with adjusted p-values less than 0.05. The blue solid curve is the fitted line for the control group, and the blue dashed line is 95% CI of the fit. The red solid curve is the fitted line for the mutant group, and the red dashed line is 95% CI of the fit.

R. Radice, G. Marra, and M. Wojtyś. Copula regression spline models for binary outcomes. *Statistics and Computing*, 26(5):981–995, 2016.

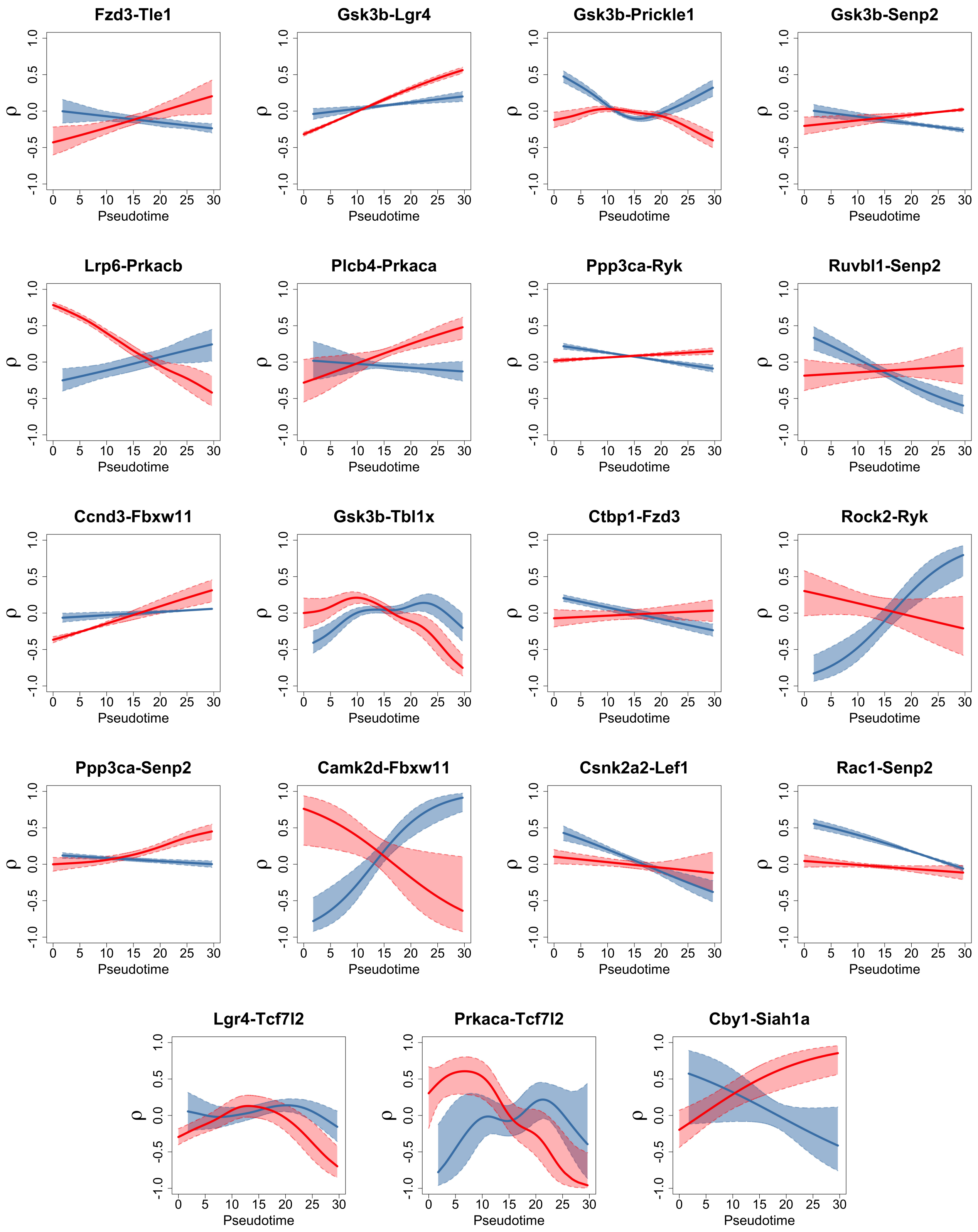


**Fig. 3**: Figure 3 (continued)

R. A. Rigby, M. D. Stasinopoulos, G. Z. Heller, and F. D. Bastiani. *Distributions for modeling location, scale, and shape: Using GAMLSS in R.* Chapman and Hall/CRC, 2019.

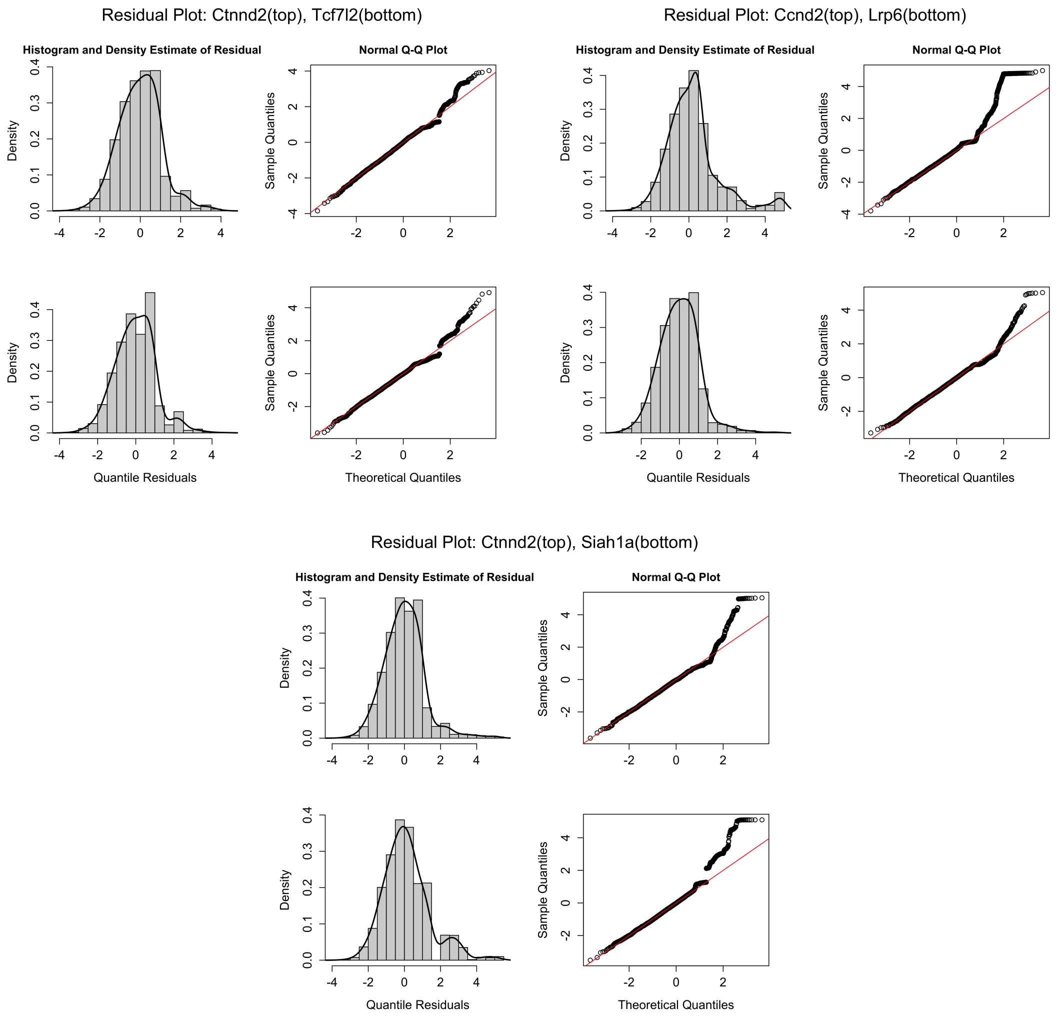


**Fig. 4**: Residual plots of normalized quantile for 3 example gene pairs. The red solid line is $y = x$.

**Table 1.** WNT signaling pathway and TF gene lists used in Experiment Data Analysis.

| | Gene |
|---|---|
| WNT pathway | *Apc, Btrc, Cacybp, Camk2d, Cby1, Ccnd2, Ccnd3, Chd8, Crebbp, Csnk1a1, Csnk1e Csnk2a1, Csnk2a2, Csnk2b, Ctbp1, Ctnnb1, Ctnnbip1, Ctnnd2, Cul1, Cxxc4, Ep300 Fbxw11, Fzd2, Fzd3, Gsk3b, Jun, Lef1, Lgr4, Lrp6, Nlk, Plcb4, Ppp3ca, Ppp3cb Ppp3r1, Prickle1, Prkaca, Prkacb, Rac1, Rbx1, Rhoa, Rock2, Ruvbl1, Ryk, Senp2 Senp2, Siah1a, Smad4, Tbl1x, Tcf7l2, Tle1, Tle4* |
| TF | *Aldh1a2, Axin2, Bmp2, Bmp4, Cga, Egr1, Fgf1, Fgf10, Fgf8,Gata2, Gli1, Gli2, Gli3 Hesx1, Isl1, Lef1, Lhx2, Lhx3, Lhx4, Mki67, Msx1, Neurod1, Neurod4, Nkx2-1, Nkx2-4 Notch2, Otx2, Pax6, Pitx1, Pitx2, Pomc, Pou1f1, Prl, Prop1, Robo2, Rxrg, Sf1, Shh Shh, Six3, Sox1, Sox2, Sox3, Tbx19, Tcf7l2, Tef, Tshb, Wnt11, Wnt3, Wnt4, Wnt5a Wnt5b, Wnt6, Wnt7b, Wnt8b, Wnt9a* |

S. N. Wood. Stable and efficient multiple smoothing parameter estimation for generalized additive models. *Journal of the American Statistical Association*, 99(467):673–686, 2004.

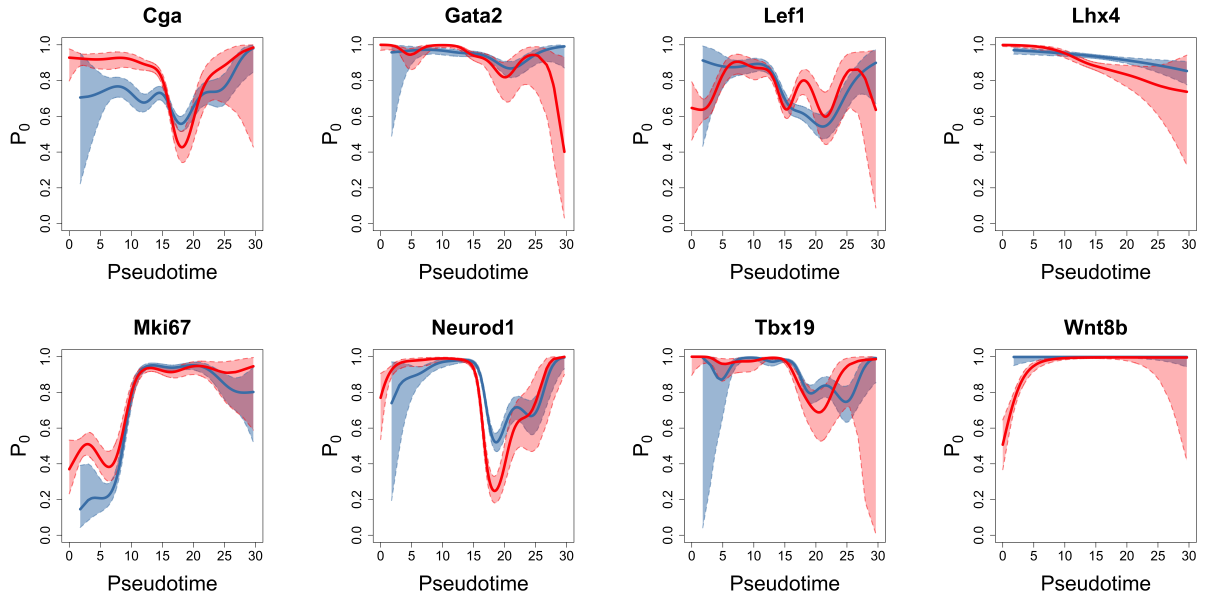


**Fig. 5**: More zero-inflation plots.
